# A novel NLP-based method and algorithm to discover RNA-binding protein (RBP) motifs, contexts, binding preferences, and interactions

**DOI:** 10.1101/2025.01.20.631609

**Authors:** Shaimae I. Elhajjajy, Zhiping Weng

## Abstract

RNA-binding proteins (RBPs) are essential modulators in the regulation of mRNA processing. The binding patterns, interactions, and functions of most RBPs are not well-characterized. Previous studies have shown that motif context is an important contributor to RBP binding specificity, but its precise role remains unclear. Despite recent computational advances to predict RBP binding, existing methods are challenging to interpret and largely lack a categorical focus on RBP motif contexts and RBP-RBP interactions. There remains a need for interpretable predictive models to disambiguate the contextual determinants of RBP binding specificity *in vivo*. Here, we present a novel and comprehensive pipeline to address these knowledge gaps. We devise a Natural Language Processing-based method to deconstruct sequences into entities comprising a target *k*-mer and its flanking regions, then use this representation to formulate RBP binding prediction as a weakly supervised Multiple Instance Learning problem. To interpret our predictions, we introduce a deterministic motif discovery algorithm to leverage our data structure, recapitulating the established motifs of numerous RBPs as validation. Importantly, we characterize the binding motifs and binding contexts for 71 RBPs in HepG2 and 74 RBPs in K562, with many of them being novel. Finally, through feature integration, transitive inference, and a new cross-prediction approach, we propose novel cooperative and competitive RBP-RBP interaction partners and hypothesize their potential regulatory functions. In summary, we present a complete framework for investigating the contextual determinants of specific RBP binding, and we demonstrate the significance of our findings in delineating RBP binding patterns, interactions, and functions.

## INTRODUCTION

RNA-binding proteins (RBPs) extensively govern the biogenesis and maturation of nascent messenger RNAs (mRNAs) in co– and post-transcriptional manners (Gerstberger et al. 2014; Bentley 2014). The human genome encodes approximately 1,500 RBPs, each of which has diverse roles in regulating nearly every step of mRNA processing, including transcription, canonical and alternative splicing, 3’ end cleavage and polyadenylation, stability, localization and transport, degradation, and translation (Gerstberger et al. 2014). RBPs accomplish these tasks by binding their mRNA targets to form messenger ribonucleoprotein (mRNP) complexes whose constituents are dynamically remodeled based on the functional requirements at each stage of mRNA processing (Gerstberger et al. 2014; Müller-McNicoll and Neugebauer 2013; Gehring et al. 2017). Thus, the activities and functions of RBPs are major determinants of the homeostatic expression and heterogeneity of the steady-state transcriptome. The dysregulation of RBPs can manifest into a variety of Mendelian (Castello et al. 2013) and somatic genetic diseases (Gebauer et al. 2021), including, most notably, neurological disorders (Nussbacher et al. 2015; Conlon and Manley 2017) and cancers (Pereira et al. 2017). Because the majority of RBPs catalogued to-date are not well-characterized, an improved understanding of RBP binding patterns, activities, functions, and interactions is needed to delineate their constitutive roles in mediating mRNA metabolism and to discern their causal roles in disease.

RBPs interact with their target mRNAs through RNA-binding domains (RBDs), which confer upon the RBPs their sequence and structural binding preferences (Gerstberger et al. 2014; Müller-McNicoll and Neugebauer 2013; Gehring et al. 2017). The complex and enigmatic binding patterns of RBPs can be attributed in part to their structural composition: they can possess single or multiple RBDs that belong to the same or different domain class (Gerstberger et al. 2014), and these RBDs may be either well-characterized or non-canonical (Ray et al. 2023), together enabling a sizable potential interactome. Due to their short motif length (between 3 to 8 nucleotides) and low nucleotide complexity (often containing only a subset of the 4 canonical nucleotides), RBPs with structurally diverse RBDs can recognize highly similar motifs, but they still assert a remarkable degree of transcriptomic binding specificity (Dominguez et al. 2018). Based on large-scale *in vitro* experiments, previous studies have posited that the flanking sequences of RBP binding sites may play a critical role in enabling RBP binding specificity by fostering conformations conducive to stable binding; however, these findings were based on short, synthetic oligonucleotide sequences in isolated, exogenous conditions lacking a standard cellular environment (Dominguez et al. 2018). Further investigation is necessary to corroborate the contribution of these flanking regions to RBP binding specificity *in vivo*.

Adding another layer of complexity is the high propensity for RBPs to interact with one another, working in tandem to jointly regulate common targets (Dassi 2017). In order to account for the vastness of the transcriptome and the diversity of associated regulatory functions, RBPs are further believed to interact combinatorially, forming complexes of varying sizes and composed of varying components (Khoroshkin et al. 2024); this is yet another attribute that may enable RBPs to achieve their observed degree of binding specificity (Lang et al. 2021; Street et al. 2024). These RBP-RBP interactions may occur via direct (i.e., requiring RNA binding and physical interaction between RBPs) or indirect (i.e., requiring RNA binding but not physical interaction between RBPs) binding modes (Street et al. 2024), and they may operate in cooperative, competitive, or mutual manners (Dassi 2017). Such interactions, which may involve as few as 2 RBPs or as many as required for large macromolecular complexes like the spliceosome, are vital to the fidelity of mRNA regulatory processes (Lang et al. 2021). Previous work has provided experimental evidence of an extensive RBP-RBP interactive network (Dassi 2017; Quattrone and Dassi 2019; Lang et al. 2021; Street et al. 2024), but the biological mechanisms for the majority of these interactions have not yet been determined. In short, the exponentially large landscape and functional implications of the RBP-RBP interactome remain underexplored (Khoroshkin et al. 2024).

Machine learning and deep learning have become increasingly prevalent in genomics applications, especially for the prediction of RBP binding (Alipanahi et al. 2015; Ghanbari and Ohler 2020; Sun et al. 2021; Sharma et al. 2021). While such applications at large have contributed to our current understanding of RBP binding, there remain significant areas that have been under– or never before investigated, such as (1) the structure, composition, and role of motifs to RBP binding and interactions, (2) the salient features within the regions flanking RBP binding motifs, (3) the *in vivo* preferences of RBP motifs and flanking regions, (4) the role of flanking regions in facilitating RBP binding and interactions, and (5) the similarities in flanking regions across RBPs. Furthermore, current motif discovery methods rely on enumerative or probabilistic approaches (D’haeseleer 2006a) but do not consider the contributions of *k*-mers in the surrounding flanking regions when constructing motifs, which may cause noise or inaccuracies to be introduced into the discovered motif. Therefore, we elected to focus on two areas of great import to mRNA regulation: the contribution of motifs and their flanking regions to RBP binding and RBP-RBP interactions, as well as the combinatorial capacity of RBP-RBP interactions. We defined domain requirements to guide our formulation and construction of a comprehensive pipeline to address these RBP knowledge gaps.

Namely, we aimed to develop an architecture-independent solution – to maximize flexibility and generalizability – that balances the performance-interpretability tradeoff intrinsic to neural networks, whose “black box” nature makes their decision-making rationales ambiguous to the user (Hwang et al. 2024). We strove to design an interpretable strategy that achieves high accuracy in RBP binding prediction and motif discovery, both of which are integral to the investigation of motif flanking regions and RBP-RBP interactions. As opposed to commonly-practiced top-down approaches that search for significant regions within a full-length, intact sequence, we instead adopt a bottom-up approach in which our goal is to deconstruct a sequence into its constituent *k*-mers and regions, discover the role and importance of each constituent, and then reconstruct the sequence based on the significant elements that were discovered.

Given that RBPs only bind motif instances that are overrepresented and that are located in a favorable environment with significant flanking regions, we sought to understand the composition, structure, and contribution of these flanking regions to sites of RBP binding events. To this end, we required a method by which to identify the overrepresented *k*-mers that constitute the motif and its proximal flanking regions, the similarities between *k*-mer instances of the binding motif, the relationships between *k*-mers comprising the motif and *k*-mers comprising the motif’s flanking regions, and the relationships between *k*-mers within the flanking regions themselves. The genomics and computational biology fields have speculated that DNA and RNA possess an underlying language and linguistic structure, whose components and inner workings remain elusive (Durbin et al. 1998). In our endeavor to understand the *k*-mers contributing to RBP binding, we regarded RNA as possessing features and structures akin to those found in natural languages and were thus motivated to model RNA using concepts from linguistics and natural language processing (NLP). While natural language has a defined lexicon, syntax, and semantics, as well as established delimiters which enable text parsing, analogous structures and representations of genomic sequences are either missing, difficult to formalize, or not yet discovered. For our work, we decided to model sentences as sequences, phrases as significant regions, and words as *k*-mers. Furthermore, a crucial similitude between natural language and genomic language is that context confers meaning, which makes NLP particularly apt for this study.

Here, we present a novel pipeline that integrates state-of-the-art concepts from natural language processing (NLP), weak supervision, and deep learning, as well as a novel motif discovery algorithm, to bridge the knowledge gaps pertaining to RBP binding patterns. To this end, we devised an NLP-based sequence decomposition strategy that establishes a new representation for candidate RBP binding sites and their flanking regions, which we then used to train neural networks to predict RBP binding under a Multiple Instance Learning (MIL) framework. We also developed a consensus-based motif discovery algorithm to interpret our predictions, demonstrating its ability to accurately recapitulate the motifs of several well-characterized RBPs. Consequently, and importantly, we systematically identified and profiled the flanking regions and binding preferences for 71 RBPs in the HepG2 cell line and 74 RBPs in the K562 cell line, the majority of which are novel, and we conducted a thorough analysis of the similarities and differences in binding patterns between RBPs. Finally, we integrated our discovered motifs and flanking regions with several analysis metrics and a new cross-prediction approach to propose novel cooperative and competitive RBP-RBP interactions, for which we additionally hypothesize underlying mechanisms and functions. Taken together, we present a comprehensive computational approach that enables characterization of the features contributing to RBP binding specificity, as well as investigation into the participants and functions of RBP interactions.

## RESULTS

### A novel computational framework to model RBP binding

We drew on theoretical concepts from linguistics and natural language processing (NLP) to design a novel computational framework to discover the motifs and sequence contexts of RBP binding sites (**Figure 1A**). We utilized merged replicate enhanced cross-linking and immunoprecipitation (eCLIP) sequencing data (**Supplemental Table S1**) (Van Nostrand et al. 2016) in the HepG2 and K562 cell lines from the ENCODE consortium due to its coverage of a large array of RBPs, its standardized, uniformly processed nature, and its use of input controls to ensure high-confident, high-quality peaks (Van Nostrand et al. 2020a). To construct our training dataset, we defined positive sequences as centered eCLIP peaks resized to a uniform length of 101 bp; for our negative set, we performed random sampling of sequences with equivalent length from matched genomic regions (**Supplemental Figure S1A**).

**Figure 1:**
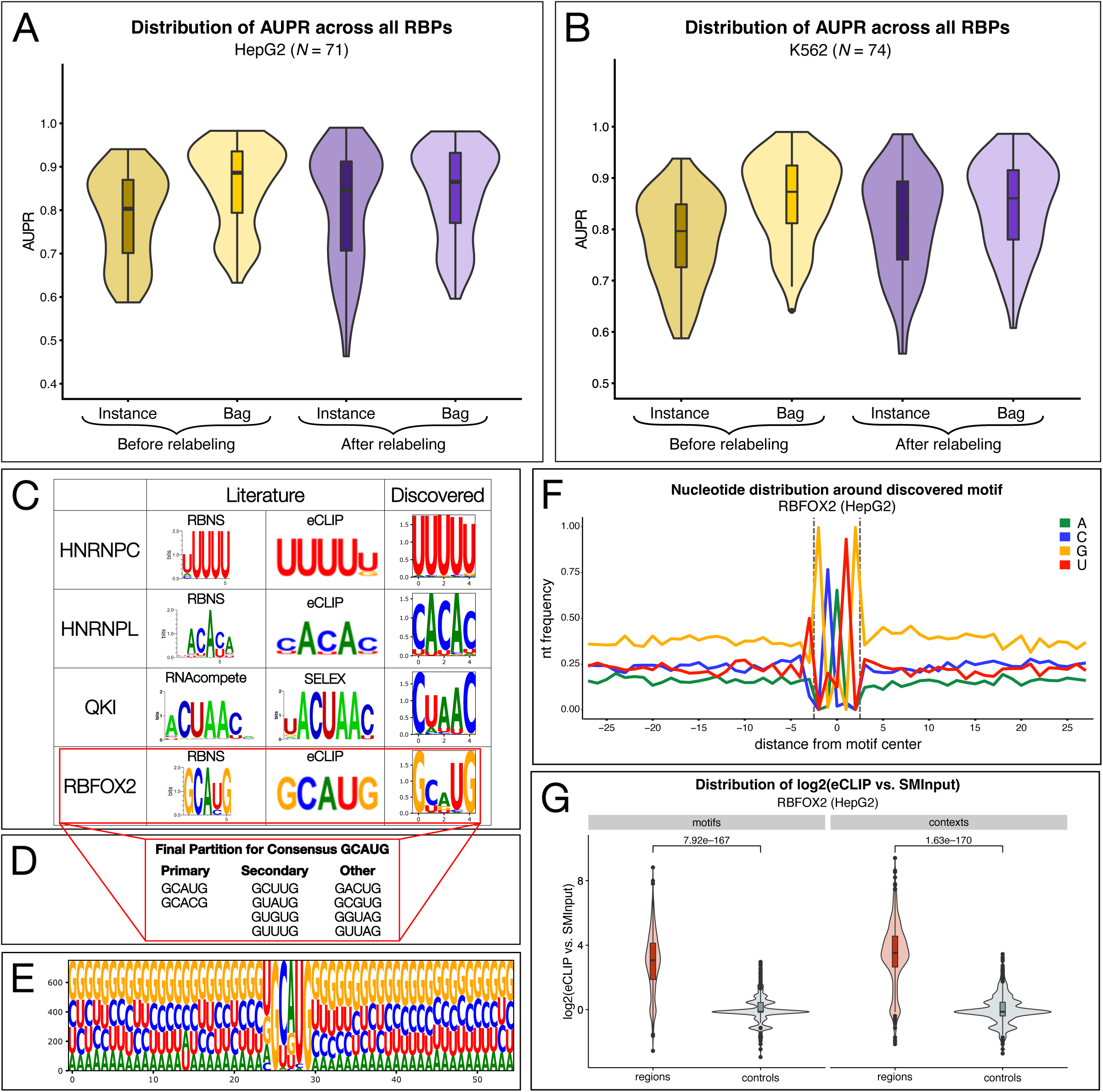
A novel computational framework and algorithm for RBP binding motif and context discovery. (A) Distribution of instance-level and bag-level model performances (in AUPR) across 71 RBPs in HepG2, before and after iterative relabeling. (B) Distribution of instance-level and bag-level model performances (in AUPR) across 74 RBPs in K562, before and after iterative relabeling. (C) Exemplification of motif discovery validation, with respect to the literature, for 4 representative RBPs in HepG2 with motifs of diverse nucleotide content. Literature motifs obtained from (Ray et al. 2013; Dominguez et al. 2018; Van Nostrand et al. 2020a). For each RBP, one *in vitro* and one *in vivo* motif logo is displayed, with the exception of QKI, for which only *in vitro* logos are shown; however, QKI’s motif has been well-characterized *in vivo* by multiple studies, such as in (Hall et al. 2013). (D) Unique instances comprising the RBFOX2 canonical GCAUG motif; 6 motif instances recapitulate known primary and secondary *k*-mers bound by RBFOX2 *in vitro* (Begg et al. 2020). (E) Context logo for RBFOX2; the target motif GCAUG is preceded by a U and enriched in a G-rich sequence context, as previously demonstrated in the literature (Dominguez et al. 2018). (F) Nucleotide preference of RBFOX2 binding, visualized as a line plot. (G) Nested violin-boxplots of the distribution of log2 (eCLIP read enrichment over size-matched input [SMInput]) for discovered motifs (left) and contexts (right), relative to matched controls. Wilcoxon rank-sum test *p*-values test whether the motifs or contexts have significantly eCLIP enrichment than their corresponding control regions.

We first introduced a linguistics-inspired method of sequence decomposition to deconstruct each sequence in our dataset into a series of regions called “contexts”, each of which consists of a central target *k*-mer and its flanking subsequences (**Supplemental Figure S1B**, **S1C**; **Supplemental Note S1**). We used this sequence representation to formulate the RBP binding prediction task as a multiple instance learning (MIL) learning problem (Dietterich et al. 1997; Herrera et al. 2016; Carbonneau et al. 2018), in which we modeled each sequence in our dataset as a “bag” and the sequence’s constituent contexts as the bag’s “instances” (**Supplemental figure S1A**; **Supplemental Note S1**). We then trained weakly-supervised neural networks to perform (1) bag-level classification, to predict whether a sequence is bound or unbound, and (2) instance-level classification, to predict which context(s) within a sequence are responsible for the sequence’s label (**Supplemental Figure S1D**; **Supplemental Note S1**). To remove labeling noise that is inherent to MIL, we also introduced a strategy called iterative relabeling, which helps to filter out “weak” contexts that have little to no contribution to an RBP binding event while simultaneously retaining “strong” contexts that have the most significant contribution to the binding event (**Supplemental Note S1**).

To evaluate how well our models can successfully distinguish between true binding sites and the matched random control sequences, we utilized the area under the precision-recall (AUPR) curve, a performance metric that assesses how well a classifier can achieve good precision (i.e., the positive predictive value, or the proportion of predicted positives that are in fact true positives) and good recall (i.e., the true positive rate, or the proportion of true positives that are correctly predicted positive) (Boyd et al. 2013); a perfect classifier would achieve an AUPR of 1.0. Looking at the distribution of AUPR values across an array of RBPs in each cell line, our models generally performed well, with a final median bag AUPR of 0.865 in HepG2 (**Figure 1A**; **Supplemental Table S2A**, **S2B**) and 0.860 in K562 (**Figure 1B**; **Supplemental Table S2C**, **S2D**). The consistency of performance across both cell lines demonstrates the robustness of our method. Notably, the bag performance was largely preserved before and after relabeling, indicating that our trained models could successfully achieve the removal of labeling noise from positive sequences while simultaneously maintaining bag-level prediction accuracy (**Supplemental Note S1**; **Supplemental Table S3**).

### A novel motif discovery algorithm for accurate discovery of RBP binding motifs and contexts

In order to derive RBP motifs and, importantly, RBP binding contexts directly from the MIL predictions, we required a method of interpretation for our models that could accommodate the context data structure. To this end, we developed a deterministic, consensus-based motif discovery algorithm that relies on empirically-derived measurements of three theoretically significant *k*-mer properties to discover motifs (**Supplemental Note S2**; **Supplemental Figure S2**).

As a proof-of-principle, we first inspected RBFOX2 (HepG2), a well-characterized RBP with established regulatory functions in mRNA splicing and mRNA stability that is implicated in neurological developmental processes and disease (Begg et al. 2020). Our algorithm was able to reproduce RBFOX2’s canonical GCAUG motif, as supported by motifs derived from both *in vitro* and *in vivo* assays (**Figure 1C**, bottom row) (Van Nostrand et al. 2020a). Previous studies have also demonstrated that approximately half of RBFOX2’s binding sites contain alternative, non-GCAUG motifs, which have since been identified *in vitro* after successive rounds of iterative enrichment calculations from a synthetic RNA library modeled solely after 3’UTRs (Begg et al. 2020). Crucially, our discovered motif contains 6 of the 8 *k*-mers identified in the *in vitro* study: 2 primary (GCAUG and GCACG) and 4 secondary (GCUUG, GUAUG, GUGUG, and GUUUG) (**Figure 1D**). To the best of our knowledge, this marks the first documentation of an *in vivo*-derived motif that successfully captures the majority of RBFOX2’s alternative lower-affinity *k*-mers.

Our RBFOX2 motif also contained 4 additional *k*-mers that have not been previously identified. GACUG and GCGUG maintain the critical G1, U4, and G5 positions of the canonical motif; although these 2 *k-*mers were not included in the set of GHNUG *k*-mers characterized in (Begg et al. 2020), they can still be denoted by this IUPAC nucleotide representation (Cornish-Bowden 1985). In conjunction with their relatively frequent occurrence in our discovered motif, especially in comparison to some of the established primary and secondary *k*-mers (**Supplemental Table S4**), these properties suggest that GACUG and GCGUG may serve as novel candidates for lower-affinity RBFOX2 secondary *k*-mers; in fact, GACUG has been previously reported in another study (Zhou et al. 2007). The remaining 2 *k*-mers, GGUAG and GUUAG, which do not share the GHNUG representation and which occur less frequently, may also present novel, lower-affinity secondary motifs with a distinct nucleotide content and lower similarity to the GCAUG consensus, or they may be attributed to noise. We reason to propose that, because they satisfy the same conditions and constraints as the known established motif instances, it is probable that these *k*-mers may be novel motif instances, and we further comment on this later in our work.

Our context logo for RBFOX2 revealed a strong U directly preceding the GCAUG motif, a finding that coincides with its well-profiled canonical *6*-mer motif, (U)GCAUG (Begg et al. 2020). The nucleotide preference of RBFOX2’s motif context also corroborated *in vitro* findings (Dominguez et al. 2018), with RBFOX2 demonstrating a clear predisposition for binding in G-rich regions (**Figure 1E, 1F**). To further validate our discovered motifs and contexts for RBFOX2, we performed a comparative analysis of eCLIP read enrichments relative to matched control regions (**Materials and Methods**). We found that motif instances located within the discovered contexts were very significantly and more strongly enriched in eCLIP read coverage compared to their matched controls containing motif instances not located within the proper sequence context (**Figure 1G**), thus providing further experimental support for our discovered motifs and contexts for RBFOX2 in HepG2.

### Robust validation reveals consistency of motif and context discovery

In total, we report the discovered motifs, contexts, and binding preferences for 71 RBPs in the HepG2 cell line and 74 RBPs in the K562 cell line (**Supplemental Table S5**, **S6**). For all RBPs in our dataset, we further demonstrate that our discovered motifs and contexts are significantly more enriched in eCLIP read coverage compared to their corresponding matched random control regions (**Supplemental Table S7**; **Materials and Methods**), which provides additional support that our method discovers true, high-confidence binding sites located within meaningful sequence contexts that exhibit preferred nucleotide preferences.

To quantitatively validate the algorithm, we demonstrate our ability to recapitulate known motifs from the literature for a series of diverse RBPs. To curate this ground-truth set, we compiled known motifs for 14 well-characterized RBPs in HepG2 and K562 independently (**Materials and Methods**). While the size of these validation sets was the same for both cell lines, two RBPs from HepG2’s validation set (PABPN1, PCBP2) were unavailable in K562, and were thus replaced with two other RBPs (PUM2, TARDBP). Our algorithm correctly discovered the motifs for 13 of 14 RBPs in the respective validation sets, yielding a consistent accuracy of ∼92.9% in both cell lines (**Supplemental Table S8**). We highlight representative examples of discovered motifs from HepG2 in **Figure 1C**. The sole RBP from the HepG2 ground-truth set for which our discovered motif did not correspond with that from the literature was PABPN1, a nuclear poly(A) binding protein (Huang et al. 2023) for which we discovered a motif consensus of GCUGG and for which the poly(A) *5*-mer was not enriched in our dataset (**Supplemental Note S3**; **Supplemental Figure S3**; **Supplemental Table S9A**, **S9B**). Similarly, the sole RBP from the K562 ground-truth set for which our discovered motif did not correspond with that from the literature was RBFOX2, and we demonstrate that the canonical GCAUG is also not enriched in the dataset (**Supplemental Note S3**; **Supplemental Figure S4**; **Supplemental Table S9C-F**).

For a more widespread comparison of context discovery, we compared our discovered contexts with *in vitro*-derived contexts for 8 RBPs that were present in our collection and deemed to have significant nucleotide preferences in (Dominguez et al. 2018), showing that we could recapitulate the putative nucleotide preferences for 5 of them (**Supplemental Table S10A**); for the remaining 3, we closely examined the contexts to explain any discrepancies observed in comparison to the *in vitro* study. For HNRNPL, we discovered a symmetrically oscillating CA-rich context, while an average preference for Cs was reported *in vitro*; however, the *in vitro* context also had a relatively strong underlying presence of As – especially at the positions adjacent to the motif – and displayed a depletion of Gs similar to that observed in our context. These differences were observed despite the fact that our discovered HNRNPL motif consensus (ACACA) was concordant with the *in vitro*-derived motif. Second, while our discovered context for IGF2BP1 did not exhibit a strong preference for any specific nucleotide, it did demonstrate a slight, albeit asymmetric, preference for Cs; on the other hand, the *in vitro* context was strongly C-rich on average. Finally, SFPQ’s discovered context was strongly G-rich, while its *in vitro* average nucleotide preference was reported to be C-rich; interestingly, the more detailed *in vitro* context for SFPQ showed that the context was actually C-rich on the left-hand side of the motif but G-rich on the right-hand side of the motif.

In short, we attribute the discordant context preferences for these 3 RBPs to two main reasons. First, an averaged representation of nucleotide preference can be misleading, as it may obscure the underlying position-specific preferences. Second, and perhaps with the most implications, in general *in vitro* experiments are inherently limited by (1) the synthetic, randomized nature of the RNAs used, which may not reflect the intrinsic nucleotide content observed in the transcriptome (e.g., repeat regions), (2) the short length of the utilized RNAs, which may not fully encompass the native context required *in vivo* for RBP binding, and (3) the absence of a natural cellular environment, causing RBPs to be studied in isolation without the presence of endogenous *trans*-factors and other signals that may influence its binding partners or target RNAs (Khoroshkin et al. 2024). The accuracy of our discovered contexts is supported not only by the 5 RBPs that corroborated *in vitro* findings, but additionally and importantly by the facts that (1) we previously established that our discovered motifs are accurate, which in turn suggests that their corresponding contexts must be valid, (2) the accuracy of our motif discovery algorithm demonstrates that the *k*-mer properties used to derive motifs are valid, and (3) two of those properties – namely, *k*-mer enrichment and *k*-mer co-occurrence – use information from the context of the motif instances to construct the motif itself.

To further demonstrate the robustness of our framework, we applied our method to RBP binding datasets from other CLIP-based assays: namely, iCLIP (König et al. 2010), HITS-CLIP (Licatalosi et al. 2008), and PAR-CLIP (Hafner et al. 2010) (**Supplemental Figure S5**). For RBPs that were present in our eCLIP dataset, we demonstrate *in vivo* motif consistency with other CLIP assays as additional validation of the high accuracy of our model. Indeed, the discovered motifs for RBFOX2 (GCAUG), QKI (CUAAC), and SRSF1 (GAGGA) corresponded with those from our eCLIP datasets (**Supplemental Figure S5A**, **S5B**). Interestingly, we noticed slight differences in the contexts of these motifs, and nucleotide preferences were generally less striking. For example, for RBFOX2 iCLIP, while a strong U preceded the canonical GCAUG motif as expected, the discovered context was not G-rich as in our HepG2 results (**Supplemental Figure S5A**, left). Similarly, for QKI HITS-CLIP, while an A preceded the canonical CUAAC as expected, the discovered context was not as strongly AU-rich as in our HepG2 and K562 results (**Supplemental Figure S5B**, left). We attribute these observations to differences in the CLIP variants: while eCLIP ensures that peaks are highly enriched over a size-matched input, which serves as a filtering mechanism to retain only high confidence binding sites that are most likely to be functional, other CLIP assays may use less stringent filtering that might result in the capture of sequence-specific but transient interactions that are not necessarily functional (Ramanathan et al. 2019), thus influencing the observed nucleotide preference.

Application of our framework to other CLIP-based assays also exemplifies the broad generalizability of our method, which allows for the discovery of RBP binding motifs from more diverse data sources than solely eCLIP. To this end, we also demonstrated our discovery of motifs for RBPs that were not present in our eCLIP dataset. Three of these RBPs had U-rich motifs: LSM6 is known to bind oligo(U)’s (**Supplemental Figure S5A**, middle) (Achsel et al. 1999, 2001), EIF4G1 has been shown to bind oligo(U)’s in yeast (**Supplemental Figure S5A**, right) (Zinshteyn et al. 2017), and ELAVL1 recognizes U-rich motifs in *in vitro* assays (**Supplemental Figure S5C**, left) (Ray et al. 2013). HNRNPA1’s motif contains the core AG(G) motif (**Supplemental Figure S5C**, middle) (Abdul-Manan and Williams 1996; Beusch et al. 2017), while IGF2BP1 contains AC-repeats present in previously published motifs (**Supplemental Figure S5C**, right) (Van Nostrand et al. 2020a; Ray et al. 2013). Lastly, we present a novel GC-rich motif for MSI2, for which previous studies typically report UAG but that was not enriched in our dataset (**Supplemental Figure S5B**, right; **Supplemental Table S9G**) (Duggimpudi et al. 2018). We further note that these CLIP datasets originated from cell types other than HepG2 and K562, which demonstrates that our framework is also generalizable to diverse cellular contexts.

Finally, as a negative control, we used our algorithm to perform motif discovery on a set of random genomic and random synthetic sequences (**Supplemental Note S4**; **Materials and Methods**). Despite the fact that both our algorithm and STREME (Bailey 2021) returned motif hits for this set of negative controls (**Supplemental Figure S6**, **S7**), we noted several distinctive differences in sequence composition and feature constitution that clearly distinguish the motifs and contexts from RBP datasets from those discovered from random sequences. First, MIL binary classification on the random sequences performed only as well as a random classifier (i.e., an AUPR of around 0.5 when 50% of observations in a dataset are positive), indicating that these models are unable to differentiate between the foreground and background distributions present within the random dataset. This performance was significantly worse than the performance for both a subset of sequence-specific RBPs and a set of non-sequence-specific RBPs (**Supplemental Figure S8A**, S**8B**). We also found that (1) the enrichment values for the motif consensuses of random motifs were significantly lower than those from sequence-specific and non-sequence-specific RBP motifs (**Supplemental Figure S8C**), and (2) the *p*-values of motif consensuses from random motifs were higher than those from sequence-specific and non-specific RBP motifs (**Supplemental Figure S8D**). We observed similar findings for the enrichment and *p*-values of all *k*-mers in each of these datasets, with the random datasets also displaying much fewer numbers of significantly enriched *k*-mers overall (**Supplemental Figure S9**). Across all of these features, sequence-specific and non-sequence-specific RBPs displayed comparable properties with one another.

Taken together, these analyses demonstrate that it is the properties of the discovered motifs and contexts, rather than the presence or absence of a hit by a motif discovery algorithm, that are informative of a good negative control (**Supplemental Note S4**). Further, these analyses demonstrate that our algorithm discovers meaningful RBP motifs and contexts that are clearly distinct from those discovered from random genomic and random synthetic sequences. This is of particular importance for interpreting the results of non-sequence-specific RBPs, which are typically known to bind to the RNA phosphate backbone rather than to nucleotide bases. We hypothesize that the discovered motifs and contexts for non-sequence-specific RBPs, which display preferences distinct from random negative controls, may be reflective of their frequent association with other sequence-specific RBPs, thus causing the CLIP crosslinking and pulldown to capture regions of RNA that demonstrate the sequence preferences of the interacting partner; we propose such a case later in this work. Importantly, these analyses definitively demonstrate that non-sequence-specific RBPs do not constitute a valid negative control for motif discovery, as their motifs and contexts are significantly different from those discovered from random sequences; thus, although these RBPs are classified as non-sequence-specific, the sequences associated with their binding sites are non-random.

### Motif and context discovery across cell lines reveals similarities and differences in binding preferences

We sought to comprehensively characterize the discovered binding contexts for the set of RBPs covered in our study. In the HepG2 cell line, we discovered novel *in vivo* contexts for a total of 71 RBPs. For the contexts of the 5 RBPs for which we corroborated with *in vitro* data (**Supplemental Table S10A**), we provide additional detail by reporting a larger context region, but also by delineating nucleotide preferences for each component of the context (below). For other RBPs, such as those known to bind polypyrimidine tracts (e.g., PTBP1, U2AF2) (Sayers et al. 2022), we recapitulated the nucleotide preferences of their contexts, but also provided a more detailed report of these preferences per position. For the remaining RBPs, which comprise the majority of the set, we present for the first time, to the best of our knowledge, a comprehensive profile of their preferred contexts for binding *in vivo*.

First, we comprehensively determined the nucleotide preferences for each RBP’s motif and context (**Supplemental Table S10B**). Notably, we observed a strong degree of symmetry for the majority of our discovered contexts (*N* = 59; 83.1%), where we define symmetry to occur if the left and right contexts flanking the motif share the same nucleotide preference (**Materials and Methods**). We noticed that the symmetric preference was for either a single nucleotide or for 2 frequently occurring nucleotides. The remaining 12 RBPs had contexts that were asymmetric in nucleotide preference, which displayed a bit more variance. In the case of ILF3, for example, each side of the context displayed preference for a single, albeit different, nucleotide, where its left context showed preference for C but its right for G. In other cases, each flank of the context displayed preference for 2 nucleotides of different identity (e.g., IGF2BP1), or the left and right contexts each displayed preference for a different number of nucleotides (e.g., PABPN1). By visualizing the position probability matrices (PPMs), we noted that some RBPs had stronger, more pronounced nucleotide preferences than other RBPs.

Furthermore, we observed that the majority of RBPs (*N* = 46; 64.8%) had preference for the same nucleotide in both their motif and context (**Supplemental Table S10B**; **Materials and Methods**); for example, G3BP1 is AG-rich in both its motif and contexts. For such RBPs in which the context’s nucleotide preferences appear to be extensions of the motif’s nucleotide preferences, we also considered that this pattern may represent a known phenomenon in which multiple motif sites occur together to form clusters (**Supplemental Note S5**; **Supplemental Figure S10**). While the remaining 25 RBPs had different nucleotide preferences in their motif and contexts, the difference in preference was generally not very striking. For example, while HLTF’s motif is AG-rich, its contexts are overall A-rich, indicating that there is still some degree of similarity between the nucleotide preference of each element. One exception to this phenomenon is QKI, which demonstrates a marked difference in the nucleotide preferences of its elements, having an AC-rich motif but a U-rich context (**Supplemental Table S5**; **Supplemental Table S10B**).

We next performed the same analyses for the 74 RBPs in K562. Similarly to HepG2, the vast majority of RBPs (*N* = 60; 81.1%) in K562 exhibited symmetric nucleotide preferences in their contexts, while only 14 (18.9%) had asymmetric contexts (**Supplemental Table S10C**). The majority of RBPs also had the same nucleotide preference in the motif and context (*N* = 49; 66.2%) (**Supplemental Table S10C**). We next compared the discovered binding patterns for RBPs with datasets in both cell lines, for which there were 40 RBPs. Broadly, we found that 60% of these RBPs shared preferences for the same nucleotide in their motifs, while 67.5% of RBPs shared preferences for the same nucleotide in both the left context and right context, respectively, and 65% of RBPs shared the same preference across both sides of the context (**Figure 2A**). An even greater majority, 82.5%, displayed the same pattern of (a)symmetry across cell lines, while 70% exhibited the same nucleotide preference in both the motif and context.

**Figure 2:**
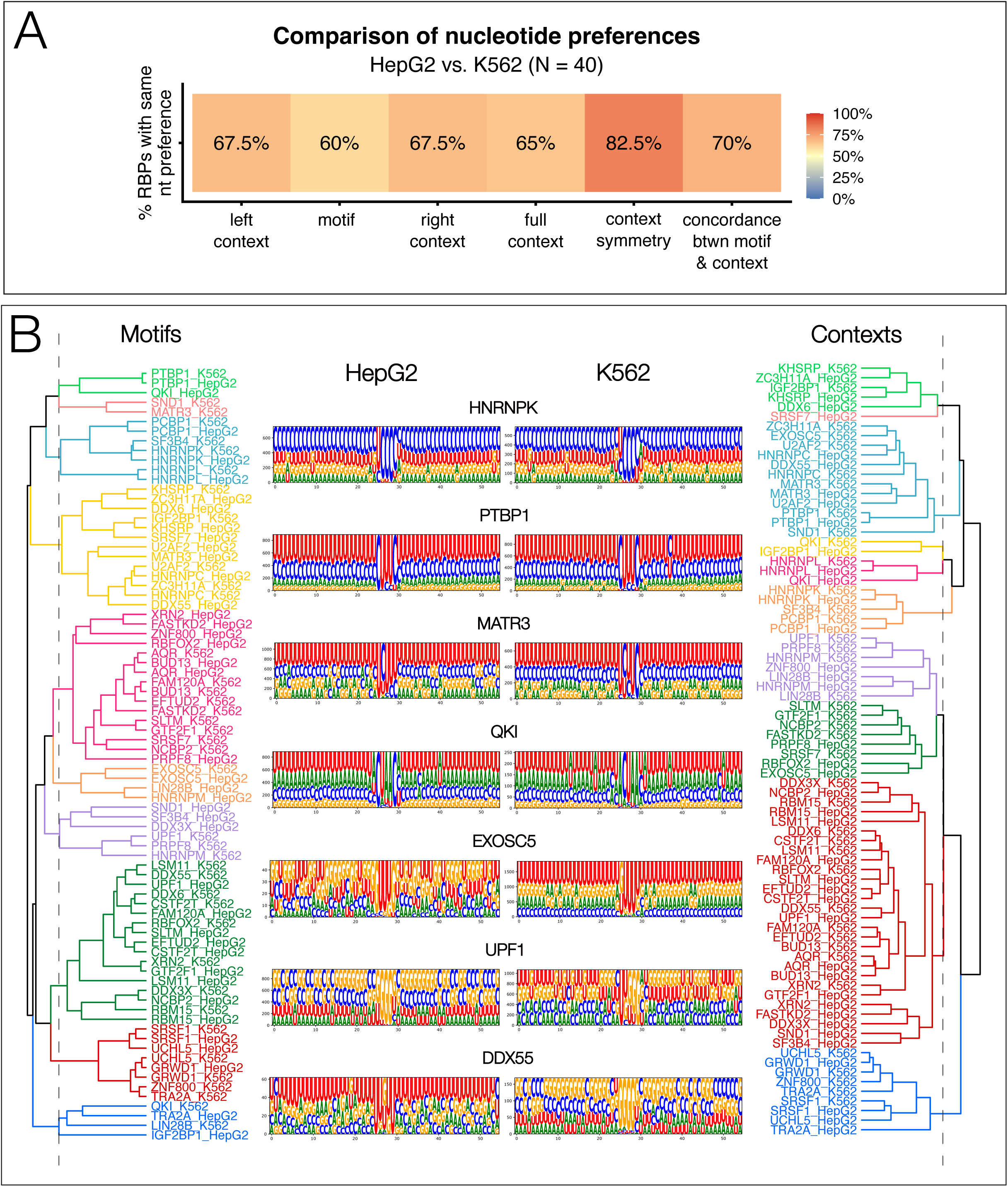
Motif and context nucleotide preferences in HepG2 and K562. (A) Comparison of nucleotide preferences between common RBPs in both cell lines. (B) Clustering of motif (left) and context (right) position probability matrices (PPMs) for common RBPs in both cell lines.

We investigated the degree of similarity in the motifs and contexts between the two cell lines for a given RBP by assessing whether the RBP from both HepG2 and K562 were assigned to the same cluster upon hierarchical clustering of their PPMs (**Figure 2B**). Only 45% of RBPs (*N* = 18) from one cell line clustered together by motif PPM with their counterpart from the other cell line (**Figure 2B**, left), while 57.5% (*N* = 23) clustered together by context PPM (**Figure 2B**, right), suggesting that both motifs and contexts may differ to a sizable degree in a cell type-specific manner. Some RBPs, such as HNRNPK and PTBP1, had strikingly similar motifs and contexts in both cell lines (**Figure 2B**, middle). For some RBPs that did not cluster together with their counterpart in the other cell line, the differences were visually apparent, such as with MATR3, whose motifs differed strongly by a single nucleotide but whose contexts were otherwise similar; for others, the difference could be explained by the fact that one motif was a “shadow” of the other (e.g., with QKI, whose motif in HepG2 was identified as [A]CUAAC but whose motif in K562 was identified as ACUAA). On the other hand, more substantial discrepancies were present for some RBPs that had similar motifs but different contexts in the two cell lines (e.g., EXOSC5) or vastly different motifs and contexts between the two cell lines (e.g., UPF1, DDX55). While these analyses can be considered preliminary, this work demonstrates the importance of studying RBP binding patterns in a cell type-specific manner, and future work will be important to investigate the degree of cell type-specificity’s effects on RBP motifs and contexts as well as their functions and implications in RNA regulatory mechanisms.

### A comprehensive characterization of the binding motifs and contexts for 71 RBPs in HepG2

For a closer, more detailed, and higher resolution inspection of RBP binding preferences, we next decided to focus on our results from the HepG2 cell line for the remainder of our work. To gain an improved understanding of our discovered motifs for the full set of 71 RBPs in HepG2, we classified each into one of 5 categories (**Supplemental Table S11**):

● <u>Consistently reported</u> (*N* = 14): The RBP has a well-characterized binding motif that has been consistently reported across multiple sources in the literature.
● <u>Multiple reported</u> (*N* = 22): Multiple different motifs have previously been reported for the RBP; we either present a motif that supports at least one of these reported motifs or a novel motif that does not correspond with any of those previously reported.
● <u>Weakly characterized</u> (*N* = 25): Limited information is available regarding the binding motif of the RBP (e.g., only 1 source available, only nucleotide preferences [but not a concrete motif] available); we present a more specific motif.
● <u>Non-sequence-specific</u> (*N* = 6): The RBP is believed to recognize features other than sequence, such as RNA secondary structure.
● <u>No existing characterization</u> (*N* = 4): To the best of our knowledge, no motif has been previously reported for the RBP, and we present a novel binding motif.

We have addressed the “Consistently reported” classification in the preceding section. In our “Multiple reported” category, our discovered motif for a given RBP often corresponded with at least one of the reported motifs. For example, FUBP3, an RBP for which we discovered a motif with a UGUGU consensus, corroborated the UG-repeat motif discovered in an *in vivo* study (Gau et al. 2011) but was distinct from the UA-rich motifs discovered *in vitro* (Dominguez et al. 2018); this discrepancy could be attributed to technical reasons underlying the environments of the respective assays used to generate the binding data. On the other hand, discrepancies between multiple reported motifs could also be attributed to biological reasons based on the RBP’s binding mode. For example, KHSRP, an RBP belonging to the same protein family as FUBP3 (Gau et al. 2011) and for which we also discovered a UGUGU motif, has been shown to bind both GU-rich and AU-rich elements *in vivo* (Briata et al. 2016; Cook et al. 2011). KHSRP has 4 KH RBDs, each having varying degrees of specificity, with the third KH domain in particular demonstrating a strong preference for G-containing sequences (García-Mayoral et al. 2008; Palzer et al. 2022). We thus hypothesize that our UGUGU motif may reflect that KHSRP’s utilization of KH-3 for RNA binding was more prevalent during eCLIP capture. Our KHSRP motif also exhibited a characteristic depletion in Cs that has previously been observed (García-Mayoral et al. 2008) (**Supplemental Table S5**).

We also examined the additional motifs that were discovered by our algorithm but not ranked first by our scoring strategy, which we refer to collectively as secondary motifs. In the case of LIN28B, we discovered a GGAGA secondary motif that supported the GGAG motif discovered for LIN28B in a previous study of its transcriptomic targets (Graf et al. 2013) as well as a GGAGA motif discovered for LIN28B’s paralog, LIN28A, with which it shares a similar structure (Wilbert et al. 2012; Maklad et al. 2023). This demonstrates that our method still has the power to discover previously reported motifs, even if not ranked first by our scoring method.

For our “Weakly characterized” category, we propose new, specific motifs for 25 RBPs that we considered to be weakly characterized. For 10 of these RBPs, a previous study had reported their candidacy for binding to rG4s (Lee et al. 2020; Busa et al. 2021; He et al. 2021), and our discovered motifs (and contexts) – which were all G-rich – supported this theory. For some RBPs that only had a reported motif from a single source, our discovered motifs were concordant, thus providing additional support; for example, PPIG and PRPF8, which both have GUGAG motifs (**Supplemental Table S5**) (Feng et al. 2019). For other RBPs, we propose novel motifs (e.g., GGGUG for BUD13, CACAC for CDC40, GAAGA for FXR2; **Supplemental Table S11**).

Our “Non-sequence-specific” category contains a number of RBPs that have previously been hypothesized to bind in a non-sequence-specific manner. Many of these RBPs are DEAD-box helicases, which recognize the RNA sugar-phosphate backbone as opposed to its nucleotides (Linder and Jankowsky 2011), while the other two have either been reported to have double-stranded RBDs (ILF3) (Li et al. 2020) or to bind highly structured RNAs (XPO5) (Wang et al. 2020). We nevertheless present novel binding motifs for these RBPs, hypothesizing that the motifs may instead reveal the sequence preferences of other *trans*-factors that interact with the RBP in question, as we elaborate on in the following sections. Finally, to the best of our knowledge, the motifs for 4 of the 71 total RBPs have not been previously characterized, comprising our “No existing characterization” category. This set includes EXOSC5, HLTF, RPS3, and ZNF800.

### Investigating the relationships of binding patterns between RBPs

Equipped with our characterization of RBP binding preferences, we next sought to understand the similarities and differences in these preferences between RBPs. First, we used HDBSCAN (McInnes et al. 2017) to cluster the flattened PPMs of the motifs and contexts, respectively, of all RBPs, then visualized the dimension-reduced clusters using a two-dimensional UMAP embedding (McInnes et al. 2018) (**Figure 3A, 3B**). Using HDBSCAN and UMAP with the Euclidean metric to compute distances between points – with the former technique seeking to assign clusters based on point proximity and the latter technique aiming to preserve the relative distances observed in high-dimensional space in its lower-dimensional mapping – we can use the results as a method of motif comparison, where a small Euclidean distance and same cluster membership suggest that two RBPs in question may possess similar motif and context patterns, as well as similar nucleotide binding preferences.

**Figure 3:**
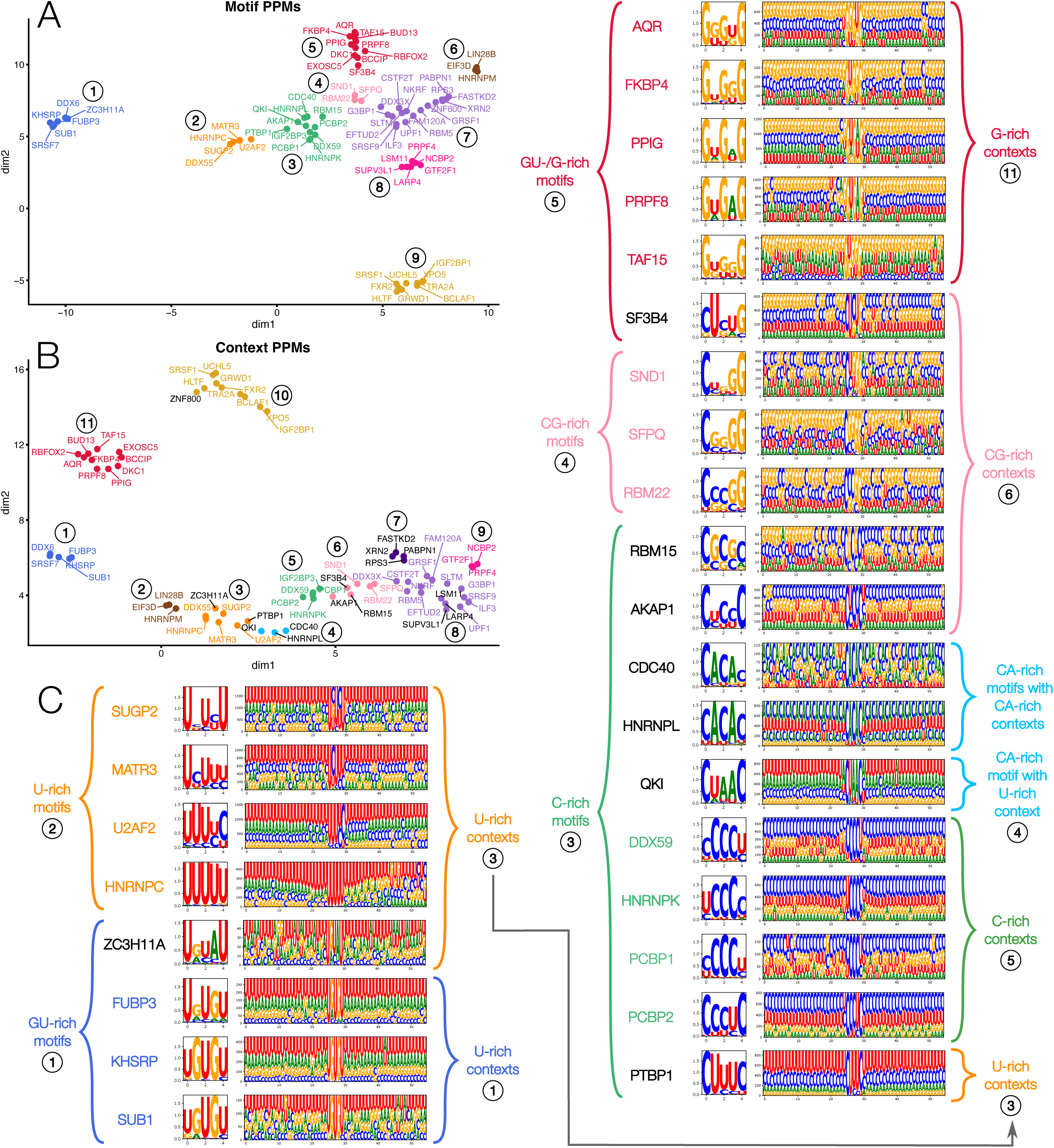
RBPs cluster based on similarities between their respective motifs and contexts. Clustering of motif (A) and context (B) PFMs with HDBSCAN and visualization with UMAP. Clusters are assigned number labels, with cluster colors roughly mapping between the two embeddings. (C) Motif and context logos for representative RBPs from selected clusters; cluster labels and nucleotide preferences underlying the clustering assignments from panels A and B for target (left) and context (right) motifs are labeled in the periphery. RBPs whose names are labeled in black are those that migrated clusters between the target and context clustering, with the cause for migration evident from the corresponding logos.

Indeed, we observed that RBPs clustered together by nucleotide preference. Clustering by motif yielded 9 clusters in total (**Figure 3A**), with distinct groups forming for RBPs with generally UG-rich (Cluster 1), U-rich (Cluster 2), C-rich (Cluster 3), CG-rich (Cluster 4), and A-rich (Cluster 9) motifs, each with a moderate degree of variation (**Figure 3C**). Cluster 6 contained 3 RBPs with relatively different motifs from one another but are characterized by their strong G1 and U5 positions. The last three clusters consisted of RBPs with largely G-rich motifs, but their division into subgroups can be attributed to the presence of a distinct C5 position (Cluster 8), strong G’s at the odd-indexed or solely peripheral positions and U’s at either or both of the even-indexed positions (Cluster 5), and the remaining, predominantly G-rich motifs (Cluster 7). The latter cluster is fairly large (*N* = 17), with a considerable degree of variation present, but with less well-defined patterns, which may help to explain why it was not further divided into subgroups.

In our context-based clustering results, we observed that the majority of RBPs remained clustered with the same members as in the motif-based clustering, with 16 of the 71 (22.5%) migrating to a new cluster (**Figure 3B**, labeled in black). We sought to understand the underlying causes that fostered these migrations, and we did so by carefully inspecting the motif and context logos of the associated RBPs (**Figure 3C**). For example, due to its strong CUUUC consensus, PTBP1’s motif was distinct enough from the set of RBPs having U-rich motifs (Motif Cluster 2, orange) and instead clustered with the set of RBPs having C-rich motifs (Motif Cluster 3, green). However, upon the addition of context, PTBP1’s overwhelmingly strong preference for U’s in its binding context led to its subsequent clustering with the set of RBPs having U-rich contexts (Context Cluster 3, orange), as opposed to remaining in the same cluster that it previously belonged to, whose members ultimately have C-rich contexts (Context Cluster 5, green). A similar phenomenon was observed for CDC40, HNRNPL, and QKI which, despite demonstrating a binding preference for C’s and A’s, have motifs with strong enough C’s that they were also clustered with the set of RBPs having C-rich motifs (Motif Cluster 3, green). When contexts were incorporated, these 3 RBPs detached to form their own cluster (Context Cluster 4, blue). CDC40 and HNRNPL both had CA-rich contexts, although the latter had strong U undertones. Interestingly, in spite of its context preference for Us, QKI was still included in this cluster, likely due to its nucleotide preference similarities in motif with CDC40 and HNRNPL. This demonstrates a case in which the motif is more influential than the context in the clustering outcome. In short, the migration of certain RBPs between the motif and context clustering demonstrates that context plays an important role in differentiating between binding outcomes, particularly for RBPs sharing similar target motifs.

We next sought to examine the similarities and differences between RBP binding patterns using a probabilistic approach, reasoning that the PPM representation of motifs and contexts fundamentally comprises a series of position-specific probability distributions across the 4 canonical nucleotides. To this end, we used the Kullback-Leibler Divergence (KLD) to quantify the degree to which one probability distribution *Q* (i.e., the approximating distribution) can approximate another probability distribution *P* (i.e., the reference distribution), where smaller values of KLD represent a closer approximation (Bishop and Nasrabadi 2006). Unlike the Euclidean distance metric, the KLD does not qualify as a *bona fide* metric in the formal, mathematical sense due to its asymmetric nature, and thus we instead refer to KLD as a measure of comparison. By computing the cumulative KLD across each position of a given pair of PPMs, we can obtain a quantitative method to compare PPMs between RBPs (**Materials and Methods**). For this reason, we used the KLD as a quantitative measure of the distributional similarities and differences between RBP motifs and contexts for all future analyses (**Supplemental Note S6**; **Supplemental Figure S11**).

We also computed the cumulative KLD between RBPs in a pairwise manner, performed hierarchical clustering on the global KLD profiles across all RBPs, and annotated each branch in the resulting dendrogram based on the corresponding nucleotide preferences of the motif (**Figure 4A**) and context (**Figure 4B**). Groups of RBPs with mutually low KLDs are visible as patches of blue in the heatmap. Notably, the hierarchy of the dendrogram allowed for ease of visualization and interpretation of the comparison between RBP binding patterns. For example, for motif KLDs (**Figure 4A**), the first main branch of the dendrogram splits based on whether the motif is rich in G’s or rich in C’s, U’s or A’s; because this is a major divide, and based on the number of RBPs in each of the two sets, we observe that (1) the majority of RBPs in this collection have predominantly G-rich motifs (albeit with some variation) and (2) the preference for G’s is a major differentiator between the two main groups. On the other hand, when we performed the same analyses for the context KLDs, we observed deviations from the patterns observed in the motif KLD clusters, in that the nucleotide preferences at the context level did not always heavily influence the clustering of RBPs; some clusters exhibited a strong, consistent nucleotide preference, while other clusters did not. Such behavior may be attributed to (1) the increased length and complexity upon addition of the contexts, (2) the combined influences of both the motif and the contexts and/or (3) the overall nucleotide preference of each RBP not being the decisive factor in clustering, as RBPs may have contexts with sufficient similarity to be assigned to same cluster while also having different nucleotide preferences (**Figure 4B**).

**Figure 4:**
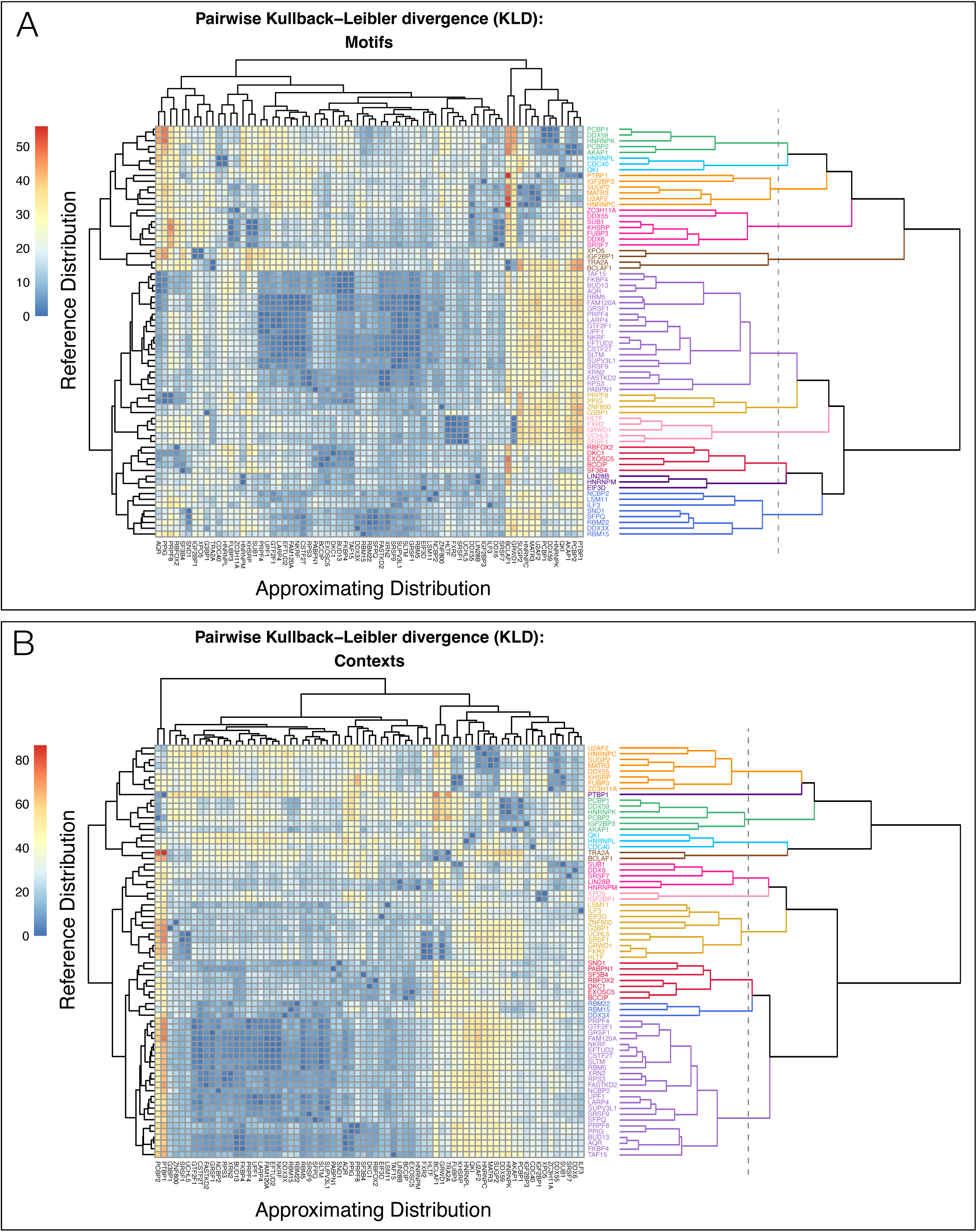
Clustering of cumulative Kullback-Leibler Divergence (KLD) values. (A) Heatmap and dendrogram of cumulative motif KLDs between RBP pairs. (B) Heatmap and dendrogram of cumulative context KLDs between RBP pairs.

### Enrichment analyses in differential gene regulation demonstrate the functionality of discovered contexts

To demonstrate the functionality of our discovered motifs and contexts, we leveraged the RBP knockdown (KD) shRNA-seq datasets from the ENCODE Consortium, for which data were available for 60 RBPs in our HepG2 dataset (**Supplemental Table S1D**) (Luo et al. 2020; Van Nostrand et al. 2020a). We first used DESeq2 (Love et al. 2014) to identify differentially expressed genes upon KD of each RBP, then computed the degree to which our discovered contexts were enriched in differentially expressed (DE) genes relative to a background expressed gene set (**Materials and Methods**). To closely examine the functional effects of each RBP’s KD, we stratified these context enrichment calculations based on (1) the direction in which a DE gene was over– or under-expressed (i.e., up– or down-regulated) and (2) the genomic region in which the RBP’s context was located, with a specific focus on regions known to most strongly influence gene expression (i.e., 5’UTR, CDS, 3’UTR) (**Materials and Methods**). In total, we found that 45 of the 60 RBPs with KD shRNA-seq data (75%) had enriched contexts in DE genes in at least one set of conditions (i.e., in any group defined by a direction-region pair; **Figure 5A**). Furthermore, 23 RBPs demonstrated enriched contexts in up-regulated genes, while 39 RBPs demonstrated enriched contexts in down-regulated genes, suggesting that most of the RBPs in this dataset positively regulate gene expression under normal conditions. Some RBPs had enriched contexts in genes that were differentially expressed in both directions, while other RBPs had enriched contexts in more than one genomic region of differentially expressed genes.

**Figure 5:**
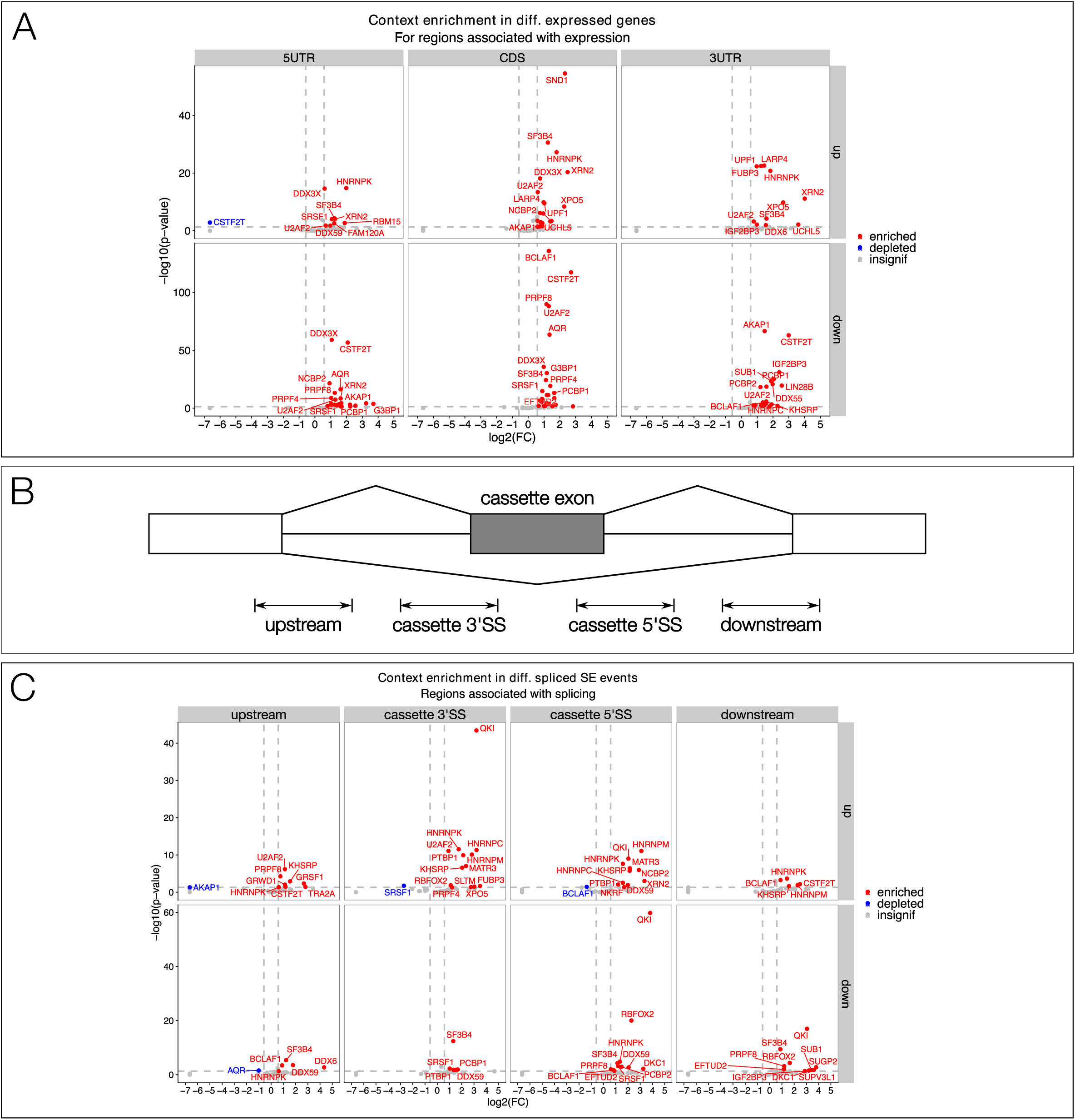
Significant enrichment of discovered contexts in RBP KD experiments reveals functional insights. (A) Volcano plot of RBP context enrichment in up-(top row) and down-(bottom row) regulated genes upon RBP KD, and stratified by whether the context enrichment is observed in 5’UTRs (left column), CDSs (middle column), and 3’UTRs (right column). For each panel, the top 12 most significant points are labeled with the RBP name. (B) Schematic demonstrating the 4 regions of interest for differential splicing analyses. (C) Volcano plot of RBP context enrichment in up-(top row) and down-(bottom row) differentially spliced (DS) skipped exon (SE) events upon RBP KD, and stratified by the region of interest in which the context enrichment is observed. Up-regulation of events indicates increased exon inclusion, while down-regulation of events indicates increased exon skipping.

23 of the 45 RBPs with enriched contexts in DE genes have previous annotation of regulatory roles in gene expression-related functions such as 3’end processing, RNA stability, or translation, thus representing 88.5% of the RBPs in our dataset that had annotations for these functions (Van Nostrand et al. 2020a). For the remaining 22 RBPs, 11 have annotations for roles in splicing. The final 11 are primarily composed of RBPs that have been previously annotated as “Novel RBPs” (Van Nostrand et al. 2020a); therefore, based on their significant context enrichment in differentially expressed genes upon RBP KD, we hypothesize that these RBPs (DDX55, DDX59, FAM120A, FUBP3, GTF2F1, HLTF, NKRF, SUB1, UCHL5) have functions in the regulation of gene expression.

We hypothesized that RBPs with enriched binding contexts located in specific genomic regions of up– or down-regulated genes would be indicative of their regulatory functions (**Figure 5A**). Because 3’UTRs are hotspots for binding by RBPs that regulate mRNA stability, we first examined context enrichments in 3’UTRs. UPF1, whose discovered contexts were primarily located in 3’UTRs (**Supplemental Figure S12A**), demonstrated significantly enriched contexts in the 3’UTRs of target genes that were up-regulated upon UPF1 KD (enrichment = 1.95; *p*-value = 4.71e-23; **Figure 5A**, top right), which supports its known function as an RNA helicase that plays important roles in diverse RNA decay pathways (Kim and Maquat 2019). On the other hand, CSTF2T, a component of the cleavage stimulation factor (CSTF) complex with regulatory roles in 3’end processing and alternative polyadenylation (APA) (Yao et al. 2013), demonstrated significant context enrichment in the 3’UTRs of target genes that were down-regulated upon CSTF2T KD (enrichment = 7.93; *p*-value = 1.15e-63; **Figure 5A**, bottom right), suggesting that the absence of CSTF2T may lead to widespread alterations in 3’end processing that ultimately destabilize its RNA targets. CSTF2T’s functions are generally not well-characterized, and its additional context enrichment observed in other genomic regions of its downregulated targets upon CSTF2T KD (**Figure 5A**, bottom left and middle) may indicate additional regulatory roles, particularly given that the majority of its discovered contexts (∼78%) reside in proximal and distal introns (**Supplemental Figure S12B**). As a final example in 3’UTRs, IGF2BP3 exhibits context enrichment in both up-(enrichment = 1.96, *p*-value = 0.00757) and down-regulated (enrichment = 5.26, *p*-value = 9.52e-32) genes, which aligns with its known functions in regulating RNA degradation (Liu et al. 2024) and RNA stability (Huang et al. 2018), respectively (**Figure 5A**, right; **Supplemental Figure S12C**).

We next sought to validate our discovered motifs and contexts in relation to splicing regulation. We used rMATS-turbo (Wang et al. 2024) to detect and quantify differential splicing events between the knockdown and respective control experiments for each RBP, focusing primarily on the most prominent class of event – skipped exon (SE) (**Materials and Methods**; **Supplemental Figure S13**, **S14**). Because splicing events occur in very localized regions, we first defined 4 regions of interest to analyze [adapted from (Yee et al. 2019); **Figure 5B**], then computed the degree to which our discovered contexts were enriched in differentially spliced (DS) SE events relative to a background set, stratified by direction of regulation and region in which the context is located (**Figure 5C**; **Materials and Methods**).

In total, we found 34 of the 60 RBPs with KD shRNA-seq data (56.7%) had enriched contexts in DS events in at least one set of conditions. 23 RBPs had enriched contexts in up-regulated events, while 18 RBPs had enriched contexts in down-regulated events. 19 of the 34 RBPs (55.9%) with enriched contexts in DS events have previous annotation of regulatory roles in splicing regulation and the spliceosome, thus representing 67.9% of the RBPs in our dataset that had annotations for these functions (Van Nostrand et al. 2020a). For the 9 RBPs in our dataset with previous annotation of splicing but that did not have enriched contexts in DS events, we hypothesize that some may have more predominant functions in other classes of splicing events, such as BUD13, which is known to suppress intron retention (Frankiw et al. 2019), while others have very small datasets that ultimately impacts the detection of context enrichment in the identified DS events, such as SRSF9, which has only ∼3000 sequences in its starting dataset. For the remaining 15 RBPs of the 34 with enriched contexts in DS events, 11 also had enriched contexts in DE genes, while 4 did not; we hypothesize novel functions in splicing regulation for these RBPs. In total, across our differential expression and differential splicing analyses, we detected context enrichment for 52 of the 60 RBPs (86.7%) with KD shRNA-seq data.

We discovered direction– and region-specific context enrichment for RBPs with known splicing patterns that are informative of RBP function. For example, RBFOX2 demonstrated significantly enriched contexts at the 5’ splice site (SS) of cassette exons that were down-regulated upon RBFOX2 KD (enrichment = 4.85, *p*-value = 9.67e-21; **Figure 5C**, bottom row, 3^rd^ column). Furthermore, its motif consensus GCAUG was also significantly enriched in the cassette exon 5’SS of down-regulated events for both event-centric (enrichment = 1.67, *p*-value = 1.77e-16, **Supplemental Figure S15A**) and context-centric *k*-mer enrichments (enrichment = 1.87, *p*-value = 1.62e-04, **Supplemental Figure S15B**) (**Materials and Methods**). Indeed, RBFOX2 is known to promote exon inclusion when bound to sites proximally downstream of the cassette exon (Yeo et al. 2009), which supports our findings. In a similar vein, RBFOX2 has been observed to promote exon skipping when bound to sites proximally upstream of the cassette exon, albeit with a weaker signal relative to its downstream binding sites (Yeo et al. 2009). We observe similar findings, with RBFOX2’s contexts being ∼2.3 fold more enriched in the cassette 5’SS of down-regulated events than in the cassette 3’SS of up-regulated events (enrichment = 2.13, *p*-value = 1.40e-02, **Figure 5C**, top row, 2^nd^ column). Finally, we leveraged our KD analyses to study each of our proposed novel motif instances for RBFOX2 (**Supplemental Note S7**).

Notably, QKI demonstrated the most highly significantly enriched contexts, particularly in two cases that exhibit its positional effects on splicing regulation. First, QKI had significantly enriched contexts at the 3’SS of cassette exons in up-regulated DS events, signifying increased exon inclusion upon KD (enrichment = 9.18, *p*-value = 3.88e-44; **Figure 5C**, top row, 2^nd^ column). We also confirmed that our discovered QKI motif consensus CUAAC was enriched in the cassette exon 3’SS using both event-centric (enrichment = 1.65, *p*-value = 7.81e-08, **Supplemental Figure S16A**) and context-centric (enrichment = 2.28, *p*-value = 1.29e-04, **Supplemental Figure S16B**) methods of *k*-mer enrichment computation (**Materials and Methods**). Indeed, under normal conditions, QKI is known to bind upstream of cassette exons to promote exon skipping (Hall et al. 2013; Hinkle et al. 2022; Montañés-Agudo et al. 2023). Second, QKI had significantly enriched contexts at the 5’SS of cassette exons in down-regulated DS events, signifying increased exon skipping upon KD (enrichment = 14.07, *p*-value = 1.62e-60; **Figure 5C**, bottom row, 3^rd^ column), and the QKI consensus CUAAC was once again enriched in the cassette exon 5’SS (event-centric: enrichment = 2.24, *p*-value = 7.65e-16, **Supplemental Figure S16A**; context-centric: enrichment = 1.79, FDR = 1.94e-03, **Supplemental Figure S16B**). These findings corroborate QKI’s known functions in binding downstream of cassette exons to promote exon inclusion under normal conditions (Hall et al. 2013; Hinkle et al. 2022; Montañés-Agudo et al. 2023).

In closing, we demonstrate that our discovered contexts are not only enriched in regions implicated in splicing regulation, but also that the directional and positional context enrichment patterns are indicative of RBP functions in regulating expression and splicing, thus providing further validation for the results of our framework.

### Cross-prediction: a novel approach to investigate RBP binding specificity

The determinants governing RBP binding specificity, particularly for those RBPs sharing similar motifs and contexts, remains an important outstanding question. RBPs recognizing similar sequence patterns must be able to differentiate between candidate sites *in vivo* in order to maintain the degree of specificity necessary for efficient and accurate mRNA regulation. We asked whether we could simulate the biological process of binding site selection *in silico*, especially to determine whether our models are capable of capturing the binding specificity exhibited by RBPs that recognize similar motifs. To this end, we developed and implemented a novel strategy that we refer to as cross-prediction, whereby the model trained on one RBP’s contexts is presented with a distinct set of contexts belonging to another RBP and predicts the binding outcome for each context in that set (**Figure 6A**). The cross-prediction performance, an aggregated measure across all contexts in the dataset, serves as a proxy for how successfully an RBP, by way of its model, can distinguish between its cognate binding sites versus those belonging to another RBP. Because this approach implicitly compares the salient features learned by each RBPs’ model, it provides a measure of comparison between the binding patterns of different RBPs.

**Figure 6:**
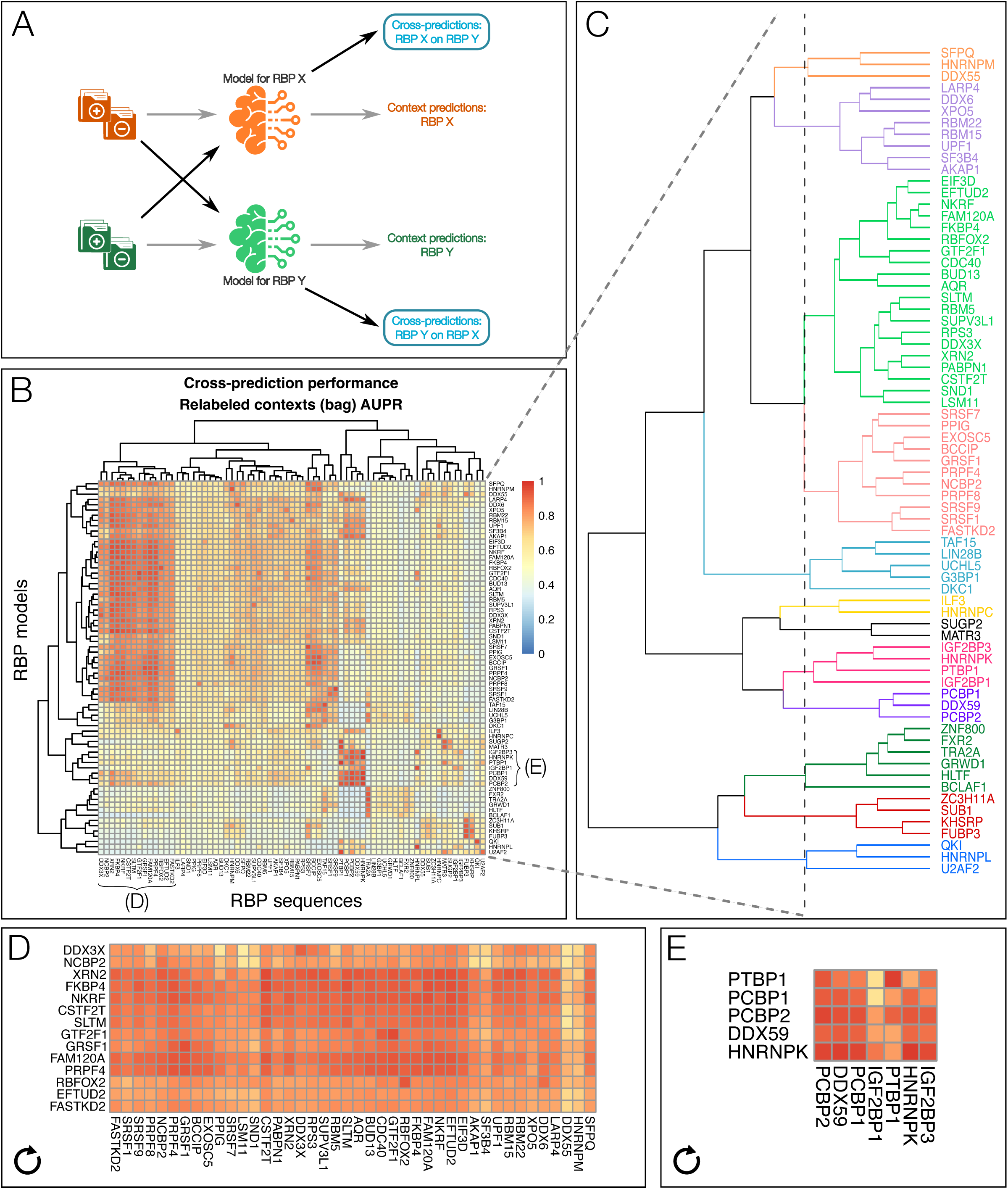
Cross-prediction as a novel approach to reveal similarities in global binding patterns between RBPs. (A) Schematic demonstrating the operations involved in the cross-prediction approach. (B) Clustered heatmap of the pairwise cross-prediction AUPRs between all RBPs. Regions highlighted in panels D and E are labeled in the heatmap. (C) Enlarged dendrogram from the clustered heatmap in panel B, revealing hierarchical clustering of RBPs based on similarity of their respective cross-prediction profiles across all other RBPs. (D-E) Highlighted regions of high cross-prediction performance from the heatmap in panel B; the arrow symbol in the bottom left corner indicates that the inset was rotated 90° clockwise relative to the original heatmap in B.

We hypothesized that a strong cross-prediction performance between a given pair of RBPs may suggest that the RBPs share certain salient features that are significant to their respective binding activities, including similarity in motifs, contexts, nucleotide preferences, and targets; it may also signify that the motif of one RBP is strongly enriched in the contexts of another RBP’s motif. At the global level, a high cross-prediction performance may indicate structural, behavioral, and functional similarity between the two involved RBPs. In cases where the cross-prediction between two RBPs is strongly asymmetric, it suggests that the RBP possessing the higher cross-prediction performance recognizes a larger percentage of the salient features present in the second RBP’s contexts than the other way around. We performed cross-prediction in a pairwise manner to comprehensively compare the binding patterns and activities of all RBPs. Only 316 pairs, or 6.36% of the total 4970 possible pairs (i.e., 71 choose 2), had a cross-prediction AUPR greater than 86.5% (i.e., the median cognate performance from above), indicating that the vast majority of RBP pairs demonstrate binding patterns that are distinct from one another (**Supplemental Table S12**). We subsequently generated a heatmap of cross-prediction performances, which provides an effective medium for visualizing the pairs of RBPs with strong cross-predictions (**Figure 6B**). We also conducted hierarchical clustering on the RBPs based on their respective cross-prediction profiles across the entire set of RBPs (**Figure 6C**), where assignment to the same cluster is suggestive of RBPs exhibiting similar global binding patterns to one another.

Of note, certain collections of RBPs demonstrated striking cross-prediction patterns (**Supplemental Note S8**), with the largest group (**Figure 6B**, top left; **Figure 6D**, inset) being most prominent: more than 50% (40 of 71 RBPs, excluding HNRNPM and DDX55) predicted relatively well on a set of 14 RBPs. This set of 14 comprised RBPs with strongly G-rich motifs and contexts, and different combinations of these RBPs clustered together by motif and context similarity measures (**Figures 3**, **4**). Upon further investigation, we noticed that each of the 14 RBPs have either exhibited binding of RNA G-quadruplexes (rG4s) (CSTF2T, DDX3X, FAM120A, GRSF1, GTF2F1, PRPF4), or showed binding enrichment at putative rG4s (EFTUD2, FASTKD2, FKBP4, NCBP2, NKRF, RBFOX2, SLTM, XRN2) in previous studies (Herdy et al. 2018; Lee et al. 2020; Busa et al. 2021; Dumas et al. 2021; Bourdon et al. 2023). The considerable degree of universality exhibited by cross-predictions on these 14 RBPs may indicate that their contexts contain highly similar and prevalent nucleotide patterns and/or structures; furthermore, the commonality between target genes for certain subsets of these RBPs (Lee et al. 2020) may suggest that these regions undergo coordinated regulation by a large number of factors, which might in turn reveal the importance of their target genes.

A group of RBPs (DDX59, HNRNPK, PCBP1, and PCBP2) with predominantly C-rich motifs and contexts (**Figure 3**) showed reciprocated, strong cross-prediction (**Figure 6E**, **Supplemental Table S12**). The lower cross-prediction by IGF2BP1/3 could be attributed to the more variable content of their motifs and contexts that did not exhibit a very strong preference for any given nucleotide. Importantly, PTBP1 also performed relatively well on the 4 strongly C-preferring RBPs (and vice versa) (**Supplemental Table S12**), suggesting that they together possess similar global binding patterns, despite PTBP1’s distinct, highly dissimilar U-rich motifs and contexts (**Figure 3**); PTBP1 did not cluster with any of the 4 C-preferring RBPs by motif and context KLD. This exemplifies that our cross-prediction results are informative regarding the identification of potential relationships between RBPs, as they can reveal what other analyses cannot.

Taken together, the novel cross-prediction strategy we introduce proves informative of the degree of similarity between the global recognition and binding patterns of RBPs. Importantly, we posit that a strong cross-prediction performance and/or membership to the same cross-prediction cluster may be suggestive of potential RBP-RBP interactions. While the clustering assignments observed by cross-prediction may differ with those produced by HDBSCAN (**Figure 3**) or KLD (**Figure 4**), each of the three measures provides an alternative viewpoint of the underlying data (i.e., pattern-based, Euclidean, and probabilistic, respectively). As we demonstrate below, integration of these features and the information they carry not only helps to provide a more complete depiction of the relationship between RBPs, but also helps to increase our confidence in interaction discovery – especially if the corresponding RBPs consistently cluster together across methods.

### A novel method to discover proximal interactions between RBP-RBP pairs

Given that RBPs frequently interact cooperatively or competitively to jointly regulate common targets (Dassi 2017), we developed a method to investigate RBP-RBP interactions. We first devised a conceptual delineation of the potential RBP-RBP interaction modes into concrete categories based on properties of the involved partners, including whether the RBPs bind proximally or distally to one another, whether one or both RBPs bind RNA, and whether the RBPs physically interact (**Figure 7A**; **Supplemental Note S9**). We used this classification to outline a scope within which we performed our RBP-RBP interaction discovery. Because our analyses are context-based, do not yet incorporate RNA secondary structure data, and do not yet account for long-range interactions across genomic features, we limited our method to consider only proximal interactions between pairs of RBPs (**Figure 7A**, left-hand major branch). We define 4 proximal interaction binding modes (**Supplemental Note S9**).

**Figure 7:**
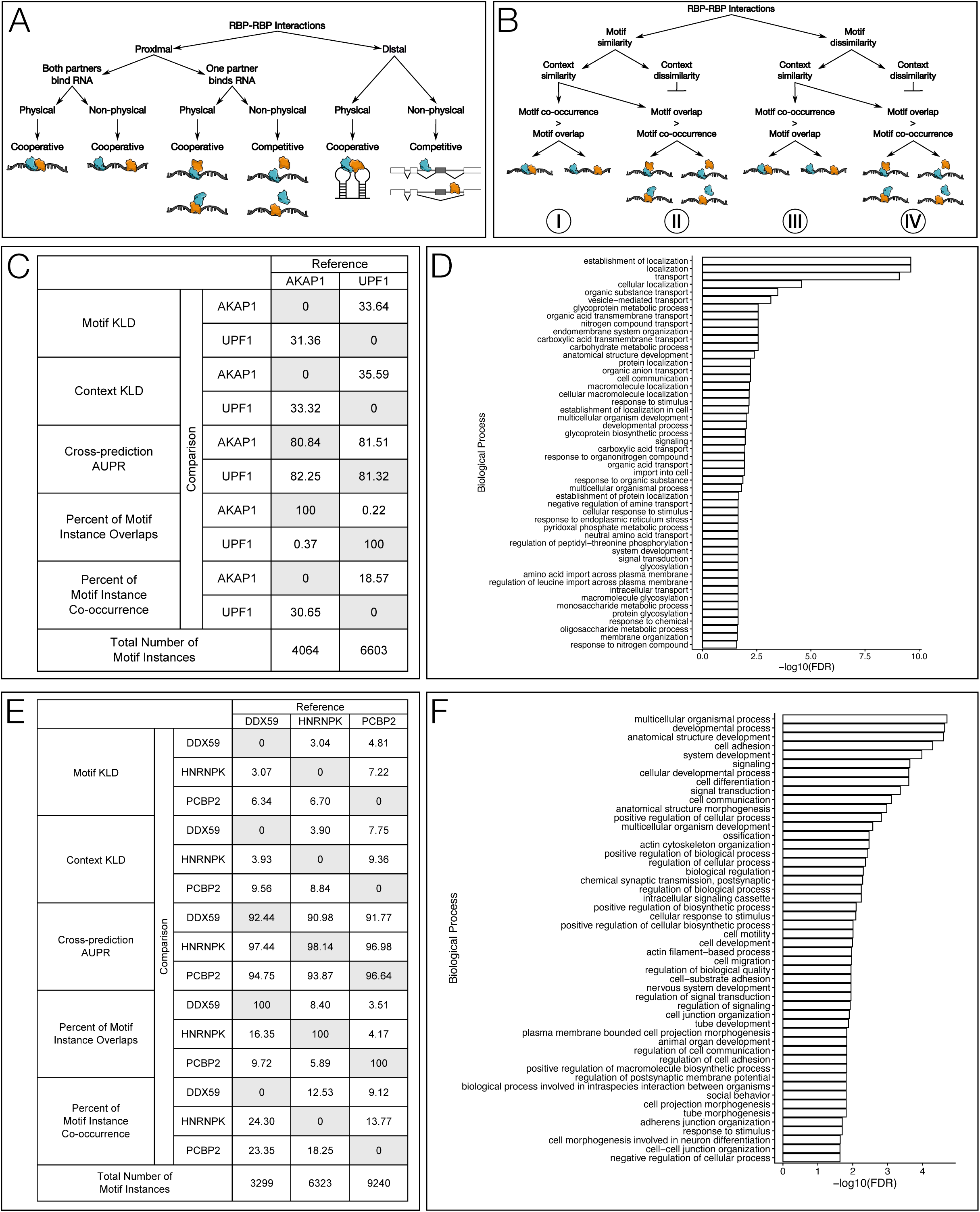
A method to infer RBP-RBP interactions. (A) Conceptual representation for the classification of RBP-RBP interaction categories, which depend on properties of the involved components. Distal interactions, while demonstrated here, are not considered in this analysis. (B) Interaction discovery method, represented as a decision tree based on features of the candidate RBP-RBP pair. Each path (I through IV) along the tree represents a set of possible interaction modes based upon the fulfilled conditions along the path. A candidate pair may be involved in a cooperative interaction, competitive interaction, or, potentially, both. (C) Feature summary and (D) GO analysis (top 50 terms with FDR < 0.05 shown) for the proposed interaction between AKAP1 and UPF1. (E) Feature summary and (F) GO analysis (top 50 terms with FDR < 0.05 shown) for the proposed interaction between DDX59, HNRNPK, and PCBP2.

Within this scope, we formulated an RBP-RBP interaction discovery method based on empirical observations and guided by reported interactions in the literature, which allowed us to define a set of characteristic features that we represent as a set of conditions in the form of a decision tree (**Figure 7B**). Each branch of the tree in turn represents a set of conditions that must be satisfied for a given candidate RBP-RBP pair to qualify as belonging to the corresponding interaction category (**Figure 7B**; **Supplemental Note S9**). The set of conditions used in our interaction discovery method comprise motif similarity, context similarity, motif co-occurrence, and motif overlap. As a strategy to reduce the search space of candidate interacting pairs, we began our investigation by referencing the clustering results from KLD and cross-prediction analyses, reasoning that RBPs assigned to the same cluster(s) may share similar motifs, contexts, and/or global binding profiles, and thus have a stronger likelihood of being potential interacting pairs, especially if the RBPs cluster together repeatedly across multiple metrics.

To validate the conditions of this method, we first examined known interacting RBP pairs established in the literature as positive controls (**Supplemental Table S13**). We started by examining pairs of RBPs with strong, reciprocated cross-prediction scores from **Figure 6E**. We noticed that HNRNPK and PCBP2 consistently demonstrated high similarity to one another, as witnessed by their respectably low motif and context KLDs (**Supplemental Table S13A**), membership to the same cluster in global KLD comparisons (**Figure 4**), and remarkably strong and symmetric cross-prediction performances (**Figure 6E**; **Supplemental Table S13A**). We discovered C-rich motifs and contexts for both of these RBPs (**Figure 3C**), which is supported by their belonging to the poly(rC) binding protein family (Choi et al. 2009). As heterogeneous ribonucleoproteins, their functional range is extensive and diverse, shuttling between the nucleus and cytoplasm to regulate pre-mRNA splicing, mRNA transport, mRNA stability, and mRNA translation (Kim et al. 2000; Lang et al. 2021). Experimental evidence has revealed an interaction between HNRNPK and PCBP2, with hypotheses that they may form an hnRNP complex to subsequently facilitate secondary interactions with regulatory effectors (Kim et al. 2000), but it remains unclear whether this interaction is cooperative or competitive, and whether one or both partners bind target mRNAs. Based on our finding that their motif co-occurrence substantially exceeds their motif overlap (**Supplemental Table S13A**), we believe that HNRNPK and PCBP2 have a higher propensity to form cooperative interactions in which both partners bind RNA (**Figure 7B**, Path I). For the less frequent cases in which their motifs overlap, this could be attributed to a cooperative mechanism in which the RBPs physically interact but only one partner binds RNA, or a competitive mechanism in which the RBPs compete for a binding site (**Figure 7B**, Path II). This case study serves as a validation of our method’s ability to detect an established RBP-RBP interaction with existing experimental evidence, and we additionally propose a novel hypothesis for their potential mode(s) of interaction.

With this positive control and others (**Supplemental Note S10**), we demonstrate that our proposed method successfully identifies known RBP-RBP interactions. These examples also demonstrate the combinatorics of RBP-RBP interactions (Khoroshkin et al. 2024), with certain RBPs (e.g., PCBP2, PTBP1, MATR3) being involved in multiple different interaction pairs, thus pointing to the exponential scale at which the full range of human RBPs interact. In total, we discovered 796 interactions with our method, which accounts for approximately 32% of the total 2,485 possible RBP-RBP pairs in our collection (**Supplemental Table S14**; **Supplemental Note S9**, **S11**; **Supplemental Figure S17**). As further validation, we cross-referenced our method with known, documented interactions in the BioGrid database (Oughtred et al. 2021) (**Supplemental Note S9**, **S11**). Of the 436 documented interacting RBP-RBP pairs that were present both in our collection and in BioGrid, 145 of these interactions (33.36%) were detected by our method, demonstrating our method’s ability and efficacy. We next sought to further investigate novel interacting pairs detected by our method and to hypothesize their potential functions and mechanisms.

#### NMD factor UPF1’s strong association with AKAP1 suggests its surveillant role in local translation at organelle membranes

We noted that AKAP1 and UPF1, despite visually appearing to have dissimilar motifs and nucleotide preferences (**Supplemental Table S5**), occurred in the same cross-prediction cluster (**Figure 6C**), which may be suggestive of a potential interaction. The relatively large divergence between their target motif logos (**Figure 7C**, “Motif KLD”), with AKAP1’s being predominantly C-rich (consensus: CUCCC) and UPF1’s being predominantly G-rich (consensus: GGGGG), indicated dissimilarity between the binding motifs of the 2 RBPs. Conversely, the binding contexts of the 2 RBPs exhibited some degree of similarity: AKAP1’s context logo appears C-rich but has G-rich undertones, while UPF1’s context logo is GC-rich. This context similarity is especially reflected in the fact that the context KLD value increased only slightly compared to the motif KLD value (from 31.36 for motif to 33.32 for context for AKAP1 vs. UPF1, and from 33.64 for motif to 35.59 for context for UPF1 vs. AKAP1; **Figure 7C**, “Context KLD”), despite being a cumulative sum across a considerably larger region of 55 nt compared to the target length of 5 nt (**Supplemental Note S9**, **S12**). Their relatively high and reciprocated cross-prediction scores (**Figure 7C**), as well as their membership in the same cross-prediction cluster (**Figure 6C**), indicate that the 2 RBPs likely share similar global binding patterns. Finally, as expected from their highly divergent motif PFMs, AKAP1’s (*N* = 3775) and UPF1’s (*N* = 6225) motif instances shared very little overlap (*N* = 14; **Figure 7C**, “Percent of Motif Instance Overlaps”); however, their respective co-occurrence rates were relatively high, with 30.65% of AKAP1’s overlapping UPF1’s and 18.57% of UPF1’s overlapping AKAP1’s (**Figure 7C**). Taken together, these features correspond with our conditions for cooperative interactions along Path III, and we therefore hypothesized that AKAP1 and UPF1 may work together in a synergistic manner to regulate common targets.

To further support our hypothesis, we demonstrate that overlapping AKAP1-UPF1 contexts have significantly higher eCLIP enrichment from both AKAP1 (**Supplemental Figure S18A**) and UPF1 (**Supplemental Figure S18B**) datasets than contexts belonging to either RBP that are not implicated in the interaction (**Materials and Methods**). We also demonstrate that overlapping AKAP1-UPF1 contexts were significantly enriched in the 3’UTRs of differentially expressed genes upon knockdown of both AKAP1 and UPF1 independently (**Supplemental Note S12**).

We sought to understand how this RBP pair may cooperate. We observed that the vast majority (93.3%) of overlapping AKAP1-UPF1 contexts were located in 3’UTRs, which are known to have roles in mRNA translation regulation (Mayr 2019). Both RBPs have documented functions in translation regulation: AKAP1 spatially regulates the local translation of essential mitochondrial proteins at the mitochondrial outer membrane (Gabrovsek et al. 2020; Sharma and Fazal 2024), while UPF1 is an essential RBP in the nonsense-mediated decay (NMD) pathway (Kim and Maquat 2019) (**Supplemental Note S12**). Upon examination of the gene set containing overlapping contexts, we not only found a number of genes encoding important mitochondrial proteins, but we also found that the gene set was significantly enriched in localization (e.g., protein localization), membrane (e.g., endomembrane system organization), and transport (e.g., vesicle-mediate transport) gene ontology (GO) terms (**Supplemental Note S12**), leading us to hypothesize that AKAP1’s functions of local protein synthesis may also occur at components of the endomembrane.

In light of these findings, we propose a cooperative mechanism between AKAP1 and UPF1 in which they work jointly, by way of Path III (**Figure 7B**), to ensure accurate local translation of specialized mRNA transcripts into full-length protein products at certain organelles. Our results suggest that UPF1 maintains a sustained interaction with AKAP1, whereby UPF1 is on vigilant standby to intervene in circumstances of local translation where AKAP1 has anchored mRNA transcripts containing premature termination codons to organelle membranes. This could present a quality control mechanism for the accurate translation and efficient use of proteins that are essential to organelle function, especially at the mitochondria and the endomembrane system, such that the resultant proteins become immediately available for insertion into the membrane or for import into the organelle. In summary, we infer novel subcellular localizations for AKAP1, and we propose a novel mechanism for the cooperative interaction between AKAP1 and UPF1.

#### The DEAD-box RNA helicase DDX59 facilitates the mRNA regulatory roles of HNRNPK and PCBP2

Interestingly, DDX59 demonstrated highly similar features with both HNRNPK and PCBP2 individually (**Figure 7E**), despite being a distinct RBP belonging to a large family of highly-conserved DEAD-box RNA helicases (Shamseldin et al. 2013; Linder and Jankowsky 2011). Of note, DDX59’s model demonstrated superior performance on the contexts of HNRNPK (AUPR = 97.44%) and PCBP2 (AUPR = 94.75%) compared to its own contexts (AUPR = 92.44%), suggesting a very strong degree of similarity in the underlying patterns of their respective binding regions. DDX59 also exhibited a higher degree of motif co-occurrence and motif overlap with HNRNPK and PCBP2, compared to those observed between the known interactors HNRNPK and PCBP2 (**Figure 7E**), which provides additional support for a potential interaction with DDX59. Although DEAD-box helicases interface with the RNA backbone of their targets in a non-sequence-specific manner (Linder and Jankowsky 2011; Jarmoskaite and Russell 2011), we discovered a C-rich nucleotide binding preference (**Figure 3C**) for DDX59, which may be a byproduct of frequent associations with poly(rC) binding proteins like HNRNPK and PCBP2 and subsequent co-immunoprecipitation. While the exact functions of DDX59 have not yet been deduced, its family members have annotated roles in ATP-dependent local RNA duplex unwinding, RNA clamping, and RNP assembly, stabilization, remodeling, and disassembly (Linder and Jankowsky 2011; Bohnsack et al. 2023). Based on our empirical evidence, we hypothesize that DDX59 interacts cooperatively with both HNRNPK and PCBP2 by any or all of the following possible mechanisms: (1) unwinding short regions of double-stranded RNA to enable binding by the single-stranded RNA-preferring KH domains of HNRNPK and PCBP2, (2) serving as an anchor point at which HNRNPK and PCBP2 can interact to form an hnRNP complex, (3) remodeling hnRNPs, and/or (4) disassembling hnRNPs. This hypothesis is further supported by the fact that HNRNPK has previously been shown to interact with a number of DEAD-box helicases to enact various regulatory processes (Chen et al. 2002; Li et al. 2019; Good et al. 2019; Chen et al. 2021).

Because each pair of RBPs within this trio exhibited strong features indicative of cooperative interactions, we leveraged transitive relations to hypothesize that DDX59, HNRNPK, and PCBP2 may also interact in unison to form a complex (**Supplemental Note S13**). To investigate this possibility, we performed a three-way intersection between their respective contexts (**Materials and Methods**) and found that 21.79% of DDX59’s contexts overlapped with both HNRNPK’s and PCBP2’s, with most of these events located in introns or coding sequences. From GO analysis of the genes containing context overlaps between all three RBPs, we discovered enrichment for anatomical structure development and its child term anatomical structure morphogenesis (**Figure 7F**); further inspection of additional child terms for both revealed many related to the development of the face (face, head, & craniofacial suture morphogenesis), oral cavity (hard & soft palate morphogenesis, roof of mouth development, tooth eruption), and digits (limb joint & appendage morphogenesis) (Binns et al. 2009). Mutations in DDX59 have been associated with orofaciodigital syndrome, a disease which leads to varied disfigurements in features of the mouth, face, and digits (Shamseldin et al. 2013; Faily et al. 2017), which may help to explain the discovered GO terms.

In conclusion, we not only hypothesize novel and specific roles for DDX59 in regulating mRNA processing via its strong association with either of both of the hnRNPs HNRNPK and PCBP2, but we also utilize combinatorics and transitive inference to propose an interaction between DDX59, HNRNPK, and PCBP2 – either as individual pairs, a complex of three proteins, or both – that holds mechanistic, functional, and disease significance, thereby demonstrating the utility of our approach.

## DISCUSSION

In this paper, we presented a novel, comprehensive computational pipeline to study the binding preferences, patterns, activities, functions, and interactions of RBPs. Inspired by state-of-the-art concepts from linguistics and NLP, we developed a sequence representation strategy that enabled us to clearly delineate RBP binding motifs and their contexts, and we subsequently leveraged Multiple Instance Learning to predict RBP binding of these contexts in a weakly supervised fashion. We also developed an accurate, consensus-based motif discovery algorithm to identify the binding and context motifs for 71 RBPs in the HepG2 cell line and 74 RBPs in the K562 cell line. Importantly, we discovered novel *in vivo* contexts for these RBPs, which is an important contribution to understanding the inherent nucleotide binding preferences exhibited by RBPs *in vivo* as well as their role in facilitating RBP binding specificity. With feature integration and transitive inference, we were able to leverage these findings and introduce a method for identifying known and novel RBP-RBP interactions, illustrating examples of each.

Our approach to RBP-RBP interaction discovery allowed us to identify many novel interacting pairs, and we were additionally able to hypothesize functional mechanisms pertaining to mRNA regulation for a subset of those pairs. In total, we characterized 2,485 RBP-RBP pairs (**Supplemental Table S14**; **Materials and Methods**). Based on our findings, the majority of detected interactions are likely cooperative, and this appears to align with previous work that has remarked on the biological and technical challenges involved in detecting competitive interactions (Lang et al. 2021). In addition, the mode of interaction between RBPs (Street et al. 2024) greatly influences its ability to be detected by our method, which for the time being addresses proximal, RNA-mediated interactions between pairs of RBPs. Long-range cooperative interactions in which RBPs bind distally from one another and are subsequently brought into closer proximity by virtue of RNA secondary structures may be more difficult to detect (Dassi 2017), as are long-range competitive interactions in which RBPs do not directly contend for the same binding site but instead bind distally from one another to promote divergent outcomes, such as in the proposed regulatory mechanism of alternative splicing in muscle differentiation between PTBP1 and QKI (Hall et al. 2013) (**Figure 7A**). For other binding modes, such as the cooperative and competitive modes in Path II (**Figure 7B**), we do not yet have the resolution to distinguish whether or not the feature set indicates a cooperative or competitive interaction, for which other types of data will be required. Though not explored in detail here, BCLAF1 and TRA2A present a strong example for the cooperative and/or competitive modes of Path II (**Supplemental Table S14**), and it would be of great interest to understand by which mode and for which functions they interact. We also do not consider homomeric interactions here. We additionally note that, due to its reliance on the percentage of motif instance and context overlaps, our approach is more attuned to detect RBP-RBP interactions that occur more frequently, which in turn implies that these interactions have more widespread regulatory roles on a larger set of mRNA targets. Indeed, the case studies we highlighted here (**Figure 7**) are those with very prominent features. On the other hand, less frequently occurring interactions that may govern the regulation of a small, specialized subset of mRNAs – which are still detected but have much lower overlap percentages – will require more detailed investigation in order to elucidate their precise functions.

By using the theory of transitive relation, we were also able to propose a novel *n*-way interaction (*n* > 2), as exemplified by our DDX59 case study. Importantly, this potential cooperativity between DDX59, HNRNPK, and PCBP2 has significant physiological and disease implications, thereby demonstrating the utility of our discovery. As additional support for this approach, we were also able to detect a putative interaction between RBPs recognizing G-rich motifs (namely, CSTF2T, FAM120A, GTF2F1, and PRPF4; **Supplemental Table S15A**), which has previously been identified (Lee et al. 2020) and therefore serves as a form of validation for our use of transitivity. We further propose that interactions may occur between other members of the 14 rG4-binding candidates isolated from our cross-prediction results (**Figure 6D**; **Supplemental Table S15B**); for example, the highly similar G-rich motif, preceded by a U and followed by a C, within a strongly G-rich context, belonging to ETUD2, FAM120A, GRSF1, and NKRF may be indicative of a potential interaction between the 4 RBPs. It is logical, therefore, to also extend this method in the future to infer larger macromolecular complexes. In fact, we were able to obtain preliminary results in this direction, identifying pairwise interactions between multiple components of the (pre-)catalytic spliceosome (**Supplemental Table S16**), with some RBPs being associated with either the U2, U5, or U4/U6 snRNPs, or the U4/U6 x U5 tri-snRNP complex (Sayers et al. 2022), and many having been captured together in existing Cryo-EM structures of the spliceosome (Szklarczyk et al. 2023). Two RBPs in this set, AQR and BUD13, have not, to the best of our knowledge, been reported in the literature, and we present them as a potential novel interacting pair within the spliceosome that is also supported by their discovered interactions with other spliceosomal proteins (**Supplemental Tables S14**, **S16**). We may even be able to obtain a finer resolution by using our feature integration and transitive inference to build up such complexes piecewise based on progressive degrees of similarity. To facilitate detections of these large-scale interactions, the cross-prediction and KLD dendrograms may prove helpful, as their inherent hierarchical nature may provide clues regarding the combinatorics, hierarchy, and order in which RBP-RBP interactions are formed. We also note that, while we used the KLD and cross-prediction clusters to guide our identification of candidate RBP pairs, this does not preclude the possibility that interactions also occur between RBPs from different clusters; such cases, which will also contribute to the discovery of larger complexes, require further examination. Overall, our results provide a promising advancement and outlook for charting the vast landscape of RBP-RBP interactions.

Finally, future avenues could entail analyses from numerous angles. Firstly, additional studies will be needed to investigate the proposed mechanisms of novel interactions discussed in this paper. The many interactions we identified but did not elaborate on here (**Supplemental Table S14**) will also require additional exploration to understand their underlying mechanisms, functions, and scale. We are also interested in extending our analyses to encompass mutual interactions, which we were able to preview in our AKAP1-UPF1 case study: the gene set containing context overlaps between the 2 RBPs contains the UPF1 gene itself, indicating that (1) UPF1 regulates the translation of its own cognate mRNA, a process referred to as an autogenous interaction, and (2) AKAP1 is involved in the regulation of UPF1’s mRNA, a process referred to as a heterogeneous interaction (Dassi 2017). Solely within our collection of RBPs, our preliminary results reveal a strong prevalence of mutual interactions: ∼97% (69 of the 71) are involved in any type of mutual interaction, while 100% of those detected mutual interactions demonstrated heterogeneous interactions and ∼30% (*N* = 21) also demonstrating autogenous interactions. The full extent of the mutual interaction landscape will become clearer as these analyses are expanded to encompass the entire set of human RBPs, as will their functional consequences and implications. Other biological questions of interest include (1) to what extent RBP binding patterns and RBP-RBP interactions exhibit cell type specificity, (2) how RNA secondary structures contribute to enable specific RBP binding and distal RBP-RBP interactions, (3) how RBP binding preferences are influenced by cooperative and competitive interactions with microRNAs (Ciafrè and Galardi 2013; Kim et al. 2021), (4) how RBP activities, functions, and interactions can be further elucidated via RBP regulatory network modeling, and (5) how the discovered RBP binding patterns and interactions play a role in disease mechanisms. In conclusion, the method we have presented herein serves as a valuable contribution to the field’s current knowledge of RBP binding patterns, context preferences, and interactions, all enabled by the advent of state-of-the-art machine learning, deep learning, and modeling techniques; we believe our work sets an important foundation for future studies in these areas.

## MATERIALS AND METHODS

### Comparative analyses of eCLIP read enrichment over discovered motifs and contexts

As further validation, we performed comparative analyses to demonstrate that our discovered motifs and contexts are significantly more enriched in eCLIP read coverage compared to a set of matched control regions. To define a set of comparable control regions, for every motif instance in our discovered motif, we randomly sampled a genomic instance of the same *k*-mer that met the following conditions: (1) belongs to a gene expressed in the same cell line (i.e., HepG2), (2) belongs to the same chromosome as the motif instance, (3) belongs to the same genomic region as the motif instance, (4) when extended symmetrically to the same length as the discovered contexts (i.e., 55 nt), does not overlap with any discovered contexts.

We next downloaded the eCLIP read alignment .bam files from the ENCODE portal for all RBPs and their corresponding size-matched input experiments (**Supplemental Table S1C**). For each RBP, we performed the following procedure for both (1) the discovered motifs and contexts and (2) the matching control “motifs” and “contexts”, based on previous eCLIP studies (Van Nostrand et al. 2016, 2020a, 2020b). We first used bedtools coverage (Quinlan and Hall 2010) to determine the number of eCLIP reads overlapping each region (normalized by read depth of the sequencing experiment) and averaged across both replicates; we also computed normalized eCLIP read coverage overlapping each region for the single corresponding size-matched input (SMInput) experiment.

Next, we computed each region’s eCLIP enrichment by:

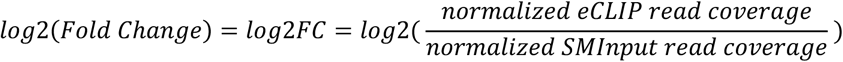

To compute the significance of each region’s enrichment, we ran a chi-square test if the raw read count overlapping the region was greater than 5 reads for both the eCLIP and input experiments, and ran the Fisher’s exact test otherwise. Because each eCLIP replicate had a different raw read count, we used Fisher’s method to combine the resulting p-values from both replicates (Fisher 1973). Finally, we performed multiple test correction across all regions for a given RBP by computing the FDR. We designate a region as significantly enriched in eCLIP if the log2FC is greater than 0 and the FDR is less than 0.05.

To compare whether discovered motifs had a significantly greater eCLIP enrichment than their matched controls, we used a Wilcoxon rank-sum test, then visualized the corresponding distributions as nested violin-box plots. We performed the same for the comparison between discovered contexts and their matched controls.

We also leveraged the eCLIP enrichments to support our discovered RBP-RBP interactions, following a similar procedure to the one above. For a given RBP-RBP pair, we computed the eCLIP enrichment independently for both RBPs implicated in the interaction, over four sets of regions: (1) contexts belonging to RBP #1 that are not implicated in the interaction (i.e., don’t overlap with contexts belonging to RBP#2), (2) contexts belonging to RBP#1 that are implicated in the interaction (i.e., overlap with contexts belonging to RBP#2), (3) contexts belonging to RBP#2 that are implicated in the interaction, and (4) contexts belonging to RBP#2 that are not implicated in the interaction. We again used the Wilcoxon rank-sum test to determine whether the predicted interaction’s binding sites (containing overlapping or co-occurring motifs) demonstrated significantly stronger eCLIP enrichment than each RBP’s non-interacting binding sites.

### Quantification of motif discovery accuracy

We quantified the accuracy of our motif discovery algorithm by curating a set of ground-truth motifs for RBPs with strongly characterized binding patterns in the literature, which we define as having specified logos, consensus sequences, or nucleotide preferences that are concordant across multiple sources. This resulted in a set of 14 RBPs each for HepG2 and K562. For this evaluation, we opted against performing a cross-reference analysis using tools that compare user-provided motifs with those from existing databases in order to determine the identity of the proteins to which the motifs belong (Bailey et al. 2015); while such an approach is certainly useful in specific circumstances, we deemed it unsuitable for our purposes because (1) most available databases are transcription factor-centric, (2) the available RBP databases are primarily derived from *in vitro* data only, (3) the available RBP databases are limited in the number of RBPs and motifs they contain with respect to the full scope of literature available, and (4) the outcomes of such analyses may be misleading, as they do not account for RBPs which may recognize different binding motifs depending on which RBD is interacting with its target mRNA at the time of experimental capture.

### Calculation of sequence and context *k*-mer enrichments

To compute *k*-mer enrichments, we performed the following procedure. For sequence-level *k*-mer enrichments, for each unique *k*-mer, we computed the ratio between the proportion of positively-predicted and negatively-predicted sequences (from the baseline model) containing the *k*-mer. For context-level *k*-mer enrichments, for each unique *k*-mer, we computed the ratio between the frequency of the *k*-mer as the target of positively-predicted contexts and the frequency of the *k*-mer as the target of negatively-predicted contexts (from the contexts model). For each, we used the associated counts to perform chi-square testing and obtain a *p*-value for each *k*-mer indicating the significance of its enrichment. We considered a *k*-mer enriched if it had an enrichment greater than 1 and a p-value less than 0.05.

### Random motif discovery

To generate a set of random genomic “peaks”, we implemented the following procedure. For each sampled sequence, we (1) randomly selected a chromosome from {chr1-chr22, chrX}, (2) randomly selected a genomic region from {CDS, 3’UTR, 5’UTR, 5’SS, 3’SS, proximal intron, distal intron} from the chromosome selected in step 1, (3) randomly selected a position within the region selected in step 2 to serve as the central position, and (4) extended +/-50 nt around the central position selected in step 3. We repeated this procedure 10,000 times, making sure that each sampled sequence did not overlap eCLIP peaks from any RBP from neither the HepG2 nor K562 datasets. The final set, containing 10,000 random sequences of length 101 nt, served as our set of random “positive” sequences.

To generate the set of corresponding “negative” sequences, we performed the same procedure we used to derive the set of negative sequences for training our models (**Supplemental Note S1**). Briefly, for each “peak” in our positive set, we randomly sampled a sequence of the same length from a matched chromosome and genomic region, and resampled as needed to ensure that no sequence overlapped with the “positive” set or with any eCLIP peaks. The final set served as our set of random “negative” sequences, leading to a total random dataset size of 20,000 sequences. For comparison, we used the same procedure to generate datasets of size 2,000 and 10,000.

To generate a set of random synthetic “peaks”, we generated random sequences of length 101 nt by sampling from the alphabet {A, C, G, T} under a uniform distribution. We similarly generated sets of 2,000, 10,000 and 20,000 sequences, and performed the same analyses on each (i.e., motif discovery with our algorithm and with STREME).

For each of the 6 random datasets we generated, we first deconstructed the set of random sequences into contexts using our sequence decomposition method, then ran our motif discovery algorithm on the resulting contexts. For comparison, we also submitted the contexts to STREME.

To contrast the properties of the results from random sequences (i.e., prediction AUPR, consensus/*k*-mer enrichments and *p*-values, etc.) with those of RBPs, we selected a subset of 7 sequence-specific RBPs (HNRNPC, HNRNPK, MATR3, PTBP1, QKI, RBFOX2, SRSF1) as well as 6 non-sequence-specific RBPs (DDX3X, DDX55, DDX59, DDX6, ILF3, XPO5) from our work in the HepG2 cell line for comparison.

### Determination of motif and context nucleotide preference and symmetry

We determined the nucleotide preference of each RBP’s motif and context by examining its PFM representation. Starting with the left context, we identified the most prevalently occurring nucleotide for every position, then ranked each of the four nucleotides in descending order based on the number of positions at which each occurred most prevalently. We also computed the ratio of the number of positions for each nucleotide relative to the maximum of these values. To account for cases in which RBPs demonstrate a clear preference for more than one nucleotide (e.g., HNRNPL), we deemed any nucleotide for which its position count was at least half that of the top-ranking nucleotide as also demonstrating a strong preference. We performed these operations for the left context, motif, and right context individually to obtain nucleotide preferences at a finer resolution, and finally performed it for the full context (excluding the motif) to determine the overall nucleotide preference of the context. We deemed a context symmetric if its preferred nucleotide(s) in the left and right contexts were the same. Similarly, we compared the preferred nucleotide(s) in the motif and the full context to determine whether they shared the same preferences.

### Computation of cumulative motif and context Kullback-Leibler Divergence (KLD)

The Kullback-Leibler Divergence (KLD) is a statistical measure that quantifies the degree to which one probability distribution *Q* (i.e., the approximating distribution) is different from another probability distribution *P* (i.e., the reference distribution). In information theory, the KLD is also known as the relative entropy, where smaller values indicate that *Q* is a closer approximation of *P*, and the minimum value of 0 indicates that the two probability distributions are identical (Bishop and Nasrabadi 2006). The KLD does not qualify as a metric in the formal mathematical sense because it does not satisfy two main properties that are required of a metric. First, the KLD is asymmetric, such that KLD(*P*, *Q*) does not necessarily equal KLD(*Q*, *P*). Second, the KLD does not satisfy the triangle inequality: if two approximating distributions *Q* and *S* are each independently similar to the true distribution *P*, it does not necessarily mean that *Q* and *S* are similar to one another. Thus, we use KLD as a statistical measure of distribution similarity.

In our application, we reasoned that the position probability matrix (PPM), which we use to represent both motifs and contexts, fundamentally comprises a series of position-specific probability distributions across the 4 canonical nucleotides, where each position has its own distribution. For a given position in a PPM, we can use the KLD to approximate the similarity between the corresponding nucleotide probability distributions at that position for any given pair of RBPs. For a single position within any given sequence, the KLD is computed as:

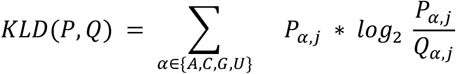

Note that the KLD has some finite maximum value that is dependent on each distribution, provided that no values in the distribution are equal to 0.

Because we sought to compare the distributions across the full length of the motifs and contexts, we herein introduce the cumulative KLD, which sums together the position-specific KLDs for the entire motif and context, respectively:

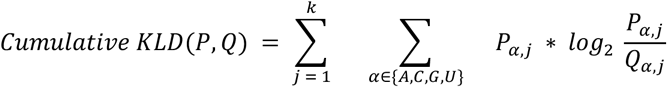

where *k* = 5 for motifs and *k* = 55 for the full contexts. Thus, the values that we present in this work are cumulative KLDs that represent the sum of position-specific KLDs across an entire PPM.

### Processing of RBP knockdown shRNA-seq data

#### Differential expression analyses

To functionally connect our motifs and contexts to the regulation of gene expression, we ran DESeq2 (Love et al. 2014) on the shRNA-seq raw count data, defining any gene whose expression in the KD dataset was 1.5 times greater or less than the expression in the corresponding control dataset (i.e., log2FC > log2(1.5)) and whose corresponding *p*-value and FDR were both less than 0.05. We also defined the set of expressed genes in any experiment as those whose average TPM across 2 replicates was greater than 0 in at least one of the control or KD conditions. For each RBP, we designated the set of differentially expressed (DE) genes as the foreground set, and the remaining expressed but not DE genes as the background (bkgd) set.

We then computed the extent to which our discovered contexts were enriched in DE genes relative to background, based on the general formula:

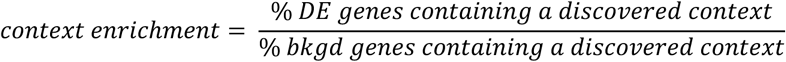

To more closely examine the functional effects of RBP KD, we reasoned that this general metric required stratification on two levels in order to prevent information loss: (1) based on whether or not the DE gene was up-or down-regulated, and (2) based on the genomic region in which the RBP’s context is located Therefore, we computed a directional, region-based context enrichment as follows:

For direction *d*, s.t. *d* ∈ {*up, down*}, and region *r*,, s.t. *r*, ∈ {5′ *UTR*, *CDS*, 3′*UTR*}:

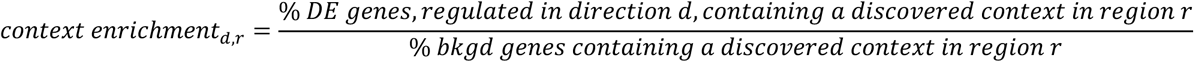

#### Differential splicing analyses

To functionally connect our discovered motifs and contexts to the regulation of alternative splicing, we ran rMATS-turbo (Wang et al. 2024) on the shRNA-seq bam files (**Supplemental Table S1D**). We focused primarily on the skipped exon (SE) class of events, as they were the most prevalent event class, representing approximately 70% of all detected events for each RBP (**Supplemental Figure S12A**). We filtered detected SE events using the following criteria: (1) events must have a replicate-averaged PSI value between 0.05 and 0.95 in at least one of the control or knockdown conditions (Dominguez et al. 2018; Van Nostrand et al. 2020a), and (2) events must have a minimum total inclusion + exclusion junction read coverage of 10 (Dominguez et al. 2018). This filtering yielded approximately 13,439 SE events (on average) across all RBPs, representing 32.78% (on average) of all detected SE events across all RBPs (**Supplemental Figure S12B**).

We defined an event as being differentially spliced (DS) if its deltaPSI between control and KD conditions was greater than 0.05, and both its *p*-value and FDR were less than 0.05 (Van Nostrand et al. 2020). We defined all remaining SE events as background (bkgd) events. The number of DS events demonstrated more variability across RBP KD experiments, with approximately 1,029 DS events (on average) per RBP, representing 7.31% (on average) of all filtered SE events per RBP (**Supplemental Figure S13A**, **S13B**). The majority of RBPs demonstrated a preference for up– or down-regulated events upon knockdown, which provides insight into their regulatory effects on exon inclusion or exon skipping under normal conditions (**Supplemental Figure S13C**). For enrichment analyses, we defined the set of DS events as our foreground and all other filtered SE events as our background.

To determine whether our discovered motifs and contexts are enriched in these events, we first defined regions of interest that have been established to contain sequence elements for splicing factors to bind. We defined 4 regions of interest that are typically implicated in the splicing of cassette exons (adapted from Yee et al. 2018) at (1) the 3’ end of the upstream exon [“upstream”], (2) the 3’SS of the cassette exon [“cassette 3’SS”], (3) the 5’SS of the cassette exon [“cassette 5’SS”], and (4) the 5’ end of the downstream exon [“downstream”] (**Figure 5B**). Each of the 4 regions span 50 nucleotides (nt) into the associated exon and 300 nt into the associated intron, except for cases that fall within the following exceptions: (1) if the upstream or downstream exons are less than 50 nt long, the region only extends to the exon boundary, (2) if the cassette exon is shorter than 100 nt long, the region only extends halfway into the exon, and (3) if either of the introns are shorter than 600 nt long, the region only extends halfway into the intron.

Similarly to the computation of context enrichments in our DE analyses, we also computed the extent to which our discovered contexts were enriched in DS SE events relative to background, based on the general formula:

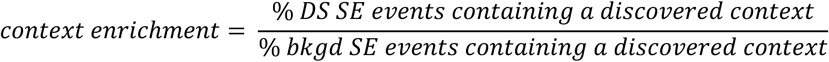

It thus follows that the directional, region-based context enrichment is computed as:

For direction *d*, s.t. *d ∈*{*up, down*}, and region *r*, s.t. *r ∈*{*upstream*, *cassette5prime*, *cassette3prime*, *downstream*}:

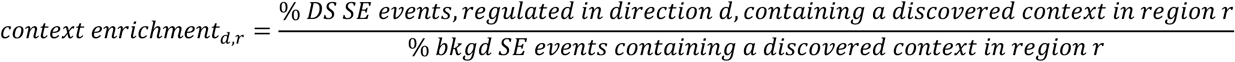

We computed *p*-values and visualized our results in a similar manner to the differential expression analyses.

We used bedtools getfasta (Quinlan and Hall 2010) to fetch the genomic sequences corresponding to all regions of all events. We computed *k*-mer enrichments using one of two methods. In the first method, which is event-centric, we computed the enrichment of *k*-mer *i* in region *r*, s.t. *r*, ∈ {*upstream*, *cassette5prime*, *cassette3prime*, *downstream*}, in DS events regulated in direction *d*, s.t. *d* ∈ {*up, down*}, as follows:

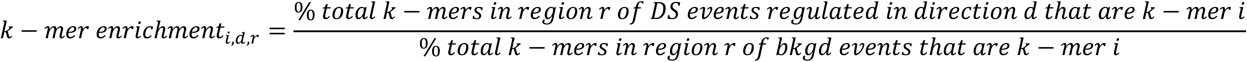

In the second method, which is context-centric, we computed the enrichment of *k*-mer *i* using our original starting context dataset. We first used bedtools intersect (Quinlan and Hall 2010) to isolate any contexts that overlap either DS or background events, then defined our foreground as contexts overlapping DS events and our background as contexts overlapping background events. For any context overlapping both a DS event and a background event, we considered that context as part of the background and removed it from the foreground. We then computed the context-centric enrichment for *k*-mer *i* in region *r*, s.t. *r*, ∈ {*upstream*, *cassette5prime*, *cassette3prime*, *downstream*}, in DS events regulated in direction *d*, s.t. *d* ∈ {*up, down*}, as follows:

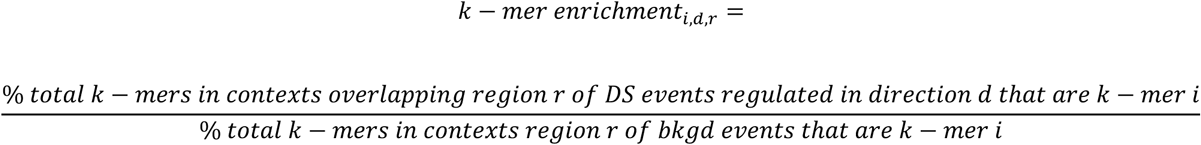

For both methods of *k*-mer enrichment computation, we evaluated statistical significance and visualized results as previously described.

### Context intersections for DDX59, HNRNPK, and PCBP2

To identify distinct interactions between contexts belonging to any pair of RBPs within this trio, we first performed Bedtools intersect (Quinlan and Hall 2010) between the contexts of two RBPs (e.g., HNRNPK and PCBP2), then performed Bedtools intersect again, this time with the contexts of the third RBP (e.g., DDX59) using the –v option; this produced the set of contexts from HNRNPK and PCBP2 that overlapped with each other but that simultaneously did not overlap with DDX59’s contexts. We repeated this procedure for the other two pairs (HNRNPK and DDX59; PCBP2 and DDX59).

To identify all overlaps between contexts belonging to each of the three RBPs, we sought to account for all possible combinations of orderings (with respect to genomic coordinates) in which these overlaps could potentially occur. For example, three contexts – one from each RBP – may all overlap one another simultaneously. In other cases, the three contexts may sequentially (with respect to genomic coordinates) overlap each other in a staggered manner, such that there is continuity between all three, but they do not all overlap with one other; the order in which the three can overlap in such a manner is combinatorial. We first performed intersections with Bedtools (Quinlan and Hall 2010) between two RBPs (e.g., HNRNPK and PCBP2) to identify their set of overlapping contexts. Then, we intersected the third RBP’s contexts (e.g., DDX59) independently with the subset of HNRNPK’s contexts that overlapped PCBP2’s, and with the subset of PCBP2’s contexts that overlapped HNRNPK’s; note that both of these operations are required to capture all possible interactions with DDX59. We merged the output that resulted from these last two intersections and filtered out duplicates to arrive at the subset of DDX59’s contexts that overlap both HNRNPK’s and PCBP2’s contexts. Finally, to account for the combinatorics of the possible order of overlaps, we repeated this multi-step intersection procedure two more times: once where we first intersected HNRNPK’s and DDX59’s contexts, then with PCBP2’s contexts; and once where we first intersected PCBP2’s and DDX59’s contexts, then with HNRNPK’s contexts. Note that all intersections were performed in a strand-specific manner.

### Gene Ontology (GO) analysis

In accordance with the pipeline used to process eCLIP data at the ENCODE portal (Luo et al. 2020), we used the GENCODE v29 annotation (Harrow et al. 2012) for all gene-level analyses. For a given RBP-RBP pair, we used Bedtools (Quinlan and Hall 2010) to intersect the genomic coordinates of their motif instances and co-occurrences with the GENCODE annotation to determine in which genes they were located; the resulting unique set of genes served as our test set for GO analysis. To derive a background set, we used all genes that were expressed in an ENCODE poly(A) RNA-seq data set from the same HepG2 cell line (experiment accession: ENCSR000CPE; file accession: ENCFF168QKW; TPM threshold = 1). We used the gProfiler package (Kolberg et al. 2023) to perform GO analysis of biological processes, using FDR < 0.05 as a cutoff to identify significantly enriched GO terms.

## DATA AVAILABILITY

Data is publicly available at the ENCODE portal. Accession for eCLIP data used in this study are detailed in **Supplemental Table S1**.

## CODE AVAILABILITY

Code is publicly available at https://github.com/orbitalse/novel_RBP_discovery_methods.

## AUTHOR CONTRIBUTIONS

S.I.E.: Problem Formulation, Conceptualization, Data Curation, Data Processing, Investigation, Formal Analysis, Methodology, Visualization, Writing of Manuscript (original draft), Review of Manuscript, Editing of Manuscript. Z.W.: Research Area, Discussions, Review of Manuscript. All authors reviewed and approved the final version of the manuscript.

## ACKNOWLEDGEMENTS

We thank Dr. Athma A. Pai for critical reading of and feedback on the manuscript.

## FIGURE CAPTIONS

**Supplemental Figure S1: Overview of the NLP representation, MIL formulation, and implementation of the RBP binding prediction task**. (A) Sequence definition and Multiple Instance Learning formulation. Positive and negative observations, derived from matching genomic regions, are deconstructed into their constituent contexts with the novel sequence decomposition method (B-C), then organized into bags. The precise RBP binding region within the positive sequence is highlighted in green and represents the main prediction objective, indicating the specific contexts that confer upon the parent sequence its positive label. (B) The first phase of the sliding window-based sequence decomposition strategy deconstructs a sequence into candidate target *k*-mers. (C) The second phase of sequence decomposition constructs the context elements for each candidate target generated in panel B. (D) Process diagram of the model training and iterative relabeling pipeline.

**Supplemental Figure S2: A novel algorithm for motif and context discovery.** (A) Phase I: Identification of candidate consensuses. (B) Phase II: Identification of candidate motif instances, using position-dependent similar *k*-mers. (C) Phase III: Identification of motif instances with *k*-mer co-occurrence. (D) Phase IV: Motif construction. (E) Phase VI: Context discovery.

**Supplemental Figure S3: A closer inspection of the PABPN1 motif and contexts in HepG2.** (A) Motif (top) and context (bottom) logos for PABPN1 in HepG2. (B) Top 10 motifs discovered by STREME; the known PABPN1 poly(A) motif is absent. The 7th-ranked motif corresponds to our discovered GCUGG motif. (C) Enrichment of eCLIP reads, relative to size-matched input, at discovered motifs and contexts compared to corresponding random control regions. Wilcoxon rank-sum test *p*-values test whether the motifs or contexts have significantly eCLIP enrichment than their corresponding control regions.

**Supplemental Figure S4: A closer inspection of the RBFOX2 motif and contexts in K562.** (A) Motif (top) and context (bottom) logos for RBFOX2 in K562. (B) Top 10 motifs discovered by STREME. (C) Enrichment of eCLIP reads, relative to size-matched input control, at discovered motifs and contexts compared to corresponding random control regions. Wilcoxon rank-sum test *p*-values test whether the motifs or contexts have significantly eCLIP enrichment than their corresponding control regions.

**Supplemental Figure S5: Motifs discovered from (A) iCLIP, (B) HITS-CLIP, and (C) PAR-CLIP assays**. Datasets for each RBP were downloaded from the following studies. RBFOX2 (H9): (Wu et al. 2016); LSM6 (HEK293): (Nabeel-Shah et al. 2024); EIF4G1 (K562): (ENCODE Project Consortium et al. 2020); QKI (mesHMLE): (Pillman et al. 2018); SRSF1 (H1299): (Fish et al. 2019); MSI2 (K562): (Park et al. 2014); ELAVL1 (T24): (Chen et al. 2019); HNRNPA1 (HEK293T): (Levengood et al. 2024); IGF2BP1 (HepG2): (Liu et al. 2024).

**Supplemental Figure S6: Motif discovery for random genomic sequences.** (A) Discovered motif, context, and nucleotide preferences for the random dataset comprising 20,000 total sequences (10,000 “positive” and 10,000 “negative”) derived from the human genome. (B) Top 10 motif hits returned by STREME. (C) Background nucleotide distribution for all 10K “positive” sequences (leftmost column), followed by the background nucleotide distribution for the same 10K sequences stratified by genomic region of origin (remaining 7 columns). The percentage of each genomic region in the dataset is annotated below each region. The context nucleotide preference for As and Us (panel A) reflect the background nucleotide preferences of the positive sequences.

**Supplemental Figure S7: Motif discovery for random synthetic sequences.** (A) Discovered motif, context, and nucleotide preferences from the random dataset comprising 20,000 total sequences (10,000 “positive” and 10,000 “negative”) derived from a uniform distribution of each of the 4 canonical nucleotides. There is no nucleotide preference in the contexts, thus reflecting the underlying uniform distribution. (B) Top 10 motif hits returned by STREME.

**Supplemental Figure S8: Properties of random motif discovery.** (A) Precision-recall (PR) curves for a select subset of 7 sequence-specific RBPs, 3 sets of random genomic sequences of total sizes 2K, 10K, and 20K sequences, and 3 sets of random synthetic sequences of total sizes 2K, 10K, and 20K sequences. (B) PR curves for a set of 6 non-sequence-specific RBPs, 3 sets of random genomic sequences, and 3 sets of random synthetic sequences. (C) Distribution of the enrichments (log2 scale) for the motif consensuses of all RBPs, the select subset of sequence-specific RBPs, non-sequence-specific RBPs, and the random genomic and random synthetic sets. (D) Distribution of the *p*-values (−log10 scale) for the motif consensuses of the same datasets as in panel C.

**Supplemental Figure S9: Properties of random motif discovery, continued.** (A) Distribution of the enrichments for all significantly enriched *k*-mers in the select subset of 7 sequence-specific RBPs and the random genomic and random synthetic sets. (B) Distribution of the *p*-values for all significantly enriched *k*-mers in the same datasets as in panel A. (C) Distribution of the enrichments for all significantly enriched *k*-mers in the subset of 6 non-sequence-specific RBPs and the random genomic and random synthetic sets. (D) Distribution of the *p*-values for all significantly enriched *k*-mers in the same datasets as in panel C.

**Supplemental Figure S10: Investigation of motif clusters.** (A) Distribution of the number of motif instances per sequence for HNRNPK. (B) Positional distribution of motif instances across the full length of the 55nt context sequences, displaying the percent of sequences with a motif instance at each position for HNRNPK. (C) Example sequence with motif instances forming a cluster. (D) Distribution of the number of motif instances for RBFOX2. (E) Positional distribution of motif instances across the full length of the 55nt context sequences, displaying the percent of sequences with a motif instance at each position for RBFOX2.

**Supplemental Figure S11: Interpretation of KLD values.** (A) Correlation between cross-prediction scores and context KLD values for each RBP-RBP pair that satisfies the cross-prediction conditions defined for interaction discovery. (B) Distribution of context KLD values, stratified by ranges of cross-prediction scores. (C) Histogram of context KLD values for RBP-RBP pairs that satisfy the cross-prediction conditions defined for interaction discovery.

**Supplemental Figure S12: Breakdown of contexts by genomic region.** (A) Breakdown of all contexts (left) and contexts in DE genes (right) by genomic region for UPF1. (B) Breakdown of all contexts (left) and contexts in DE genes (right) by genomic region for CSTF2T. (C) Breakdown of all contexts (left) and contexts in DE genes (right) by genomic region for IGF2BP3.

**Supplemental Figure S13: Breakdown of detected splicing events across all 60 RBPs in our HepG2 dataset that have shRNA-seq knockdown (KD) data**. (A) Barplot of the number of splicing events detected by rMATS for each RBP, stratified by event type. Skipped exon (SE) events represent approximately 70% of events across RBPs. (B) Barplot of the number of SE splicing events that passed filtering criteria (**Materials and Methods**).

**Supplemental Figure S14: Breakdown of detected differentially spliced (DS) skipped exon (SE) events across all 60 RBPs in our HepG2 dataset that have shRNA-seq knockdown (KD) data.** (A) Barplot of the number of differentially spliced (DS) skipped exon (SE) events detected for each RBP. (B) Barplot of the percent of filtered SE events (**Supplemental Figure S12B**) that are differentially spliced. (C) Breakdown of the percentages of directional differential splicing regulation for each RBP. Up-regulated signifies a positive deltaPSI and increased exon inclusion, while down-regulated signifies a negative deltaPSI and decreased exon inclusion (i.e., increased exon skipping).

**Supplemental Figure S15: *k*-mer enrichment in differentially spliced (DS) skipped exon (SE) events for RBFOX2.** (A) Event-centric *k*-mer enrichment. All enriched and depleted *k*-mers are labeled. (B) Context-centric *k*-mer enrichment. Only enriched and depleted RBFOX2 motif instances are labeled.

**Supplemental Figure S16: *k*-mer enrichment in differentially spliced (DS) skipped exon (SE) events for QKI.** (A) Event-centric *k*-mer enrichment. All enriched and depleted *k*-mers are labeled. (B) Context-centric *k*-mer enrichment. Only enriched and depleted QKI motif instances are labeled.

**Supplemental Figure S17: Inspection of the peak overlap-centric method for RBP-RBP interaction discovery**. (A) Comparison of the distribution of bidirectional cross-prediction scores [left: RBP1’s predictions on RBP2’s contexts; right: RBP2’s predictions on RBP1’s contexts] for RBP-RBP interactions discovered by this work (green) and interactions discovered by using peak overlaps alone (orange). Statistical testing performed using a Wilcoxon test. (B) Histogram of eCLIP peak lengths for peaks from all RBPs in HepG2 that were used in this study. (C) Distribution of eCLIP peak lengths for peaks from each RBP in HepG2 that were used in this study. Statistical differences (assessed by the Wilcoxon test) are present between distributions (not shown).

**Supplemental Figure S18: eCLIP read enrichments supporting the proposed interaction between AKAP1 and UPF1.** (A) Nested violin-boxplots of the distribution of (A) log2(AKAP1 eCLIP read enrichment over size-matched input [SMInput]) and (B) log2(UPF1 eCLIP read enrichment over SMInput) for (i) AKAP1 contexts that do not overlap with UPF1 contexts (orange), (ii) AKAP1 contexts that overlap with UPF1 contexts (green, left), (iii) UPF1 contexts that overlap with AKAP1 contexts (green, right), and (iv) UPF1 contexts that do not overlap with AKAP1 contexts. In panel A, Wilcoxon rank-sum test *p*-values test whether the distribution on the right-hand side is significantly greater than the distribution on the right-hand side. In panel B, Wilcoxon rank-sum test *p*-values test whether the distribution on the right-hand side is significantly greater than the distribution on the left-hand side.

## SUPPLEMENTAL TABLES

Supplemental_Table_S1.xlsx: ENCODE data experiment and file accessions

Supplemental_Table_S2.xlsx: MIL performance and statistics

Supplemental_Table_S3.xlsx: Reduction in witness rate

Supplemental_Table_S4.xlsx: RBFOX2 motif instance itemization

Supplemental_Table_S5.xlsx: Motifs, contexts, and nucleotide preferences for all RBPs in HepG2

Supplemental_Table_S6.xlsx: Motifs, contexts, and nucleotide preferences for all RBPs in K562

Supplemental_Table_S7.xlsx: eCLIP enrichment statistical testing for motifs and contexts relative to controls

Supplemental_Table_S8.xlsx: Motif discovery ground-truth comparison

Supplemental_Table_S9.xlsx: *k*-mer enrichments in sequences and contexts for select RBPs

Supplemental_Table_S10.xlsx: Context discovery comparison and characterization

Supplemental_Table_S11.xlsx: Motif characterization

Supplemental_Table_S12.xlsx: Cross-prediction performance

Supplemental_Table_S13.xlsx: Features supporting known interactions

Supplemental_Table_S14.xlsx: Interaction discovery for all RBP-RBP pairs

Supplemental_Table_S15.xlsx: Features of candidate interacting rG4-binding proteins

Supplemental_Table_S16.xlsx: Features of spliceosomal proteins

