## Supplemental Notes for "A novel NLP-based method and algorithm to discover RNA-binding protein (RBP) motifs, contexts, binding preferences, and interactions"

### SUPPLEMENTAL NOTE S1

#### A novel linguistics-inspired sequence decomposition method and multiple instance learning (MIL) formulation for RBP binding prediction

##### Sequence definition

We used the ENCODE REST API to download merged replicate enhanced cross-linking and immunoprecipitation (eCLIP) peaks from the ENCODE portal (Luo et al. 2020; Van Nostrand et al. 2020a) (**Supplemental Table S1A**, **S1B**). For each RBP, we assigned each of its peaks to a genomic region based on the classification scheme and priority order from (Van Nostrand et al. 2020a). We anchored each peak at its center and extended in both directions by 50 bp to achieve a uniform sequence length of 101 bp (**Supplemental Figure S1A**). We also filtered out any sequences that remained unassigned by genomic region, contained non-canonical nucleotides, or originated from chromosomes other than chr1-22 and chrX. We defined the resulting sequences as our positive observations. To derive an analogous set of negative observations, we randomly sampled 101 bp sequences from matched genomic regions (to maintain representative nucleotide content) and matched chromosomes (to ensure balanced cross-validation groups), resampling as needed to ensure a disjoint set with respect to the positive observations (**Supplemental Figure S1A**). We divided our entire dataset into 12 cross-validation groups, assigning each sequence to a group based on its chromosome of origin, as in the chromCV method (Moore et al. 2020). For each RBP, we designated the largest of the 12 groups as the test dataset, with the remaining 11 groups serving as the training dataset.

##### Sequence decomposition

We drew on theoretical concepts from natural language processing (NLP) to design an appropriate method of sequence representation for RBP binding sites and their flanking regions. We operated under the assumptions that (1) each peak sequence represents a putative RBP binding site, (2) each binding site has a significantly greater abundance of enriched *k*-mers with respect to corresponding control sequences, and (3) these enriched *k*-mers contribute to the RBP-RNA interaction either as the motif instance being recognized by the RBP or as a contextual element within the flanking RNA sequence which promotes the binding event. To gain insight into RBP binding patterns, we sought to identify which of these enriched *k*-mers constitute the RBP binding site, and which comprise the binding site's flanking regions.

With this paradigm, we designed a strategy to deconstruct our sequences into a series of regions, which we term "contexts," each consisting of three elements: a central *k*-mer called the "target," representing a candidate RBP binding site, and two sets of multiple *k*-mers each representing the target's flanks, called the "left context" and "right context," respectively. We employ a sliding window-based decomposition strategy consisting of two stages. In the first stage, a sliding window of size $k_{target}$ and step $p_{target}$ was used to deconstruct a given sequence into a series of consecutive *k*-mers, each of which represents a potential target (**Supplemental Figure S1B**); these series of *k*-mers, each representing a full sequence, were used to train the baseline model (see below). In the second stage, a second sliding window of size $k_{contexts}$ and step $p_{contexts}$ was used to isolate a specified number of *k*-mers $n_{contexts}$ in the left and right flanking regions of a given potential target to generate the left and right context, respectively, for that target (**Supplemental Figure S1C**). The derived structure, comprising the left context, target, and right context, is collectively referred to as a "context"; these contexts were used to train the context model (see below).

Our approach introduces a new representation structure that clearly demarcates the significant entities constituting an RBP binding event (i.e., target and context), and it does so in an unbiased manner by treating every *k*-mer as a candidate target for RBP binding. It is also highly flexible in implementation, with customizable parameters that enable exploration of targets and contexts independently having different *k*-mer lengths, window strides, region sizes, and gap presence. We decided to conduct our study using *5*-mers for both the target and context elements due to its demonstration as a robust and representative RBP motif length (Dominguez et al. 2018), a window stride of 1 to consider every target as a potential binding site, and 5 *k*-mers each in the left and right context elements to allow inspection of a respectably-sized region for contextual effects.

##### Formulation of the RBP binding prediction task as a Multiple Instance Learning (MIL) problem

Our main objective is to increase the resolution of RBP binding prediction by identifying the precise context(s) per positive sequence that are necessary for an RBP binding event to occur. Multiple Instance Learning (MIL) – a branch of weakly supervised machine learning characterized by its arrangement of observations as "bags" and subsets of those observations as the bags' "instances" (Dietterich et al. 1997; Herrera et al. 2016; Carbonneau et al. 2018) – is a particularly apt framework due to its natural ability to model the context data structure. In our formulation, we denote a sequence as a bag and the sequence's constituent contexts as the bag's instances (**Supplemental Figure S1A**). We adhere to the collective MIL assumption that a bag's instances constitute a random representative sample from the parent bag's inherent probability distribution, such that each instance has an equal contribution to the label of the bag (Foulds and Frank 2010), but the instance labels themselves are unknown (Herrera et al. 2016).

Within this MIL framework, we sought to determine the identity of each instance label in order to discern which region(s) within a sequence significantly contribute to the associated RBP binding event. To this end, we trained neural networks to predict RBP binding using standard encoding, embedding, and hyperparameter techniques from NLP and deep learning (**Supplemental Figure S1D**; see below). To assign each instance an initial label for training, we implemented a label inheritance strategy (Carbonneau et al. 2018) that assigns the sequence label to all contexts derived from that sequence; (i.e., all contexts derived from positive sequences were labeled positive and all contexts derived from negative sequences were labeled negative). We then performed binary classification at the instance-level to determine the identity of positive context(s) per sequence – which signify a contribution to RBP binding – and at the bag-level using an averaging aggregation technique, which allows assessment of model performance with respect to the ground-truth sequence labels (Herrera et al. 2016; Foulds and Frank 2010).

### Model pre-processing, construction and training pipeline (Supplemental Figure S1D)

#### *Data encoding and embedding*

We trained a keras (Chollet and Others 2015) Tokenizer on the training dataset to compile a dictionary of *k*-mers present in our corpus. We then used the trained Tokenizer to encode the data as either a Bag-of-Words $m$ x $n$ matrix (such that $m$ = number of observations and $n$ = vocabulary size) or with an index-based encoding using the dictionary's word index (such that each constituent *k*-mer in the context would be represented by its index in the learned dictionary); in the latter case, the encoded form is fed as input to the Embedding layer of the neural networks. We indicate below, as needed, the encoding method used in each step of model training. We used the trained Tokenizer to apply the same operations to the test dataset.

#### *Constructing the model architecture*

We tested multiple different neural network architectures: namely, a shallow neural network containing a sole hidden layer with either a Bag-of-Words encoding (for the baseline model; see below) or an embedding layer (for the context model; see below); a convolutional neural network (CNN) with max pooling; and three forms of a recurrent neural network (RNN), each consisting of a convolutional layer, max pooling layer, and a different RNN architecture (long short-term memory [LSTM], bidirectional LSTM, and gated recurrent unit [GRU]). In preliminary testing stages of the project, we found that the more complex model architectures did not confer a significant advantage in overall performance compared to a simple neural network model. In line with our goals to devise an architecture-independent method for context discovery and analysis, we thus settled on using a simple neural network architecture for this study. Our neural networks contain an embedding layer (in the case of our context model, whose input is our encoded context data), a hidden layer with a ReLu activation function (whose size, or number of neurons, is determined by values from hyperparameter tuning [see below]), a dropout layer with a rate of 0.5 (to counter overfitting) and an output layer with a sigmoid activation function (to enable binary classification). We used the Adam algorithm for stochastic gradient descent and the binary cross-entropy loss function, which is suitable for binary classification tasks (Chollet and Others 2015).

#### *Model training and hyperparameter tuning*

For RBP binding prediction, we first trained a baseline model on the *k*-mer based sequence representation from the first stage of sequence decomposition (encoded as a Bag-of-Words matrix) to obtain a reference model. Next, we trained an initial contexts model on the contexts from the second stage of sequence decomposition (encoded as a series of dictionary indices). We performed hyperparameter tuning to attain baseline and context models for each RBP with the strongest performance. To minimize runtime, we opted to use the RandomizedSearchCV approach from scikit-learn (Pedregosa et al. 2011) as opposed to the GridSearchCV approach, which exhaustively tests each combination of hyperparameter values. We considered 4 hyperparameters influential to the neural network's performance, and we specified the following ranges for each hyperparameter based on preliminary exploratory analyses (not shown):

- Embedding dimension: [8, 16, 32, 64, 128]
- Number of neurons: [16, 32, 64, 128, 256, 512, 1024, 2048]
- Batch size: [32, 64, 128, 256, 512, 1024, 2048]
- Number of epochs: [2, 3, 4, 5, 6, 7, 8, 9, 10]

We performed 25 iterations of random hyperparameter value sampling, training, cross-validation, and testing. To select the optimal set of hyperparameters from the tuning results, we defined criteria that balanced a model's generalization gap (measured by the difference in training and test performance) with its overall performance (measured by the test performance). We set a small threshold (starting at 0.005) and identified the tuning iterations whose difference in training and test performance was less than this threshold; if no tuning iterations met this requirement, we progressively increased the threshold at small increments until at least one tuning iteration demonstrated a generalization gap below the threshold. Next, for the subset of iterations that met the generalization gap requirement, we selected the iteration that had the maximum test performance for the baseline and context models, respectively. Using the optimized hyperparameter values from the selected iterations, we trained the final baseline and contexts models on the full training dataset.

### Iterative relabeling

In our goal to identify the precise regions within a positive sequence that contribute to its binding event, we sought to filter out noise that is intrinsic to label inheritance; in other words, we sought to remove weak contexts within positive sequences that do not possess salient features identified by the predictive model. To this end, we implemented a strategy that we term iterative labeling, in which each training iteration's instance-level predictions are used to reassign context labels prior to the subsequent iteration of training (**Supplemental Figure S1D**).

First, using the baseline model as our gold standard, we ran the set of contexts (encoded as a Bag-of-Words matrix) through the trained baseline model to get the baseline's predictions on the contexts, then used the resultant predictions to assign new labels to the contexts. Contexts originating from negative sequences were not relabeled, as they are not expected to contain any salient features and are not associated with a binding event. Contexts originating from positive sequences (i.e., having an initial label of 1) were relabeled based on whether their respective predicted probabilities were greater than 0.5 (relabel as 1) or below 0.5 (relabel as 0), a process which we refer to as relabeling.

We then performed, in an iterative manner, additional rounds of both retraining the contexts model and relabeling the contexts using the resultant predictions; we refer to this process as iterative relabeling. Note that iterative relabeling should be run until a given termination condition is met; for example, until convergence, until a pre-defined number of iterations is completed, or until the first occurrence of overfitting is detected. To minimize runtime, we ran iterative relabeling for a total of 3 iterations. To select the best-performing iteration of relabeling, we implemented a procedure akin to that for the selection of the best-performing iteration of hyperparameter tuning (see above), incrementally applying a generalization gap threshold to identify iterations with the smallest difference in training and test performance, then selecting from these the iteration with the highest test performance.

### Evaluation of model performance

#### *Area under the precision-recall (AUPR) curve*

Because the data was imbalanced after relabeling (with respect to the relative number of positive and negative observations), we used the area under the precision-recall (AUPR) curve to evaluate the performance of our context predictions (Grau et al. 2015). We computed AUPRs at the instance- and bag-levels before and after relabeling. Our models generally performed well, with a final median bag AUPR of 0.865 in HepG2 (**Figure 1A**; **Supplemental Table S2A**, **S2B**) and 0.861 in K562 (**Figure 1B**; **Supplemental Table S2C**, **S2D**). Despite an apparent dispersion in the distribution of post-relabeling context performance, we attribute this phenomenon to the natural discrepancy between instance labels of two successive relabeling iterations.

***Computation of MIL witness rates***

We computed the witness rates (i.e., number of positive contexts per bag) (Carbonneau et al. 2018) per sequence before and after relabeling. To survey the global witness rate across all contexts belonging to a given RBP, we computed the proportion of all contexts that were assigned a positive label by our predictive model for that RBP. We then computed summary statistics, such as the median and average, to report global witness rates across all RBPs (**Supplemental Table S3**). Because our initial dataset was balanced (i.e., equal number of positive and negative observations), by definition the starting global witness rate for every RBP is 0.5. To inspect the witness rates at the local level, we computed the proportion of positively-predicted contexts per true positive sequence, performed an average across all sequences for a given RBP, and again calculated summary statistics. Because each positive sequence is initially composed of solely positive contexts due to inherited labeling, by definition the starting local witness rate for each such sequence is 1.0. We computed and compared the global and local witness rates at successive stages of training to examine the effects of iterative relabeling.

Notably, after relabeling, the bag performance was largely preserved even as the witness rate substantially decreased – by about 1.4-fold at the global level and 1.8-fold at the local level, in both cell lines (**Supplemental Table S3**) – indicating that our trained models could successfully achieve the removal of labeling noise from positive sequences while simultaneously maintaining bag-level prediction accuracy.

### SUPPLEMENTAL NOTE S2

#### A novel algorithm for the accurate discovery of RBP binding motifs and contexts

The motif discovery algorithm consists of 6 phases, described below.

**Phase I: Identification of candidate consensuses**
In the first phase, in accordance with our initial assumptions that RBP binding sites consist of overrepresented patterns, we used *k*-mer enrichments to construct a set of strong candidate consensuses, such that the set consisted of all *k*-mers that were enriched as targets of positively-predicted contexts with respect to negatively-predicted contexts (**Supplemental Figure S2A**). By only considering enriched *k*-mers as candidate consensuses, we simultaneously reduce the motif search space.

To compute *k*-mer enrichments, we designated a context as positive if its prediction was greater than 0.5, and negative otherwise. For each unique target *k*-mer, we computed its enrichment based on its frequency in the positively-predicted contexts relative to its frequency in the negatively-predicted contexts. We also used these quantities to perform chi-square testing and obtain a *p*-value for each *k*-mer indicating the significance of its enrichment. We only considered all enriched *k*-mers (i.e., enrichment > 1) as candidate consensuses, disregarding all other non-enriched *k*-mers in the motif discovery process to reduce the motif search space.

**Phase II: Identification of position-dependent similar k-mers**
In the second phase, to model the relative sequence similarity observed between a motif's consensus and its constituent instances (D'haeseleer 2006b, 2006a), and to account for the degenerate nature of RBP motifs (Müller-McNicoll and Neugebauer 2013), for each candidate consensus we assembled a preliminary list termed a "partition" containing all enriched *k*-mers with restricted sequence similarity to that consensus.

First, to create a preliminary partition for each candidate consensus, we generated one list for each nucleotide position of the consensus, where each list contained all other enriched *k*-mers that had the same nucleotide as the consensus at that position; we refer to this as position-based similarity (**Supplemental Figure S2B**). We then merged these lists by taking the intersection of the inner and outer lists, respectively, then taking the union. This operation yielded a preliminary partition consisting of *k*-mers with at most 3 mismatches (i.e., a Hamming distance of 3) relative to the corresponding candidate consensus, thus modeling the degeneracy of RBP motifs. Note that a preliminary partition is constructed on the basis of similarity to the corresponding consensus. While we use the Hamming distance as a measure of similarity between partition *k*-mers and their consensus, we do not use it to construct the partition itself.

**Phase III: Identification of motif instances with k-mer co-occurrence**
The *k*-mer enrichment and similarity requirements proved insufficient for accurate motif discovery. We thus required an additional constraint for fine-tuning each initial partition to contain only the most highly-probable instances for a given motif. We had previously observed in our data an intriguing relationship between a motif consensus and any putative motif instance, whereby the two *k*-mers occurred within the same sequence at some frequency; we termed this feature the *k*-mer co-occurrence. We also noticed that by enforcing a *k*-mer co-occurrence threshold, we could filter out *k*-mers that were less likely to be part of the true motif. In the third phase, to arrive at a final, filtered partition for each RBP, we developed a programmatic, convergence-based minimization approach to tune the *k*-mer co-occurrence threshold, such that only those *k*-mers belonging to the initial partition that met this threshold would be retained in the final partition (**Supplemental Figure S2C**).

To remove *k*-mers from the preliminary partition that are not likely to be true motif instances, we relied on each *k*-mer's co-occurrence with the corresponding candidate consensus. We defined the *k*-mer co-occurrence frequency as the frequency at which a particular partition *k*-mer occurred in the same sequence as its corresponding candidate consensus, then used a co-occurrence threshold to filter out any *k*-mers whose co-occurrence frequency was above this threshold. To determine the value of the co-occurrence threshold, we developed a minimization-based tuning procedure. We initialized the tuning procedure at a co-occurrence threshold of 0.5 and progressively increasing the threshold by small increments until a ceiling of 1; at each iteration, we computed the co-occurrence frequency of each *k*-mer in the partition, filtered out any *k*-mers whose frequency did not satisfy the threshold, constructed the resultant motif (see below) based on the remaining *k*-mers in the partition, and computed the Kullback-Leibler Divergence (KLD) between the motif PFMs from the successive iterations. We continued incrementing the threshold as long as the successive KLDs continued to decrease, and stopped tuning if the KLD began to increase (i.e., we minimized the difference between probability distributions of motifs at successive co-occurrence thresholds). We also performed tuning in the opposite direction by decreasing the threshold from 0.5 in small increments (until a floor of 0) and repeating the minimization procedure. To arrive at the final co-occurrence threshold, we selected the motif corresponding to the smallest difference in KLD between the two tuning directions.

Taken together, *k*-mer enrichment, *k*-mer similarity, and *k*-mer co-occurrence are the three necessary and sufficient conditions for our algorithm's successful motif discovery.

**Phase IV: Motif construction**

In the fourth phase, we constructed the final motifs (**Supplemental Figure S2D**). For all *k*-mers in a candidate consensus's final partition, we collated all of the *k*-mers' instances where they occurred as targets of positively-predicted contexts. For cases in which more than one motif instance was present within the same sequence, we retained only one *k*-mer by prioritization using the following conditions: (1) if the identities of the instances were different, we retained that with the higher enrichment; (2) if the identities of the instances were the same, we retained that with the higher prediction score. We then aligned the compiled instances to generate the corresponding motif for that consensus. We visualized the motif logos using WebLogo (Crooks et al. 2004) and Logomaker (Tareen and Kinney 2020).

**Phase V: Motif scoring and primary motif selection**

Our algorithm discovers all possible motifs for a given RBP dataset. In the fifth phase, we select a primary motif by using a multi-metric, iterative sorting strategy based on the enrichment and statistical significance of the motif's consensus as well as the weighted relative entropy (Tompa 1999) of the final motif.

For each RBP, the algorithm discovers all possible motifs within the dataset that satisfy the algorithm's conditions; the number of discovered motifs is different for each RBP. In the fifth phase, to select a primary motif for each RBP, we developed an iterative sorting strategy relying on the enrichment and statistical significance of the motif's consensus, as well as the weighted relative entropy (Tompa 1999) of the motif. Briefly, we first sorted motifs by the enrichment of the consensus *k*-mer, selecting the top 10 for further consideration. We then sorted these 10 motifs by the significance of their consensus's enrichment, selecting the top 5. We next sorted these 5 motifs by their weighted relative entropy (Tompa 1999), selecting the top 2. From these 2, we designated the motif whose candidate consensus had the highest enrichment as the primary motif for the RBP, which we used to perform all subsequent analyses.

**Phase VI: Context discovery**

In the sixth and final phase, we derived the motif's context by extracting the flanking regions surrounding each motif instance using symmetrical extension. We aligned the resulting 55-nt long regions and visualized the logos for the contexts using WebLogo (Crooks et al. 2004) and Logomaker (Tareen and Kinney 2020) (**Supplemental Figure S2E**).

### SUPPLEMENTAL NOTE S3

#### Comments on the quantitative validation of the motif discovery algorithm

We evaluated the accuracy of our motif discovery algorithm by cross-referencing with a ground-truth validation set containing RBPs with well-characterized motifs that have been concordantly reported across multiple publications. Based on the ground-truth sets in both HepG2 and K562, our algorithm successfully discovered motifs for 13 of the 14 RBPs, yielding a consistent accuracy of ~92.9% across both cell lines. In this Supplemental Note, we comment on the RBPs whose discovered motifs did not correspond with known motifs from the literature.

**PABPN1 (HepG2)**

In HepG2, the discovered motif for PABPN1 differed significantly from the motif reported in the literature. PABPN1 is a nuclear poly(A) binding protein with established roles in the regulation of the poly(A) tail and alternative polyadenylation (Huang et al. 2023). We discovered a motif with consensus GCUGG (**Supplemental Figure S3A**). Upon closer inspection, we found that the poly(A) *5*-mer was neither enriched in our baseline predictions (0.271-fold enriched; *p*-value = 4.83e-6; rank = 880; **Supplemental Table S9A**), nor was it enriched in our contexts predictions (0.500-fold enriched; *p*-value = 0.0111; rank = 637; **Supplemental Table S9B**), suggesting that the absence of the poly(A) *5*-mer is inherent to the dataset; other A-rich *5*-mers were similarly depleted. We also submitted our PABPN1 context predictions to STREME (Bailey 2021), the results of which also confirmed the lack of a poly(A) motif (**Supplemental Figure S3B**). Our primary motif GCUGG was ranked #7 by STREME, while our secondary motif GGGCC was ranked #5 by STREME. Finally, we confirmed that our PABPN1 motifs and contexts were significantly enriched in eCLIP reads with respect to a set of random controls from matched genomic regions (**Supplemental Figure S3C**; **Materials and Methods**).

Based on these results, we hypothesize that our discovered motif represents either (1) a novel motif for PABPN1 that reflects a non-canonical function other than the regulation of poly(A) tail formation in the nucleus (Huang et al. 2023) or (2) the motif for an as-of-yet unknown, RNA-bound RBP with which PABPN1 physically interacts, in which case the PABPN1-specific antibody would immunoprecipitate mRNAs crosslinked with the other RBP. These hypotheses may be supported by the fact that over 60% of PABPN1's discovered motif instances are located in coding sequences (not shown) and by previous reports of PABPN1 interacting with numerous spliceosomal proteins (Huang et al. 2023), which together could point to a motif with PABPN1-mediated roles in mRNA splicing.

**RBFOX2 (K562)**

The algorithm discovered the canonical GCAUG motif for RBFOX2 in both cell lines. However, while the algorithm selected GCAUG as the primary motif in HepG2, it did not do so in K562, instead selecting GGGGC as the primary motif (**Supplemental Figure S4A**). An inspection of the K562 dataset revealed explanations for this phenomenon. Firstly, we compared the enrichment and significance of both GCAUG and GGGGC in the two cell lines (**Materials and Methods**). GCAUG was significantly and highly enriched in HepG2 context predictions (12.4-fold enriched; $p$ = 2.51e-128; rank = 1) while GGGGC was much more lowly enriched (5.91-fold enriched; $p$ = 7.74e-86; rank = 13) (**Supplemental Table S9C**). Conversely, in K562, GGGGC was significantly and highly enriched in the context predictions (8.20-fold enriched; *p*-value = 4.05e-77; rank = 2) while GCAUG was very lowly enriched by comparison (3.95-fold enriched; $p$ = 3.01e-15; rank = 28) (**Supplemental Table S9D**). Because the algorithm's iterative sorting strategy for primary motif selection relies on both enrichment and significance of the consensus (**Supplemental Note S2**), the low enrichment of GCAUG in K562 may help to explain why its corresponding motif was not ranked as the primary motif for RBFOX2 in that cell line.

These findings suggest that GCAUG is not strongly overrepresented in RBFOX2 sequences originating from the K562 dataset. Indeed, whereas GGGGC was 11.3-fold enriched ($p$ = 5.79e-42) in positive sequences and ranked #3 by enrichment, GCAUG was only 1.44-fold enriched ($p$ = 0.036) in positive sequences and ranked #224 by enrichment (**Supplemental Table S9E**). By contrast, GCAUG was more highly and significantly enriched in HepG2 positive sequences (4.42-fold enriched; $p$ = 1.55e-54; rank = 13) than GGGGC (3.97-fold enriched; *p*-value = 6.90e-40; rank = 19) (**Supplemental Table S9F**). Furthermore, while the percentage of all sequences containing at least one occurrence of GGGGC was relatively high in both cell lines (HepG2: 39.3%; K562: 51.9%), there was a large discrepancy between the two cell lines for GCAUG (HepG2: 54.3%; K562: 29.1%), with the percentage of sequences containing at least one occurrence of GCAUG drastically lower in K562 compared to HepG2. Taken together, these observations help to explain why the algorithm did not select GCAUG as the primary RBFOX2 motif in K562.

The top-ranking motifs by STREME for RBFOX2 in K562 are also predominantly G-rich (**Supplemental Figure S4B**). We further demonstrate that our discovered motif and context in K562 are significantly enriched in eCLIP read coverage compared to matched random control regions (**Supplemental Figure S4C**).

### SUPPLEMENTAL NOTE S4

#### Random motif discovery as a negative control

To demonstrate that our discovered motifs and contexts represent true positive hits, we also validated our algorithm using random "peaks" as negative controls. To this end, we generated two sets of random sequences – the first derived from the human genome ("random genomic") and the second generated from a uniform distribution of the 4 canonical nucleotides ("random synthetic") (**Materials and Methods**) – then applied our motif discovery algorithm on these sets, testing sample sizes of 2000, 10000, and 20000 sequences. Our motif discovery algorithm returned hits for both types of random sequences, as did STREME (**Supplemental Figure S6**, **S7**). We observed a marked difference in the context nucleotide preferences between random genomic and random synthetic sequences, with the preferences for the former reflecting the underlying nucleotide distributions of the genomic regions of origin (**Supplemental Figure S6C**) and the preferences for the latter reflecting the uniform distribution of the canonical nucleotides (**Supplemental Figure S7A**, bottom). On the other hand, the contexts for RBPs annotated as non-sequence-specific demonstrate preferences for particular nucleotides (**Supplemental Table S5**), which is distinct from the patterns observed in these random negative controls.

We first remark that even a purely random sample of sequences is expected to contain some significantly enriched *k*-mers, although considerably fewer in number and lower in enrichment than would be observed in a true sample. For this reason, any motif discovery algorithm will discover and construct a motif, whether it is truly meaningful or not. Although any *k*-mer with an enrichment value greater than 1 (and a *p*-value less than 0.05) is considered significantly enriched, the significantly enriched *k*-mers in a purely random sample have enrichment values between 1 and 2, indicating that they are narrowly considered overrepresented; this is in stark contrast to true samples, which instead have *k*-mer enrichment values substantially greater than 1, and even surpassing 50 in some cases.

Despite the fact that both our algorithm and STREME returned motif hits for this set of negative controls, we noted several distinctive differences in sequence composition, feature constitution, and structural properties that clearly distinguish the motifs and contexts discovered from RBP datasets from those discovered from random sequences. First, when we applied our MIL prediction method on the random genomic and random synthetic sequences, the trained models performed only as well as random classifiers when 50% of observations in the dataset are positive, yielding AUPRs of around 0.5 (minimum AUPR = 0.482 for random genomic 2K; maximum AUPR = 0.515 for random synthetic 2K) (**Supplemental Figure S8A**, **S8B**). The inability of these models to differentiate between positive and negative sequences signifies that the foreground and background distributions, respectively, of nucleotides and *k*-mers do not present any significantly discriminating features for the model to learn in order to distinguish between positive and negative sequences. Furthermore, although a small proportion of significantly enriched *k*-mers are present, they are not overrepresented enough to constitute salient features from which the model can learn to differentiate between positive and negative sequences. In stark contrast, the random performance was significantly lower than the performance observed for a subset of sequence-specific RBPs, whose AUPRs ranged from 0.892 for SRSF1 to 0.981 for HNRNPK (**Supplemental Figure S8A**), and for a set of non-sequence-specific RBPs, whose AUPRs ranged from 0.79 for DDX55 to 0.924 for DDX59 (**Supplemental Figure S8B**). In the latter case, the notably higher performance compared to random signifies that even the binding sites of non-sequence-specific RBPs contain true and robust latent features that are strongly predictive of binding. The performance observed for sequence- and non-sequence-specific RBPs were largely comparable.

Second, when we inspected the motif consensuses for RBPs and random sets, we observed marked differences between the two. When considering all 71 RBPs in HepG2, the enrichment for a motif consensus (CDC40) reached as high as 86.914 (**Supplemental Figure S8C**, first violin-boxplot). The consensus enrichments of motifs discovered for a subset of 7 sequence-specific RBPs were also very large, ranging from 4.337 to as high as 50.722 (**Supplemental Figure S8C**, second violin-boxplot; median = 15.898). While the range of motif consensus enrichments for non-sequence-specific RBPs did not demonstrate the same spread (minimum = 6.232, maximum = 16.734), the median motif consensus enrichment (10.792) for non-sequence-specific RBPs was comparable to that for sequence-specific RBPs (**Supplemental Figure S8C**, third violin-boxplot). Conversely, the motif consensus enrichments for all random datasets, which were greater than 1 but less than 2, were significantly lower than those of the RBP datasets; the median enrichment for random genomic motif consensuses was 1.285 (minimum = 1.161, maximum = 1.592; **Supplemental Figure S8C**, fourth violin-boxplot), while the median enrichment for random synthetic motif consensuses was 1.187 (minimum = 1.163, maximum = 1.438; **Supplemental Figure S8C**, fifth violin-boxplot).

Similarly, the consensus *p*-values for RBP motifs were considerably smaller than those for random motifs, with the median *p*-values for the entire set of RBPs, the subset of 7 sequence-specific RBPs, and the set of non-sequence-specific RBPs being 2.89e-68, 6.45e-199, and 1.81e-57, respectively (**Supplemental Figure S8D**, first three violin-boxplots); many motif consensuses even had *p*-values approaching 0. Conversely, the consensus *p*-values for all random motifs border 0.05, with the medians for random genomic and random synthetic motifs being 0.00249 and 0.0103, respectively (**Supplemental Figure S8D**, fourth and fifth violin-boxplots).

We observed similar patterns when comparing the enrichments and *p*-values of all significantly enriched *k*-mers from random datasets with those from sequence-specific RBPs (**Supplemental Figure S9A**, **S9B**) and non-sequence-specific RBPs (**Supplemental Figure S9C**, **S9D**). In summary, we found that (1) the enrichments for significantly enriched *k*-mers from RBP datasets were much greater (i.e., exceeding far beyond 2) than those from random datasets, which lie very close to 1 but do not exceed 2; (2) the *p*-values for significantly enriched *k*-mers from RBP datasets were much smaller (i.e., approaching 0) than those from random datasets, which lie close to 0.05; and (3) the results from sequence- and non-sequence-specific RBPs both lack the randomness of features that are present in the random datasets, and (4) the results from sequence- and non-sequence-specific RBPs are comparable to one another. Importantly, we also noted that there were much fewer significantly enriched *k*-mers for the random datasets (i.e., between 15 - 40) compared to the RBP datasets (i.e., on the order of magnitude of $10^2$) (**Supplemental Figure S9**).

Taken together, these analyses demonstrate that it is the properties of the discovered motifs and contexts, rather than the presence or absence of a hit by a motif discovery algorithm, that are informative of a good negative control. In addition, both negative control sets utilized here demonstrate that our algorithm discovers meaningful RBP motifs and contexts that are clearly distinct from those discovered from random genomic and random synthetic sequences. This is of particular importance for interpreting the results of non-sequence-specific RBPs, which are typically known to bind to the RNA phosphate backbone rather than to nucleotide bases. We hypothesize that the discovered motifs and contexts for non-sequence-specific RBPs, which display preferences distinct from random negative controls, may be reflective of their frequent association with other sequence-specific RBPs, thus causing the CLIP crosslinking and pulldown to capture regions of RNA that demonstrate the sequence preferences of the interacting partner; we propose such a case later in this manuscript. Importantly, these analyses definitively demonstrate that non-sequence-specific RBPs do not constitute a valid negative control for motif discovery, as their motifs and contexts are significantly different from those discovered from random sequences; thus, although these RBPs are classified as non-sequence-specific, the sequences associated with their binding sites are non-random.

### SUPPLEMENTAL NOTE S5

#### Preliminary investigation into motif clusters

To conduct a preliminary investigation into motif clusters, we would first like to recall some important assumptions underlying our framework (**Supplemental Note S1**, **S2**) that are relevant to the topic of motif clusters. We assume that RBP binding sites are composed of a set of *k*-mers that are enriched relative to unbound sequences, and that binding sites contain a greater amount of enriched *k*-mers than unbound sequences do. We also assume that these enriched *k*-mers are clustered together to constitute a region, and we designate these as significant regions that contribute to the associated RBP binding event. In our motif discovery algorithm, we define motif instances as *k*-mers that are not only highly enriched, but that additionally satisfy the conditions for *k*-mer similarity and *k*-mer co-occurrence with the consensus.

To investigate clusters of motif sites that occur together, we selected HNRNPK, a known strong poly(C)-binding protein for which we discovered C-rich motifs and C-rich contexts. For each of the 55 nt sequences comprising the HNRNPK context logo in HepG2 (**Figures 2B**, **3C**), we first deconstructed the sequence into a series of *5*-mers, resulting in 51 *5*-mers in total. We then assessed the number of discovered HNRNPK motif instances that were present in each sequence, finding that, on average, each sequence contains about 5 motif instances (**Supplemental Figure S10A**). More broadly, the majority of sequences contained between 3 and 9 motif instances, while other sequences displayed even larger numbers of motif instances and one sequence containing a maximum of 23 motif instances. We also note that positions within a sequence that do not contain a motif instance still contain enriched *k*-mers that constitute significant regions within the sequence. For the motif instances that occur in the cluster but that are not ultimately selected as a binding site, these motif instances serve additional roles: (1) as part of the context, and/or (2) as the binding site for other interacting RBPs.

Next, we visualized the location of these motif instances across the full length of the sequences to better understand their positional distribution (**Supplemental Figure S10B**). We observe a peak at the central position - which corresponds to the location of the actual motif - indicating that 100% of the sequences contain a motif instance at that position, as expected. The remaining data is distributed relatively uniformly around this central position, eventually leveling off on both sides of the motif. We observe that approximately 7-8% of sequences contain a motif instance at the flanking positions that are farther than 5 nt from the central position. To illustrate an example, we also visualize the positional distribution of motif instances for one sequence (**Supplemental Figure S10C**).

To compare HNRNPK - an RBP for which we hypothesize binds motif sites located in clusters - with an RBP for which this behavior is not expected, we performed the same analysis on RBFOX2. Indeed, we found that sequences from the RBFOX2 context logo (**Figure 1C**) contained much fewer motif instances than HNRNPK, at only about 2 motif instances per sequence on average, and the majority of sequences containing only between 1 and 3 motif instances (**Supplemental Figure S10D**). In the same vein, we found that approximately 2% of sequences contain a motif instance in the contexts of the central motif (**Supplemental Figure S10E**). Therefore, motif instances occur at much higher frequencies in HNRNPK contexts than in RBFOX2 contexts, which suggests that motif clusters may indeed be occurring near HNRNPK binding sites.

We note that these analyses are preliminary, and that the topic of motif clusters warrants extensive further investigation, which we foresee as important future work that is enabled by our framework.

### SUPPLEMENTAL NOTE S6

#### On the interpretation of cumulative Kullback-Leibler Divergence (KLD) values

We have previously described our rationale and approach for computing cumulative KLD values (**Materials and Methods**). While cumulative KLDs provide an aggregated measure of distribution similarity across the motifs and contexts, it must be noted that distribution similarity as measured by the cumulative KLD does not necessarily imply feature similarity. For example, if a few positions within the PPM are permuted, the resulting cumulative KLD may only differ slightly from the original (unpermuted) cumulative KLD, but the features of the PPM may differ greatly. Furthermore, because the cumulative KLD is a sum across many individual probability distributions, there are many different combinations of probability distributions that could produce that sum. Thus, we interpret the cumulative KLDs jointly with the cross-prediction scores, which *do* provide a measure of feature similarity: a high cross-prediction score indicates that the most highly significant salient features are shared between any two RBPs. Like the KLD, cross-prediction is a measure, not a metric; it too is asymmetric and does not satisfy the triangle inequality.

We demonstrate that a good cross-prediction score is indicative of low cumulative KLD. Due to the challenges in interpreting a cumulative KLD, as described previously, our objective here is to define a bidirectional implication between the two measures (cross-prediction and cumulative KLD), such that we can use feature similarity as an indication of distribution similarity, and vice versa. To this end, we begin by establishing a minimum acceptable cross-prediction score that serves as a threshold of good feature similarity. As described in **Supplemental Note S9**, after much experimentation and analyses, we decided on 0.65 as a minimum acceptable cross-prediction score that is indicative of acceptable salient feature similarity between any two RBPs. Accordingly, we consider only the RBP-RBP pairs that satisfy our cross-prediction criteria for RBP-RBP interaction discovery; namely, that the cross-prediction score must be at least 0.65 in one direction, and that there must be a certain degree of symmetry for the cross-prediction score in the opposite direction (**Supplemental Note S9**). In **Supplemental Figure S11A**, we observe a negative Pearson correlation of -0.27 between cross-prediction values and cumulative context KLD values. This correlation represents that an increase in cross-prediction score is correlated with a decrease in cumulative context KLD; that is, increased feature similarity correlates with increased distribution similarity.

To help provide context and meaning to the cumulative KLD values relative to the single KLD values, we adopted the following approach to transform and compare the two sets of values. For RBP-RBP pairs with good cross-prediction scores, we use the length-normalized maximum cumulative context KLD as an upper range for which to consider two probability distributions as similar to one another. In other words: the maximum cumulative context KLD that we observe from RBP-RBP pairs with sufficient cross-prediction is 58.836 (**Supplemental Figure S11A**). If we normalize this value by the full length of the contexts (i.e., 55 nt), then we arrive at an average of ~1.07, which we use as an upper bound of distribution similarity. We also partition the range [0, 1.07] to define the degree of similarity, as follows:

- [0, 0.25): high similarity
- [0.25, 0.5): moderate similarity
- [0.5, 0.75): modest similarity
- [0.75, 1.07]: low similarity
- (1.07, infinity): dissimilar

Then, for any given cumulative KLD value, we first divide it by the length over which it has been summed, then compare this average value to the threshold: if the average is greater than 1.07, we consider the distributions to be dissimilar;  if the average is less than 1.07, we consider the distributions to be similar, and we can further delineate the degree of similarity by using the defined partitions above. We demonstrate examples further below.

Although the correlation between cross-prediction and cumulative context KLD is low, it reflects our desired objective of a relationship between cross-prediction scores and KLD values. Since our model for interaction discovery operates on defining acceptable cross-prediction values in the range [0.65, 1.00] and acceptable (average) KLD values in the range [0, 1.07], we are assured that (1) along the neighborhood of the line of best fit, we have a strong correlation between cross-prediction and KLD, in that small changes in cross-prediction scores lead to small changes in KLD values and vice versa, and (2) by defining a range of acceptable values for both variables, we have a bidirectional implication for the relationship between cross-prediction and KLD, in that acceptable values of cross-prediction guarantee acceptable values of KLD and vice versa. Thus, we are assured this bidirectional relationship between acceptable cross-prediction and acceptable KLD values as long as they fall within the 2-dimensional region [0.65, 1.0] x [0, 1.07]. Further, for points within this pre-defined 2-dimensional region, good cross-prediction scores imply good KLD values and vice versa. Similarly, this leads to the conclusion that good feature similarity implies good distribution similarity and vice versa.

The negative correlation between cross-prediction and cumulative context KLDs is also demonstrated in **Supplemental S11B**, where we observe a general decrease in cumulative context KLD distributions when stratified by cross-prediction score intervals. Furthermore, when we look at the distribution of cumulative context KLD values for RBP-RBP pair with good cross-prediction scores, we see that the majority of RBP-RBP pairs (i.e., ~81%) have values between 5 and 30 (**Supplemental Figure S11C**).

To summarize, based on our findings described above, we state the following:

- Cross-prediction defines the degree of overall feature similarity between two RBPs
    - An increase in cross-prediction score implies an increase in feature similarity
- In general, KLD defines the degree of distribution similarity, as does the cumulative KLD, with some additional modifications, considerations, and implications, as noted above
    - A decrease in cumulative KLD implies an increase in distribution similarity
- Because there is a correlation between good cross-prediction scores and cumulative KLDs:
    - A good cross-prediction score, as defined within the range [0.65, 1.0], is associated with a good cumulative KLD value, as defined within the range [0, 1.07].
    - Partitions within the range of acceptable cumulative KLD values, as defined above, indicate the degree of distribution similarity.
    - An increase in cross-prediction score implies a decrease in cumulative KLD
    - Therefore, an increase in feature similarity implies an increase in distribution similarity.

### SUPPLEMENTAL NOTE S7

#### Comments on the proposed novel motif instances for RBFOX2

We leveraged our analyses from the RBP knockdown (KD) shRNA-seq ENCODE data to demonstrate the functionality of each instance in our discovered RBFOX2 motif, which comprises 10 unique instances (**Figure 1D**). Because RBFOX2 is primarily implicated in the regulation of splicing, we investigated the presence of each of RBFOX2's motif instances in differentially spliced (DS) skipped exon (SE) events (**Materials and Methods**). We have previously demonstrated that RBFOX2's motif consensus GCAUG is enriched in DS events (**Supplemental Figure S15**). When we investigated the other 9 instances in our motif, we found that RBFOX2's other primary instance, GCACG, was also located in DS events, as expected. Of the 4 secondary instances in our motif that were also identified in (Begg et al. 2020), we found DS events associated with GCUUG, GUGUG, and GUUUG, but not for GUAUG. Finally, of the remaining 4 instances in our RBFOX2 motif that we propose as novel, we found DS events associated with GACUG, GCGUG, and GGUAG, but not for GUUAG. This analysis reveals that (1) noise may be a possibility for the single novel proposed motif instance that was not validated through KD data, and (2) our proposed novel instances demonstrate the same discovery rate in KD data as the secondary instances identified in (Begg et al. 2020). Importantly, we note that our KD analyses in this work are focused on 4 specific regions associated with the SE event class; it is very possible that the motif instances that did not occur in DS events are either located in other regions or associated with different splicing event classes other than SE events.

### SUPPLEMENTAL NOTE S8

#### Additional examples of striking cross-prediction patterns

We describe additional examples of groups of RBPs that demonstrate striking cross-prediction patterns, hypothesizing that these patterns may reveal underlying biological relationships between the RBPs in question.

MATR3 and SUGP2 demonstrated an interesting asymmetry, in which MATR3 showed a much weaker performance when predicting on SUGP2's contexts (AUPR = 78.34%) compared to SUGP2's prediction on MATR3's contexts (AUPR = 91.58%) (**Figure 6B**, **Supplemental Table S12**). Interestingly, SUGP2 performed even better on MATR3's contexts than it did on its own cognate contexts (AUPR = 82.27%) (**Supplemental Table S12**). These RBPs, both with U-rich motifs and contexts, consistently clustered together in all motif and context similarity analyses performed (**Figures 3**, **4**), and they are furthermore the sole members of their own cross-prediction cluster, indicating a strong similarity in global binding profiles.

We observe that 5 RBPs predict very highly on TRA2A (**Figure 6B**), and they are also part of the same cross-prediction cluster (**Figure 6C**), indicating that the 6 RBPs together share global binding patterns (**Supplemental Table S12**). BCLAF1 is the most highly similar to TRA2A, as the 2 RBPs consistently cluster together by motif and context similarities, and often as the sole members of the cluster in question. FXR2, GRWD1, HLTF, and ZNF800 also have AG-rich motifs and contexts, which could help to explain their strong performance on TRA2A. However, this cross-prediction pattern is strongly unilateral by way of TRA2A, as the other pairs within this set exhibit considerably low cross-prediction performance; the underlying reason remains to be seen.

Lastly, FUBP3 and KHSRP, two members of the same protein family (Gau et al. 2011), share high cross-prediction with SUB1 and ZC3H11A, suggesting a potentially novel relationship with RBPs outside of their family.

### SUPPLEMENTAL NOTE S9

#### A novel method to discover proximal interactions between RBP-RBP pairs

**Definition of proximal interaction modes for RBP-RBP interaction discovery**

We define the following 4 proximal interaction modes – 3 cooperative and 1 competitive – for consideration by our method:

- Interaction Mode 1 – **Cooperative** (proximal, dual-binding, physical): both RBPs bind the same target mRNA at distinct, non-overlapping sites to promote a shared regulatory goal. The binding sites of both RBPs are located near enough to each other such that the RBPs physically interact (**Figure 7A**, first leaf node).
- Interaction Mode 2 – **Cooperative** (proximal, dual-binding, non-physical): both RBPs bind the same target mRNA at distinct, non-overlapping sites to promote a shared regulatory goal. The binding sites of both RBPs are located far enough apart from each other such that the RBPs do not physically interact (**Figure 7A**, second leaf node).
- Interaction Mode 3 – **Cooperative** (proximal, single-binding, physical): the two RBPs physically interact with one another to regulate their target mRNA synergistically, but only one partner binds the target mRNA (**Figure 7A**, third leaf node).
- Interaction Mode 4 – **Competitive** (proximal, single-binding, non-physical): the two RBPs vie for the same or overlapping binding sites on a common mRNA target to exert opposing regulatory effects (**Figure 7A**, fourth leaf node).

**Definition of decision tree paths for RBP-RBP interaction mode classification**

We have previously described our formulation of RBP-RBP interaction discovery and our use of a decision tree in order to classify RBP-RBP pairs into interaction categories (**Figure 7B**). Based on our decision tree, we delineate the following 4 paths:

- Path I: RBPs demonstrate similar motifs and contexts, and their motif co-occurrence exceeds their motif overlap, suggesting **cooperative** mechanisms corresponding to Interaction Modes 1 and 2; whether or not the interaction is physical depends on the distance between their respective binding sites.
- Path II: RBPs demonstrate similar motifs and contexts, and their motif overlap exceeds their motif co-occurrence, suggesting either **cooperation** corresponding to Interaction Mode 3, in which the RBPs interact physically but only 1 partner binds RNA, or **competition** corresponding to Interaction Mode 4, in which the RBPs contend for the same or overlapping binding sites.
- Path III: RBPs demonstrate dissimilar motifs but similar contexts, and their motif co-occurrence exceeds their motif overlap, **cooperative** mechanisms corresponding to Interaction Modes 1 and 2; whether or not the interaction is physical depends on the distance between their respective binding sites. Path III differs from Path I in its condition for motif dissimilarity, while Path I requires motif similarity.
- Path IV: RBPs demonstrate dissimilar motifs but similar contexts, and their motif overlap exceeds their motif co-occurrence, suggesting either **cooperation** corresponding to Interaction Mode 3 or **competition** corresponding to Interaction Mode 4. Path IV differs from Path II in its condition for motif dissimilarity, while Path II requires motif similarity. Because dissimilar motifs are not likely to overlap very often, if at all, this path is expected to occur at very low frequencies.

If a candidate RBP-RBP pair does not meet the set of conditions required for any of the 4 paths, then our method considers it a non-interacting pair. Note that each of these 4 paths classifies an RBP-RBP pair to the interaction mode(s) for which it has the highest propensity, but does not preclude the possibility that the pair may interact via other modes. For example, for an RBP-RBP pair classified as cooperative according to Path I, a higher degree of motif co-occurrence than motif overlap suggests cooperation via Interaction Modes 1 and/or 2, but a non-zero motif overlap may still suggest the occurrence of interactions via Modes 3 and/or 4, albeit at lower frequencies. As such, we hypothesize that it is possible for a candidate RBP-RBP pair to exhibit both cooperative and competitive modes of interaction, but it may have stronger inclinations towards one or the other.

**Data preparation for discovery of RBP-RBP interactions**

Because the largest chromCV group from each RBP was used as its test dataset, and because the distribution of binding sites across chromosomes was different for each RBP, the contexts in the test predictions belonged to different chromosomes, making it difficult to conduct comparisons across RBPs. This particularly posed a challenge when proceeding to the RBP-RBP interaction analyses, as two important features for discovery are the relative degrees of motif instance overlap and motif instance co-occurrence (see below) which, by definition, must be detected in the same loci. To resolve this challenge, for each RBP we fit 11 additional new and independent models, such that for each model, one of the other 11 chromCV groups served as the test dataset and the remaining data served as the training dataset. For each of these additional models, we used the same hyperparameters that were used to train the original model for the RBP. Upon completion, we pooled the predictions across all 12 total models to arrive at a set of RBP binding predictions with coverage across all chromosomes, which enabled our RBP-RBP interaction discovery.

**Calculation of motif instance overlap and motif instance co-occurrence**

We used Bedtools (Quinlan and Hall 2010) to perform a strand-specific intersection of motif instances by genomic coordinate. We considered an intersection between any two motif instances – each belonging to an RBP within a given pair – by at least one nucleotide as a motif instance overlap. We performed the same operation for the contexts. We note that the resulting set of overlapping contexts contains two distinct subsets: one subset in which the corresponding motif instances themselves also overlap and the other subset in which the corresponding motif instances do not overlap. To obtain the motif co-occurrences, we performed a set difference between the full set of overlapping contexts and the set of overlapping motif instances to arrive at the set of overlapping contexts whose constituent motif instances do not overlap but instead co-occur. We divided each quantity by the total number of motif instances to obtain the corresponding percentages. We applied this procedure in a pairwise fashion to account for all possible RBP-RBP pairs.

**Application of the RBP-RBP interaction discovery method**

We implemented the decision tree in **Figure 7B** programmatically. To achieve this, we defined measures for each of the 4 characteristics in the decision tree: motif similarity, context similarity, motif instance co-occurrence, and motif instance overlap. Note that the order of operations of the conditions displayed in the decision tree in **Figure 7B** is not fixed but simply represents one possible implementation.

As a measure for motif similarity, we considered any two RBPs to have similar motifs if they belong to the same motif KLD cluster (**Figure 4**).

To define a measure of context similarity, we first considered two RBPs to have similar contexts if at least one partner had a cross-prediction score greater than 0.8, as an AUPR of this magnitude would suggest that the first RBP recognizes the majority of salient features in the contexts of the second RBP. We discovered 507 interactions, which accounted for 20.40% of the total 2,485 RBP-RBP pairs. In addition, we cross-referenced our discovered interactions with those in the BioGrid database (Oughtred et al. 2021) by downloading the BIOGRID-ALL-4.4.240.tab3.zip release and filtering out (1) non-human interaction data and (2) any interaction data for RBP-RBP pairs not present in our collection. Of the 2,485 possible pairs in our collection, 436 were documented as interacting in the BioGrid database, and we discovered 91 (20.87%) of these documented pairs. We note that while this database serves as a useful reference, it is not a comprehensive compendium of RBP-RBP interactions for two reasons: first, upon our survey of the RBP-RBP interactions within the database, most of these interactions were identified by experimental assays that detect physical interactions, and second, upon our survey of the literature, we identified reports of many RBP-RBP interactions that are not contained in BioGrid. However, cross-referencing our results with BioGrid still provides a method to approximate true positive discovery rates.

Upon examining our results, however, we noticed that a number of known interactions, including AKAP1-LARP4 (Gabrovsek et al. 2020) and EFTUD2-PRPF8 (Malinová et al. 2017), were not detected, along with other RBP-RBP pairs that we believe demonstrated strong signs of interaction but whose cross-prediction performance did not meet the threshold requirements (e.g., LARP4-UPF1, AQR-BUD13). Upon closer inspection, we observed that many of these pairs demonstrated a notable asymmetry, in which the first partner predicted relatively strongly (but below 0.8) on the second partner, but the second partner had a considerably lower score on the first partner. Despite this asymmetry, the presence of a higher score in one direction still indicates the presence of salient features in the second partner's contexts that are significant and similar to the salient features learned by the model of the first partner, and this pair could still be considered a potential interaction. We also took into consideration the importance of dataset size – in some cases, a smaller dataset for an RBP can affect model training such that the most significant salient features are not captured or learned by the model at a high resolution, which can lead to a lower score not only on its own contexts, but also on the contexts of other RBPs. Based on these facts, we concluded that a cross-prediction threshold of 0.8 was too stringent for interaction discovery, and we thus decided to lower the threshold in order to capture such cases in which there is a disparity in cross-prediction values. To address this, we tested different thresholds, cross-referencing the detected interactions with BioGrid for each test.

After experimentation, we defined two RBPs to have context similarity if at least one partner in the pair has a cross-prediction score on the second partner's contexts that is greater than 0.65. To ensure that the second partner's cross-prediction score on the first partner's contexts is not extremely low, we include an additional condition that its cross-prediction score must not be less than 5% of the 0.65 threshold. This definition resulted in the discovery of 796 interactions (32.03% of the total 2,485 RBP-RBP pairs in this study), with 145 of these interactions (33.36% of the 436) corresponding with the physical interactions in the BioGrid database, an increase of 85 additional interactions compared to the previous threshold at 0.8. Since our method considers only proximal interactions, we can infer that the 145 true positive interactions from BioGrid that were detected are proximal interactions. At these conditions, we also detected 289 more interactions overall than at the previous threshold of 0.8; although the cross-prediction scores for these 289 are lower, they still meet the other interaction conditions, which supports their discovery. Overall, these findings serve as a good validation for our interaction discovery method. Additionally, the increase in validated interactions from 91 at a threshold of 0.8 to 145 at a threshold of 0.65 suggests that higher cross-prediction thresholds can be too stringent; while the use of the lower 0.65 threshold may seem too relaxed, the detection of more reported physical interactions supports its validity. Furthermore, these findings demonstrate that an RBP-RBP pair does not have to exhibit very strong context similarity (close to 0.8 or higher) in order for the RBPs to participate in interactions with one another, provided that the other conditions of our method are also satisfied.

To determine the mode of interaction for the 796 pairs, we used inequality relationships between motif instance co-occurrence and motif instance overlap to facilitate classification of pairs into different paths (**Figure 7B**). For RBP pairs with a similar motif and similar context, we assigned them to Path I (cooperative) if their motif instance co-occurrence was greater than their motif instance overlap, and to Path II (cooperative and/or competitive) if their motif instance overlap was greater than their motif instance co-occurrence. Similarly, for RBP pairs with a dissimilar motif but a similar context, we assigned them to Path III (cooperative) if their motif instance co-occurrence was greater than their motif instance overlap, and to Path IV (cooperative and/or competitive) if their motif instance overlap was greater than their motif instance co-occurrence. Note that these classifications indicate the path (and interaction binding mode) that an RBP-RBP is most likely to follow; in other words, the interaction mode(s) that the RBP-RBP pair demonstrates the highest propensity for; however, it does not preclude the possibility that the RBP-RBP pair may interact via another mode if both their motif instance overlap and co-occurrence are non-zero, regardless of which is greater. Classifications for all 2,485 RBP-RBP pairs are available in **Supplemental Table S14**.

### SUPPLEMENTAL NOTE S10

#### Additional validation examples for proximal RBP-RBP interaction discovery

We describe additional examples of RBP-RBP pairs that serve as validation for our method of proximal interaction discovery.

For PCBP1 and PCBP2, paralogs with previously reported interactions (Zhao et al. 2022), we noticed that the percentage of motif instances overlapping and co-occurring were strongly asymmetric, with 18.78% and 24.29% of PCBP1's motif instances overlapping and co-occurring, respectively, with PCBP2's, while only 2.11% and 3.68% of PCBP2's motif instances overlapped and co-occurred, respectively, with PCBP1's (**Supplemental Table S13B**). We inspected the size of their respective datasets and observed that PCBP1's dataset was much smaller than PCBP2's, with 1,061 motif instances being discovered for PCBP1 compared to PCBP2's 9,240 (**Supplemental Table S13B**). We hypothesize that this asymmetry, in which PCBP1 has fewer motif instances but a larger proportion overlapping or co-occurring with PCBP2's, may indicate a skewed relationship in which PCBP1 relies on PCBP2 to a greater extent than the other way around. This line of reasoning can be extended to other similar cases in which the degrees of overlap and co-occurrence demonstrate asymmetry. We also noticed that PCBP1 had a higher cross-prediction performance on PCBP2's contexts (AUPR = 94.79%) than on its own (AUPR = 90.92%), suggesting that the salient features within PCBP1's contexts that were learned by its cognate model are very strong, as well as highly similar to the salient features present in PCBP2's contexts (**Supplemental Table S13B**). In addition, while our interaction discovery method classifies this pair as a cooperative interaction corresponding to Path I (**Supplemental Note S9**; **Supplemental Table S14**), we note that the degree of motif overlap is relatively close to the degree of motif co-occurrence for each RBP, which may suggest some proclivity for either a cooperative or competitive binding mode represented by Path II. Using other RBPs from **Figure 6E** as a reference, we additionally verified PCBP2-PTBP1 (**Supplemental Table S13C**; Path III, **Supplemental Table S14**), whose interactions have also been previously reported (Kim et al. 2000), and we hypothesize that this pair interacts cooperatively along Path III.

We used the KLD and cross-prediction results to guide our identification of additional positive controls. We noticed that, although their membership to the same cluster depended on the analysis, MATR3 predicted very strongly on PTBP1's contexts (AUPR = 95.75%; **Supplemental Table S13D**), and that the cross-prediction performance was even greater than MATR3's performance on its own contexts (AUPR = 91.08%), suggesting that there were significant patterns within PTBP1's contexts that are recognized by MATR3. Indeed, an interaction between MATR3 and PTBP1 has been reported to co-regulate splicing repression for a subset of MATR3 targets (Coelho et al. 2015). Based on our discovered motifs and contexts, we found that the co-occurrence between MATR3 and PTBP1 motif instances exceeds their overlap considerably (**Supplemental Table S13D**), thus supporting this established cooperative interaction; we add that the interaction may be more likely to occur in cooperative binding modes corresponding to Path I (**Figure 7B**; **Supplemental Table S14**).

As a final example, MATR3 and SUGP2 caught our attention due to their consistent clustering in all analyses performed (**Figures 3**, **4**, **6C**), even being the sole constituents of their cross-prediction cluster (**Figure 6C**), which suggests that their respective global binding profiles are significantly similar to one another while simultaneously being distinct from those of all other RBPs. These two RBPs both have binding sites enriched in antisense L1 elements (Van Nostrand et al. 2020b) and sense L2 elements (Attig et al. 2018), and multiple experimental studies have established SUGP2 as a strong interactor of MATR3 (Chi et al. 2018; Iradi et al. 2018), although the precise role of their interaction has yet to be determined. The asymmetry in their cross-prediction scores (**Supplemental Table S13E**), with SUGP2 performing noticeably stronger on MATR3's contexts (AUPR = 91.58%) than vice versa (MATR3 on SUGP2's contexts, AUPR = 78.34%), and even higher than on its own contexts (AUPR = 82.27%), suggests that MATR3's contexts contain a subset of SUGP2's contexts that have stronger saliency, while SUGP2's contexts as a whole contain more variability. This in turn may suggest a relationship of dependence in which at least some of MATR3's binding and functions are not only enhanced but perhaps even contingent on the presence of SUGP2, while SUGP2's regulatory activities may be more widespread and independent of MATR3. Interestingly, a similar degree of co-occurrence and overlap was observed for their respective motifs (**Supplemental Table S13E**), suggesting that this interaction has an equal propensity to occur both in cooperative modes corresponding to Path I and in cooperative and/or competitive modes corresponding to Path II (**Figure 7B**; **Supplemental Table S14**).

In short, we demonstrate with these positive controls that our proposed method successfully identifies known RBP-RBP interactions. These examples also demonstrate the combinatorics of RBP-RBP interactions (Khoroshkin et al. 2024), with certain RBPs (e.g., PCBP2, PTBP1, MATR3) being involved in multiple different interaction pairs, thus pointing to the exponential scale at which the full range of human RBPs interact.

### SUPPLEMENTAL NOTE S11

#### On the advantages of the RBP-RBP interaction discovery method

We performed a comparative analysis between our approach and a solely peak overlap-based approach, such as in (Van Nostrand et al. 2020b). To this end, we computed the (bidirectional) percentage of eCLIP peak overlaps for each of the 2,485 RBP-RBP pairs in our HepG2 dataset, using the same conditions for overlap as in (Van Nostrand et al. 2020b) (i.e., an overlap of at least 1 nucleotide). We considered an RBP-RBP pair to be a potential interaction if it had non-zero percent peak overlap values in both directions. Because we are not yet considering homotypic interactions in our framework, we removed this class of interactions from our analyses.

Next, we performed an assessment of the annotated interactions in the BioGrid database by cross-referencing it with both the list of known human RBPs (Gerstberger et al. 2014) and the list of RBPs in our eCLIP dataset, finding that, of the 877,535 total human protein-protein interactions in the database, 60,495 constituted RBP-RBP interactions. Of these, there were 436 interactions for which both RBPs in each pair had eCLIP data in our HepG2 dataset. Therefore, we cross-referenced our discovered interactions with this set of 436.

With our framework, we discovered a total of 796 interactions, representing 32.03% of the total 2,485 pairs in our dataset. With respect to the 436 interactions in the BioGrid database, 145 were discovered by our approach (33.26%), while 291 were not (66.74%). Conversely, the peak overlap-based approach resulted in a total of 2,360 interactions, representing 94.97% of the total 2,485 pairs in our dataset, and discovering 409 of those in the BioGrid database (93.81%) while not discovering 27 (6.19%).

We have previously described the importance of cross-prediction scores as a measure for feature similarity, as well as our definition of what constitutes a good cross-prediction score. We sought to understand the differences in cross-prediction observed for the interaction pairs detected by our framework and by the peak overlaps-based approach. We found that the cross-prediction scores for our discovered interactions were significantly greater than the cross-prediction scores for the interactions discovered by peak overlaps (**Supplemental Figure S17A**). In fact, the median cross-prediction score for interactions discovered by peak overlaps was less than 0.6, which lies outside of our definition of good cross-prediction and thus demonstrates that the majority of the interactions proposed by this method have very low feature similarity.

We also sought to understand how the variability in eCLIP peak length affects peak overlaps and the resultant detection of proposed RBP-RBP interactions. When assessing the peak length across all RBPs in our HepG2 dataset, we found significant variability in peak length, ranging from 1 nt to 432 nt, and with a mean length of ~62 nt (**Supplemental Figure S17B**). We next surveyed the distribution of peak lengths for each RBP individually (**Supplemental Figure S17C**), again observing significant variability in peak length across the different RBPs. In short, this peak variability can drastically affect the outcomes of peak overlap-based interaction discovery: longer peaks may increase sensitivity by resulting in more peak overlaps and therefore a potential overestimation, while shorter peaks may decrease sensitivity by resulting in fewer peak overlaps and therefore a potential underestimation. In our work, we utilize a standardized context length of 55 nt to guide the motif overlap and motif cooccurrence analyses that underlies our interaction discovery. This length standardization (1) ensures that the context of binding sites are considered in interaction discovery, (2) guarantees that the considered context length is the same for all RBPs in our analyses, and (3) reduces any bias that is introduced by peak length variability, which can affect each RBP differently.

Given these results, we would like to address important differences in our framework relative to a peak overlap-based approach. First and foremost, the peak overlap-based approach operates under a very strong assumption that the presence of any amount of peak overlap, no matter how small, is enough to support a potential interaction between two RBPs. However, it remains a major open question as to whether or not peak overlaps alone are sufficient to constitute an interaction. Using peak overlaps, we discovered a possible interaction for 94.97% of the 2,485 total RBP-RBP pairs in our dataset, which is an unrealistically large percentage and is highly likely to contain false positives.

While we agree that peak overlaps have the potential to be involved in detecting RBP-RBP interactions, we do not believe that peak overlaps should be the sole condition for detection, as it will either over- or underestimate the predictions. Importantly, a peak overlaps-based approach lacks the details and resolution that is needed to understand the specifics of the interaction, including information associated with the motif and contexts as well as the type of interaction. Thus, while the analysis in (Van Nostrand et al. 2020b) can be considered a precursor to our work, we note that our framework introduces many important advantages over the use and reliance on peak overlaps alone.

We elaborate on these aspects under the following attributes, which are necessary for a good framework:

- **Systematic and structured**: We present a formalized framework with a well-defined goal that follows a structured, logical, and consistent approach and is presented as an organized step-by-step pipeline.
- **Comprehensive**: Our framework consists of a complete pipeline that contains all of the necessary methods and algorithms to perform analyses and to draw conclusions from our results.
- **Clear**: Our framework consists of phases that are defined and well-organized, with a clear and logical progression between each phase, thus enabling seamless connection, continuity, and a clear path for testing, analysis, and inference. Our framework also relies on clearly defined assumptions.
- **Informative**: In our study, we introduce a comprehensive framework that enables RBP-RBP interaction discovery that has higher resolution than a solely peak overlap-based method.
    - Our discovery and utility of motifs and contexts significantly improves the **resolution** of RBP-RBP interaction discovery. Using peak overlaps alone allows only for a binary classification of any RBP-RBP pair as interacting or not interacting. However, the sole reliance on peak overlaps does not provide any motif or context information, without which there is no power to further classify hypothesized interactions into a specific category, such as cooperative or competitive. In our approach, we leverage our discovered motifs and contexts to measure the degree of motif overlap and motif co-occurrence, each of which provides additional information to hypothesize the specific interaction mode of the participating RBPs; further, it allows us to localize precisely where an interaction occurs. We consider this to be one of the major advantages of our approach.
    - Our method preserves bidirectionality, in that we consider the asymmetry of each feature (motif KLD, context KLD, cross-prediction, motif overlap, motif co-occurrence) used in our interaction discovery approach. We hypothesize that this asymmetry is an important property that is directly informative of the dependency and extent of interaction between two RBPs. For example, only 3.323% of BCLAF1's peaks overlap TRA2A's peaks, but 73.72% of TRA2A's peaks overlap BCLAF1's peaks; this may suggest that BCLAF1 has more broad functions independent of interaction with TRA2A, but TRA2A has more specific functions that rely on interactions with BCLAF1. A peak overlaps-based approach such as in (Van Nostrand et al. 2020b) loses this critical information, as it reduces the bidirectional peak overlap to a single value by taking the maximum, which limits its ability to make such hypotheses about interaction directionality.
- **Flexible**: Our approach is amenable to expansion (i.e., for the development of new methods and algorithms for further analysis and validation), extension (see below), and generalization (see below).
- **Extendable**: Because our approach for proximal interaction discovery identifies motifs and contexts that are implicated in the interactions, our framework can be further extended to also consider distal interactions; this is, in fact, one of our future directions.
- **Generalizable**: Our framework allows for greater generalizability of interaction discovery to other applications, especially when eCLIP peak data is unavailable. For example, given an RNA sequence of interest that is implicated in a certain biological process and that is hypothesized to be regulated by an interaction between RBPs, we can utilize our motifs and contexts to determine which RBPs have binding sites in that sequence. When peak data is absent, this analysis would not be possible.

In summary, despite the apparent disparity in the interaction discovery rates between our framework and the peak overlap-based approach, we remain confident that our framework provides multiple important and necessary advantages for RBP-RBP interaction discovery. A reliance on peak overlaps alone depends on an ambiguous assumption that peak overlaps are sufficient to suggest the presence of RBP-RBP interactions, which has not yet been established and can result in the detection of false positives, as we have previously described in reference to the very high percentage of discovered interactions. In our approach, we devise and introduce multiple new measures that help to provide higher confidence of RBP-RBP interaction discovery. We defined a relationship between distribution and feature similarity that guarantees an interconnection between distribution and feature prominence, and this helps to increase confidence and prediction accuracy. The symmetrical and asymmetrical nature of these measures is also advantageous, as it is informative of the dependence relationship between RBPs. Here, we present a clear, structured, systematic, and comprehensive framework to discover RBP-RBP interactions that provides many advantages that highlight the power of our framework over a peak-based approach due to its integration of RBP motifs, contexts, and feature similarities, including high resolution, informativeness, flexibility, extendibility, and generalizability.

### SUPPLEMENTAL NOTE S12

#### Additional comments on the proposed AKAP1-UPF1 interaction

**On additional support for the proposed interaction**

Although an interaction between AKAP1 and UPF1 has briefly been reported (Gabrovsek et al. 2020), the underlying mechanism has not yet been resolved. We sought to understand how this RBP pair may cooperate. AKAP1 – a member of the A-kinase anchoring protein (AKAP) family best known for its localization of protein kinase A (PKA) to different cellular compartments to facilitate signaling pathway propagation (Turnham and Scott 2016) – spatially regulates (with the help of LARP4 and PABPC1) the local translation of essential mitochondrial proteins at the mitochondrial outer membrane (Gabrovsek et al. 2020; Sharma and Fazal 2024). AKAP1 has also been shown to bind ribonucleoprotein (RNP) complexes with roles in translation and mRNA decay through its KH and Tudor domains (Gabrovsek et al. 2020). On the other hand, Up-frameshift 1 (UPF1) is an RNA helicase with numerous functions in mRNA decay pathways, the most prominent of which is nonsense-mediated decay (NMD), for which it is an essential RBP (Kim and Maquat 2019). When a premature termination codon (PTC) in an erroneously transcribed RNA is detected by the ribosome, the mRNA is targeted for degradation when UPF1 outcompetes PABPC1 (a translation-promoting factor) for binding to the eRF1-eRF3 complex (Qi et al. 2022); this in turn prevents efficient translation termination from occurring, and an activated UPF1 triggers the NMD pathway upon its phosphorylation by the kinase SMG1 (Kim and Maquat 2019).

We investigated the genomic regions containing overlapping AKAP1-UPF1 contexts, noting that the vast majority (93.3%) occurred in 3'UTRs, which are known to have roles in mRNA translation regulation (Mayr 2019). As expected, a number of these overlapping contexts occurred in genes encoding important mitochondrial proteins, such as *SDHAF1* (a component of succinate dehydrogenase, which is involved in mitochondrial metabolism), *SOD2* (oxidative stress reducer), *MRPL12/49* (two mitochondrial ribosomal proteins), *SLC25A1/10/42* (three solute carriers regulating small molecule transport across mitochondrial membranes), *TIMM13/50* and *TOMM34* (three translocases regulating protein import into the mitochondria), and numerous other genes encoding proteins that are either components of or localized to the inner or outer mitochondrial membrane (e.g., *ATAD3B*, *BCL2L1*, *MFN2*, *MUL1*, *RHOT2*, and *TMBIM6*) (Ginsberg et al. 2003; Gabrovsek et al. 2020; Rath et al. 2021; Sayers et al. 2022). Finally, we performed gene ontology (GO) analysis (Kolberg et al. 2023) on the gene set containing AKAP1-UPF1 context overlaps (**Figure 7D**; **Materials and Methods**). Despite the lack of enrichment for mitochondria-specific GO terms, the recurrence of significantly enriched localization (e.g., protein localization), membrane (e.g., endomembrane system organization), and transport (e.g., vesicle-mediated transport) terms led us to hypothesize that AKAP1's functions of local protein synthesis may also occur at components of the endomembrane. Indeed, we found many genes in the set coded for proteins with roles in the endoplasmic reticulum (ER) membrane (e.g., *ABHD4*, *DHCR7*, *ESYT1*, *LPCAT3*, *MPDU1*, *NCLN*, *POR*, *REEP6*, *RRBP1*, *TRAM2*), Golgi membrane (e.g., *CDIPT*, *CHPF*, *FUT6*, *MYO18A*, *RFNG*), trans-Golgi network (e.g., *AP1M1*, *ATG9A*, *TGOLN2*), and ER-to-Golgi transport (e.g., *ACTR1A*, *GET4*, *RAB1B*, *TEX261*, *YKT6*) (Sayers et al. 2022).

**On the bidirectional cumulative KLDs between AKAP1 and UPF1**

We demonstrate the interpretation of the cumulative KLDs for the cited example in our proposed interaction between AKAP1 and UPF1. Let the PPMs for AKAP1 represent the true distributions and the PPMs from UPF1 represent the approximating distributions. Then, the cumulative motif KLD(AKAP1, UPF1) = 31.36, whereas the cumulative context KLD(AKAP1, UPF1) = 33.32. To demonstrate the meaning of these values, we note the following 2 properties. First, the cumulative motif KLD of 31.36 represents a sum across 5 positions of the motif; therefore, on average, each position within the motif has a KLD of 6.272 (i.e., 31.36 / 5), which is not only considerably larger than zero, but also outside of our defined upper bound of 1.07, and thus indicates that (on average) each position of UPF1's motif is dissimilar to the corresponding position of AKAP1's motif. This underlies our statement that the binding motifs between the 2 RBPs are dissimilar.

Regarding the second property: the cumulative context KLD of 33.32 represents a sum across all 55 positions of the full context, including the 25 positions of the left context, the 5 positions of the motif, and the 25 positions of the right context. To derive the cumulative KLD across only the flanking positions, and independent of the motif, we subtract the cumulative context and motif KLDs (33.32 - 31.36), arriving at a cumulative KLD of 1.96 across all 50 positions surrounding the motif to each side. In other words, the cumulative KLD between motifs and contexts only increased by a small amount (i.e., 1.96), despite representing a sum across a much larger number of positions (i.e., from 5 to 55). Therefore, on average, each position of the flanking contexts has a KLD of 0.0392 (i.e., 1.96 / 50), which is not only very close to 0, but also within our defined range for high distribution similarity, and thus indicates that (on average) each position of UPF1's flanking contexts are very similar to the corresponding position of AKAP1's flanking contexts. This underlies our statement that the contexts between the 2 RBPs are similar, and the context similarity is further reflected by the strong cross-prediction performance between AKAP1 and UPF1 of 82.25%, thus indicating that AKAP1's and UPF1's contexts have both distribution and feature similarity.

We also find similar results when we conversely designate UPF1's PPMs as the true distributions and AKAP1's PPMs as the approximating distributions: the cumulative motif KLD (UPF1, AKAP1) = 33.64, representing an average KLD of 6.728 over each position of the motif, while the cumulative context KLD (UPF1, AKAP1) = 35.59, representing a small increase from the cumulative context KLD (1.95) and an average KLD of 0.0318 over each of the flanking positions. The context similarity is likewise reflected in the strong cross-prediction performance of 81.51%. While the asymmetry for each feature is apparent, the values in either direction of comparison are very similar to one another, indicating that the observed properties are reciprocated. Therefore, we conclude that the contexts of AKAP1 and UPF1 demonstrate strong and symmetric distribution and feature similarity.

**On additional support from RBP knockdown shRNA-seq data**

To further support this proposed interaction, we also integrated RBP knockdown (KD) shRNA-seq data into our analyses. We quantified differential expression across datasets, and for each dataset, computed the enrichment of discovered contexts in differentially expressed genes relative to background (**Materials and Methods**). To optimize resolution and avoid masking important biological signals, we additionally stratified this enrichment analysis by (1) direction of differential expression (i.e., up- or down-regulated) and (2) by genomic region. When analyzing AKAP1 and UPF1 independently, we found that AKAP1 contexts were significantly enriched in the 3'UTRs of genes that were down-regulated upon its KD (FC = 2.76, *p*-value = 2.7e-67), whereas UPF1 contexts were significantly enriched in the 3'UTRs of genes that were up-regulated upon UPF1 KD (FC = 1.95, *p*-value = 4.71e-23) (**Figure 5A**). To investigate the proposed interaction between AKAP1 and UPF1, we next computed the enrichment of interacting contexts (i.e., the intersection between AKAP1 and UPF1 contexts) in genes that were differentially expressed after both AKAP1 and UPF1 KD. Importantly, we found that interacting contexts were 3.29-fold enriched (*p*-value = 1.94e-46) in the 3'UTRs of genes that were down-regulated upon AKAP1 KD, and 2.18-fold enriched (*p*-value = 1.7e-09) in the 3'UTRs of genes that were up-regulated upon UPF1 KD. These findings demonstrate that the KD of each respective RBP has a significant effect on the expression of genes in which interactions between the two RBPs is implicated.

### SUPPLEMENTAL NOTE S13

#### Additional comments on the proposed DDX59-HNRNPK-PCBP2 interaction

##### Gene Ontology (GO) analysis on pairwise gene sets for all 3 pairs

To investigate the functional implications of this interaction, we first studied each pair of RBPs in this trio independently. We performed intersections between the respective contexts of the RBPs in a pairwise manner, making sure to exclude any resultant intersections that also overlapped the third RBP (e.g., context overlaps between HNRNPK and PCBP2 that do not overlap with DDX59 contexts; **Materials and Methods**). Next, we performed GO analysis on each set of intersections to determine the gene functions that are uniquely regulated by each pair (**Materials and Methods**). As expected, HNRNPK and PCBP2, as hnRNPs with a large breadth of mRNA regulatory functions (Kim et al. 2000; Lang et al. 2021), were enriched for the largest number of unique GO terms ($N$ = 88), which were also very diverse in nature; while some of these terms pertained to more specialized processes (e.g., axonogenesis, miRNA metabolic process, postsynapse organization, tricuspid valve development) or certain signaling pathways (e.g., serine/threonine kinase, Rho GTP-ase, SMAD, semaphorin-plexin), others were more general terms related to cell, tissue, organ, and organism development. HNRNPK and DDX59 were enriched for 12 unique GO terms, which included dendrite morphogenesis, dendritic spine development, and dendritic spine morphogenesis terms, suggesting that these 2 RBPs may have specific, coordinated roles in dendrite development. On the other hand, PCBP2 and DDX59 were enriched for only 3 unique GO terms, 2 of which were related to RNA Pol II transcription and the third of which was "regulation of phosphatidylinositol 3-kinase/protein kinase B signal transduction", which may suggest that they have coordinated roles in the PI3K/AKT pathway. Previous studies have demonstrated associations for HNRNPK in axon (Liu et al. 2008; Liu and Szaro 2011) and dendrite (Proepper et al. 2011; Folci et al. 2014; Leal et al. 2017) development and for PCBP2 in the PI3K/AKT signaling pathway (Qin et al. 2022; Yang et al. 2023) in various organisms. Our pairwise findings implicate novel roles for DDX59 in regulating axon and dendrite development in cooperation with HNRNPK and PI3K/AKT signaling in cooperation with PCBP2 in human.

##### GO analysis on the intersection of gene sets from all 3 RBPs

Because each pair of RBPs within this trio exhibited strong features indicative of cooperative interactions, we leveraged transitive relations to hypothesize that DDX59, HNRNPK, and PCBP2 may also interact in unison to form a complex. To investigate this possibility, we performed a three-way intersection between their respective contexts (**Materials and Methods**) and found that 21.79% of DDX59's contexts overlapped with both HNRNPK's and PCBP2's, with most of these events located in introns or coding sequences. From GO analysis of the genes containing context overlaps between all three RBPs, we discovered enrichment for anatomical structure development and its child term anatomical structure morphogenesis (**Figure 7F**); further inspection of additional child terms for both revealed many related to the development of the face (face, head, & craniofacial suture morphogenesis), oral cavity (hard & soft palate morphogenesis, roof of mouth development, tooth eruption), and digits (limb joint & appendage morphogenesis) (Binns et al. 2009). Mutations in DDX59 have been associated with orofaciodigital syndrome, a disease which leads to varied disfigurements in features of the mouth, face, and digits (Shamseldin et al. 2013; Faily et al. 2017), which may help to explain the discovered GO terms. We additionally looked at terms that were uniquely enriched for genes in the three-way intersection but not enriched in any of the three pairwise GO analyses ($N$ = 12). Along with adherens junction organization, cell-matrix/cell-substrate adhesion, and intracellular signal transduction terms, we found multiple terms related to signal transduction (intracellular signal transduction; chemical synaptic transmission, postsynaptic; regulation of postsynaptic membrane potential; excitatory postsynaptic potential). These latter GO terms reinforce our proposed role of DDX59 in the nervous system, and may suggest that DDX59 cooperates with both HNRNPK and PCBP2 to regulate signal transmission across the synapse.

##### Summary of main findings for interactions between DDX59, HNRNPK, and PCBP2

We posit several hypotheses regarding cooperative interactions, by way of Path I (**Figure 7B**; **Supplemental Table S14**), involving DDX59, HNRNPK, and PCBP2. At the molecular level, we hypothesize novel and specific roles for DDX59 in regulating pre-mRNA splicing, mRNA transport, mRNA stability, and mRNA translation, via its strong association with either one or both of the hnRNPs possessing such functions. At the developmental level, we propose that DDX59, HNRNPK, and PCBP2 work together cooperatively – either in pairs or as a trio – to regulate various aspects of nervous system development and synaptic signaling. Finally, at the phenotypic level, we uncover a potential implication for HNRNPK and PCBP2's involvement in orofaciodigital syndromes, via their strong association with DDX59. It may be that mutations in DDX59 impair its protein-protein interactions with HNRNPK and/or PCBP2, thereby disrupting the joint regulation of shared target genes or signaling pathways that are critical to proper development of the face, oral cavity, and digits. In conclusion, utilizing combinatorics and transitive inference, we propose an interaction between DDX59, HNRNPK, and PCBP2 – either as individual pairs, a complex of three proteins, or both – that holds mechanistic, functional, and disease significance, thereby demonstrating the utility of our approach.
