## Supplementary figures and images for "A novel NLP-based method and algorithm to discover RNA-binding protein (RBP) motifs, contexts, binding preferences, and interactions"

### Supplemental Figure S1

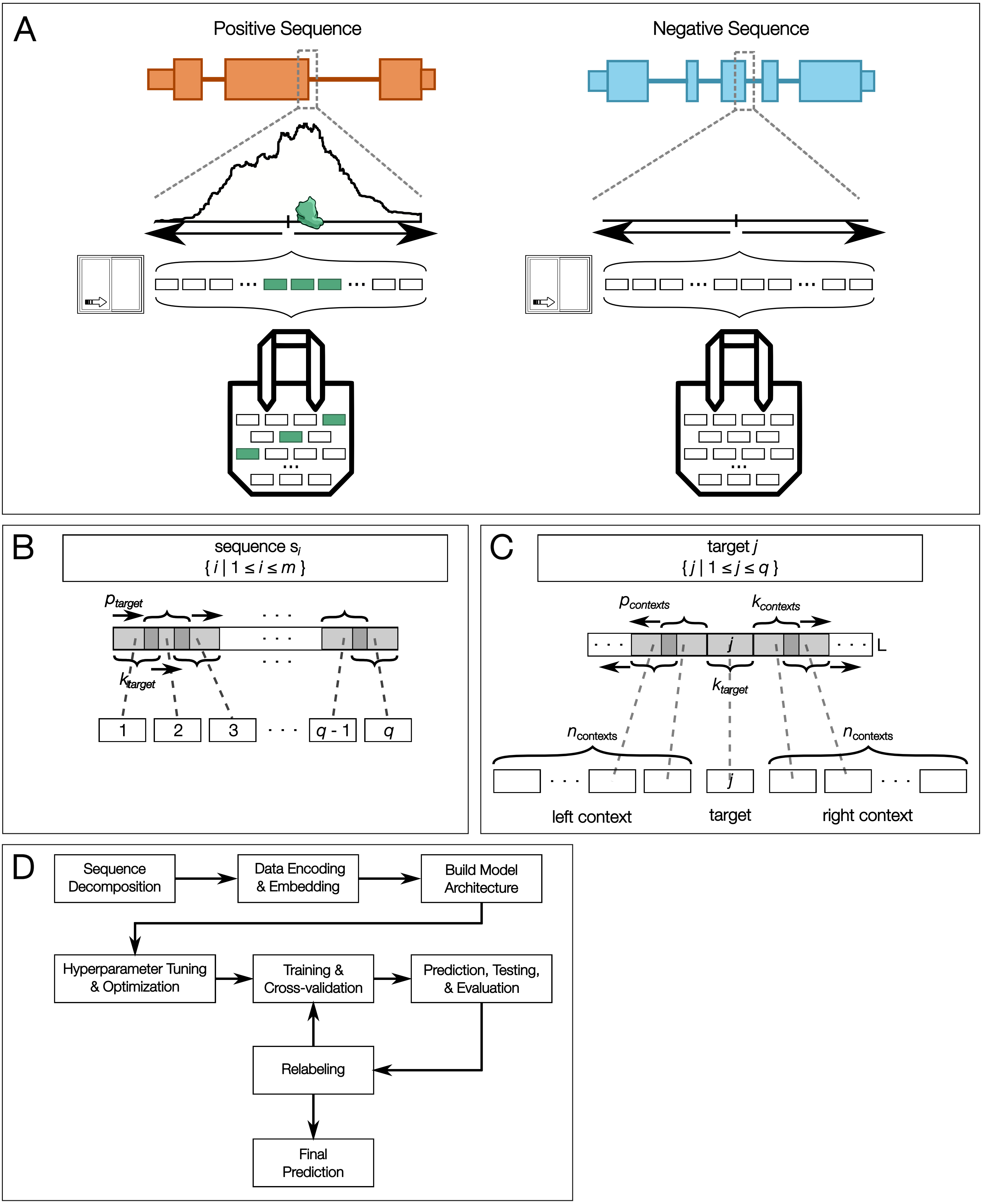


Supplemental Figure S1

### Supplemental Figure S2

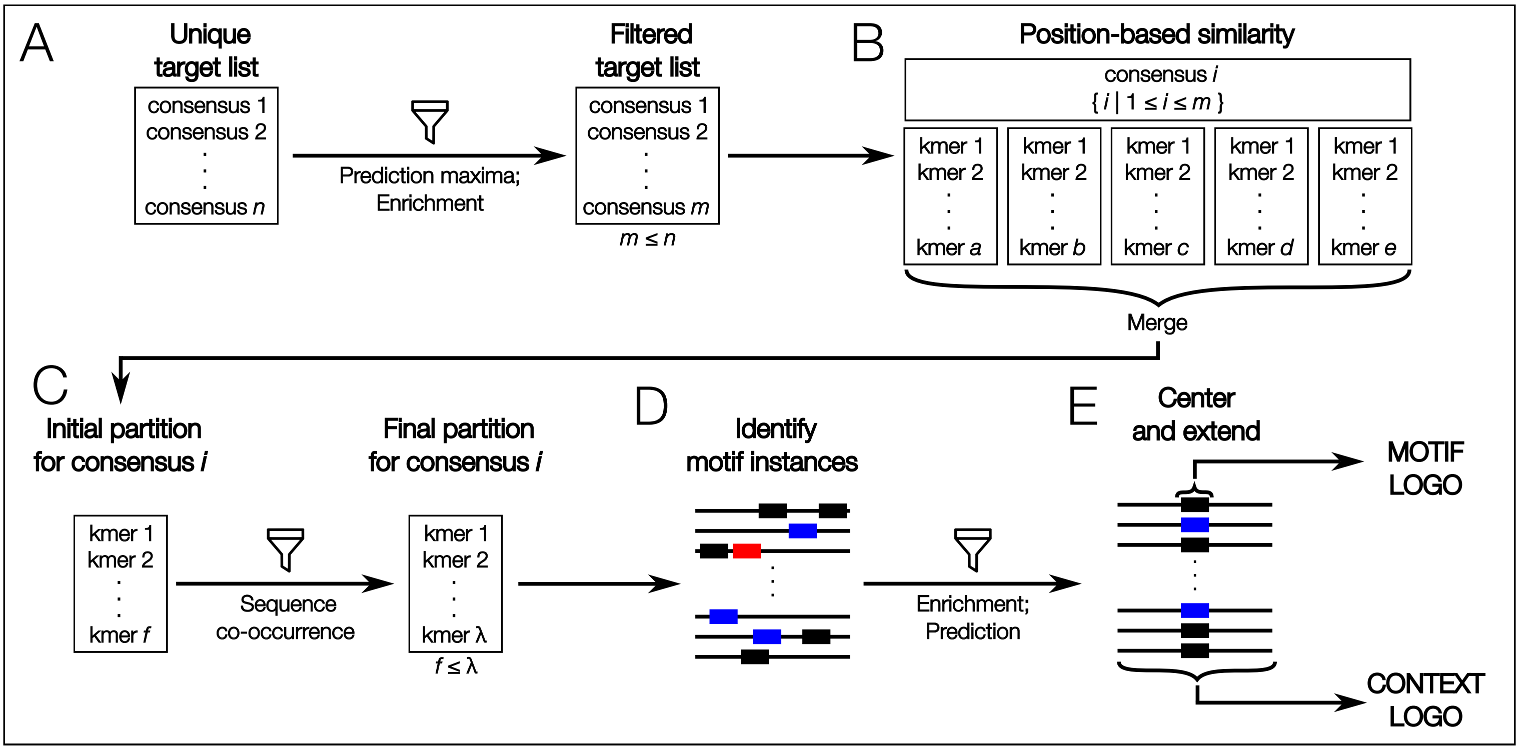

### Supplemental Figure S4

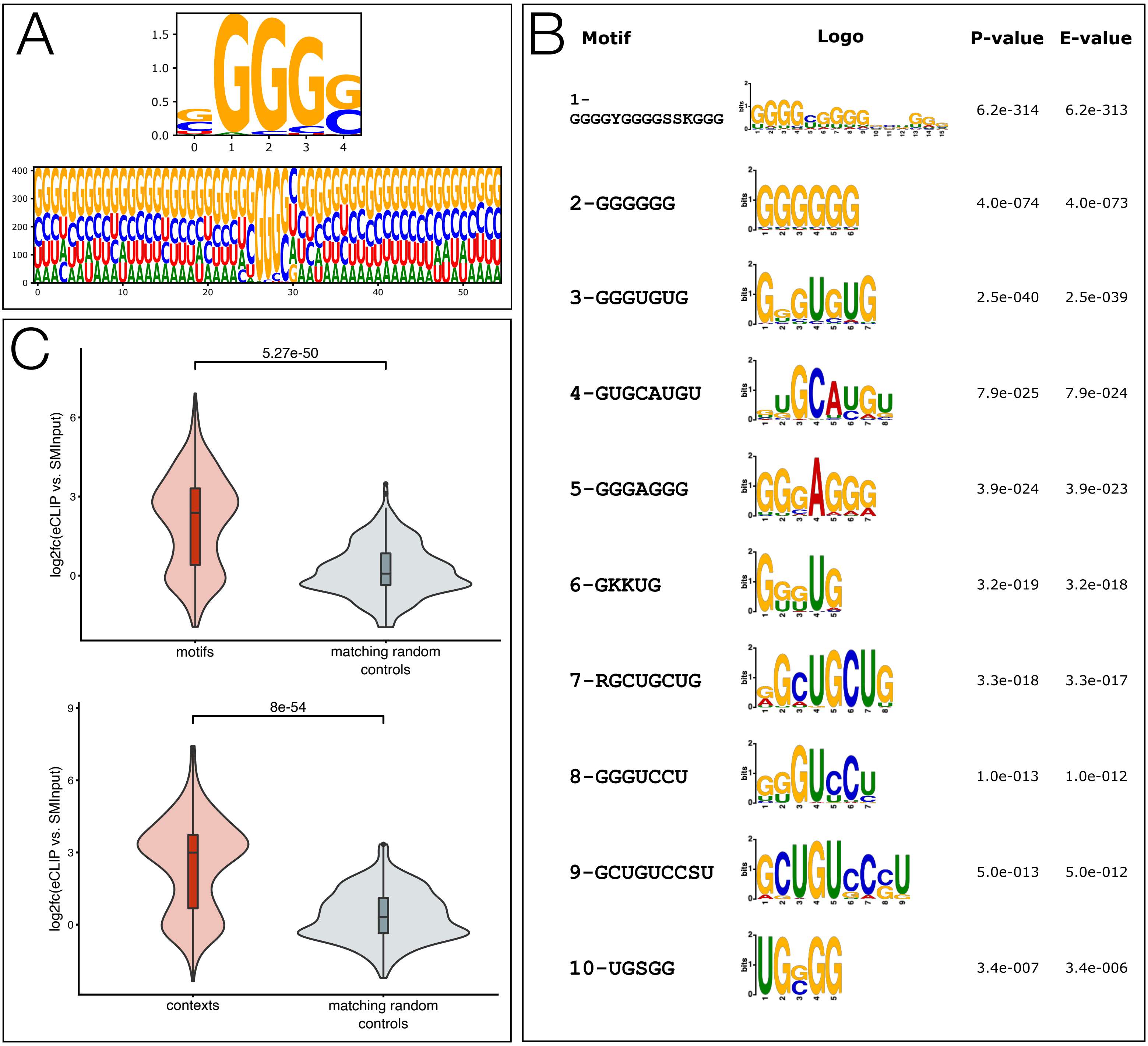


Supplemental Figure S4

### Supplemental Figure S5

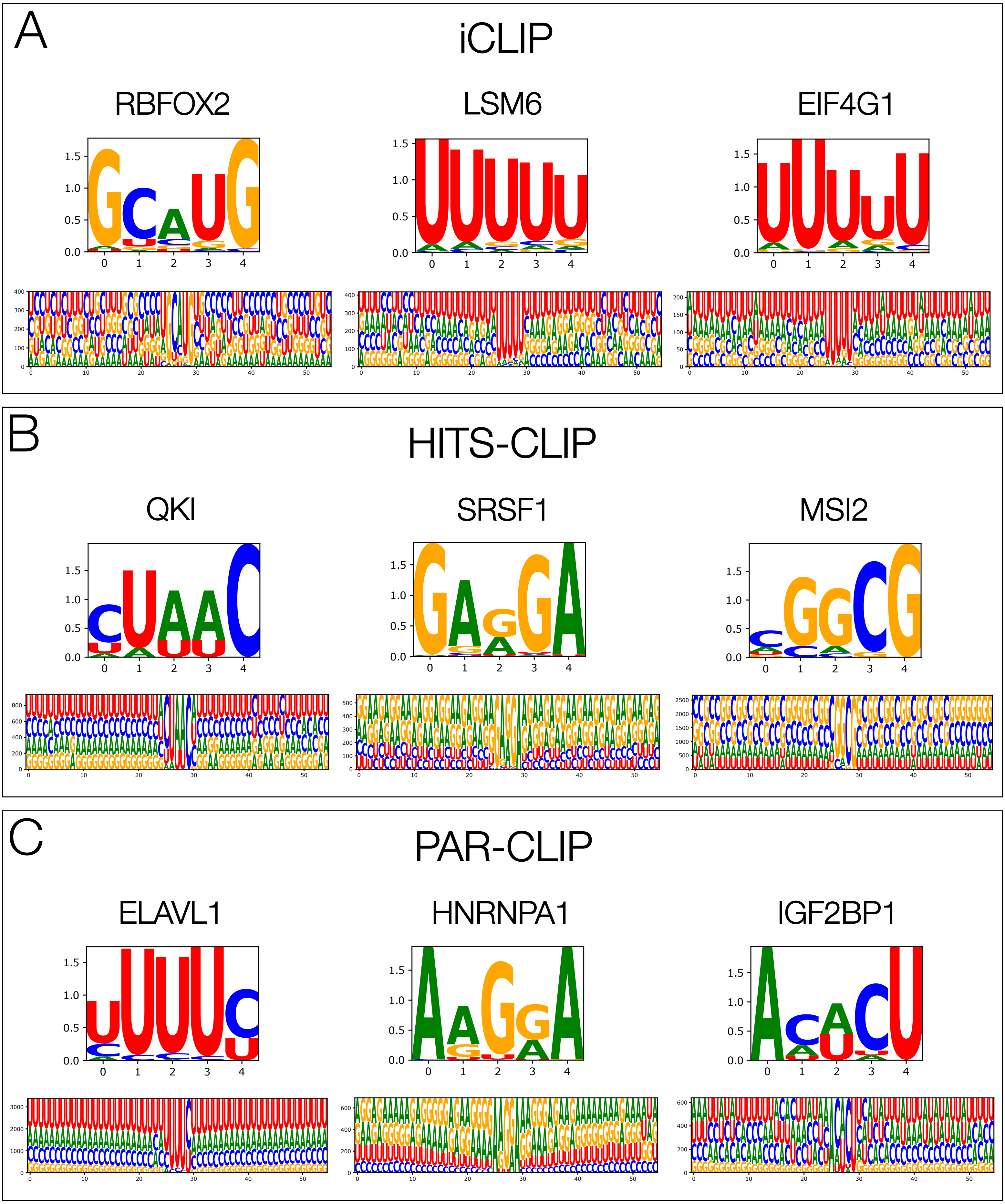

### Supplemental Figure S6

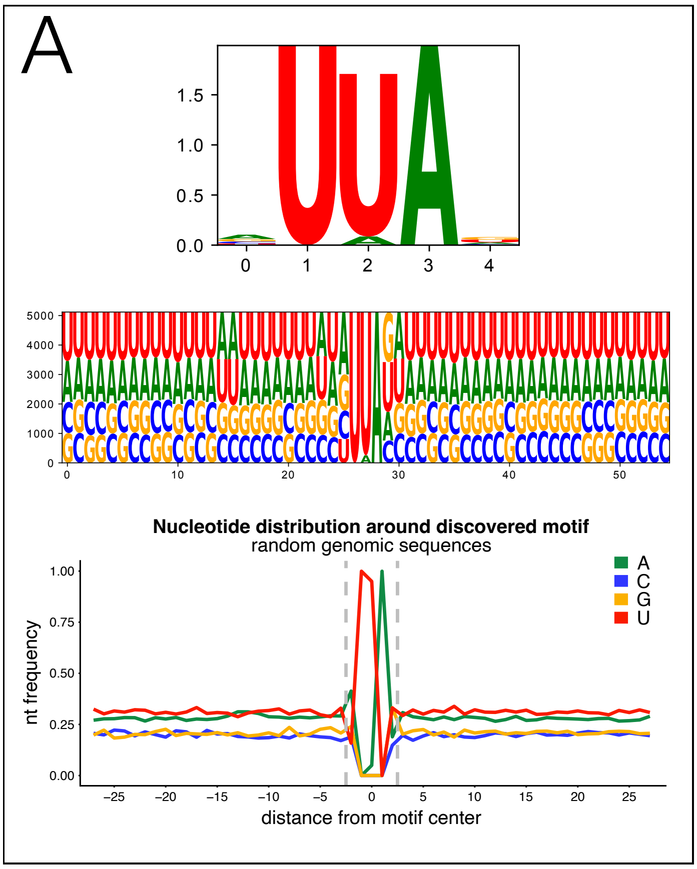

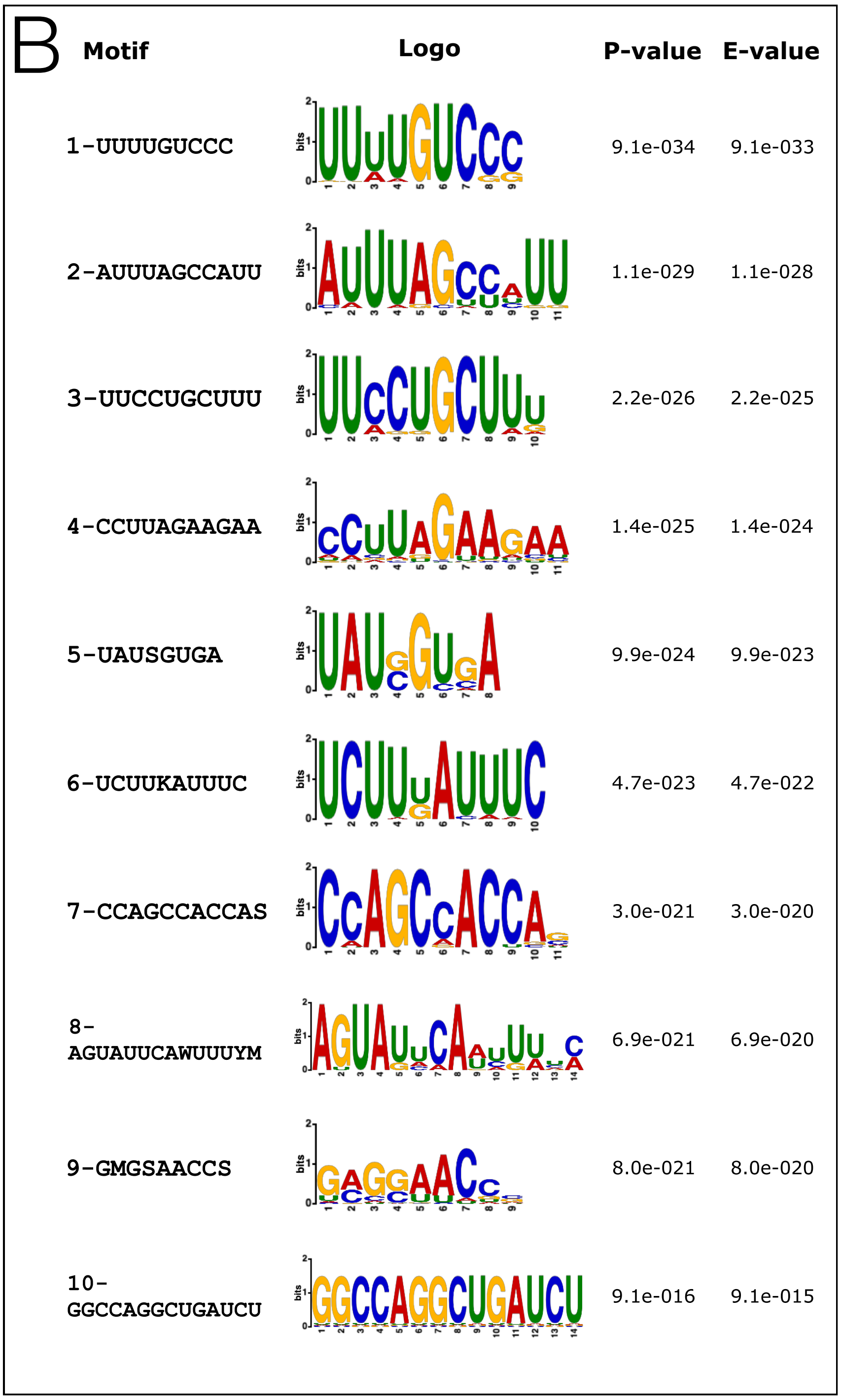

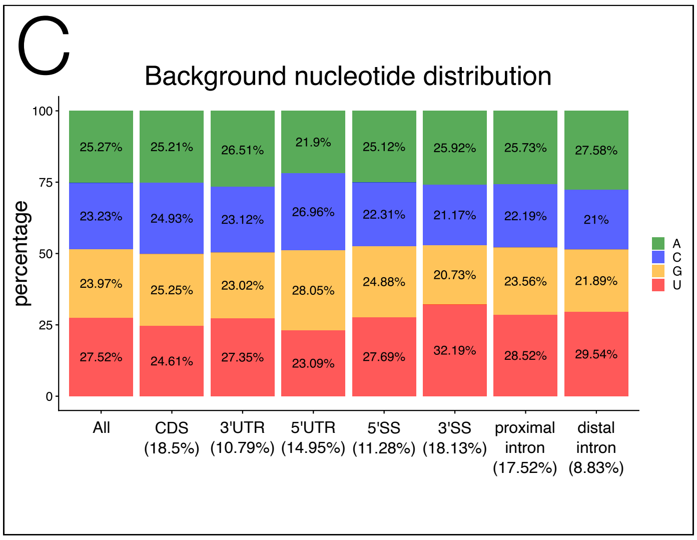

### Supplemental Figure S8

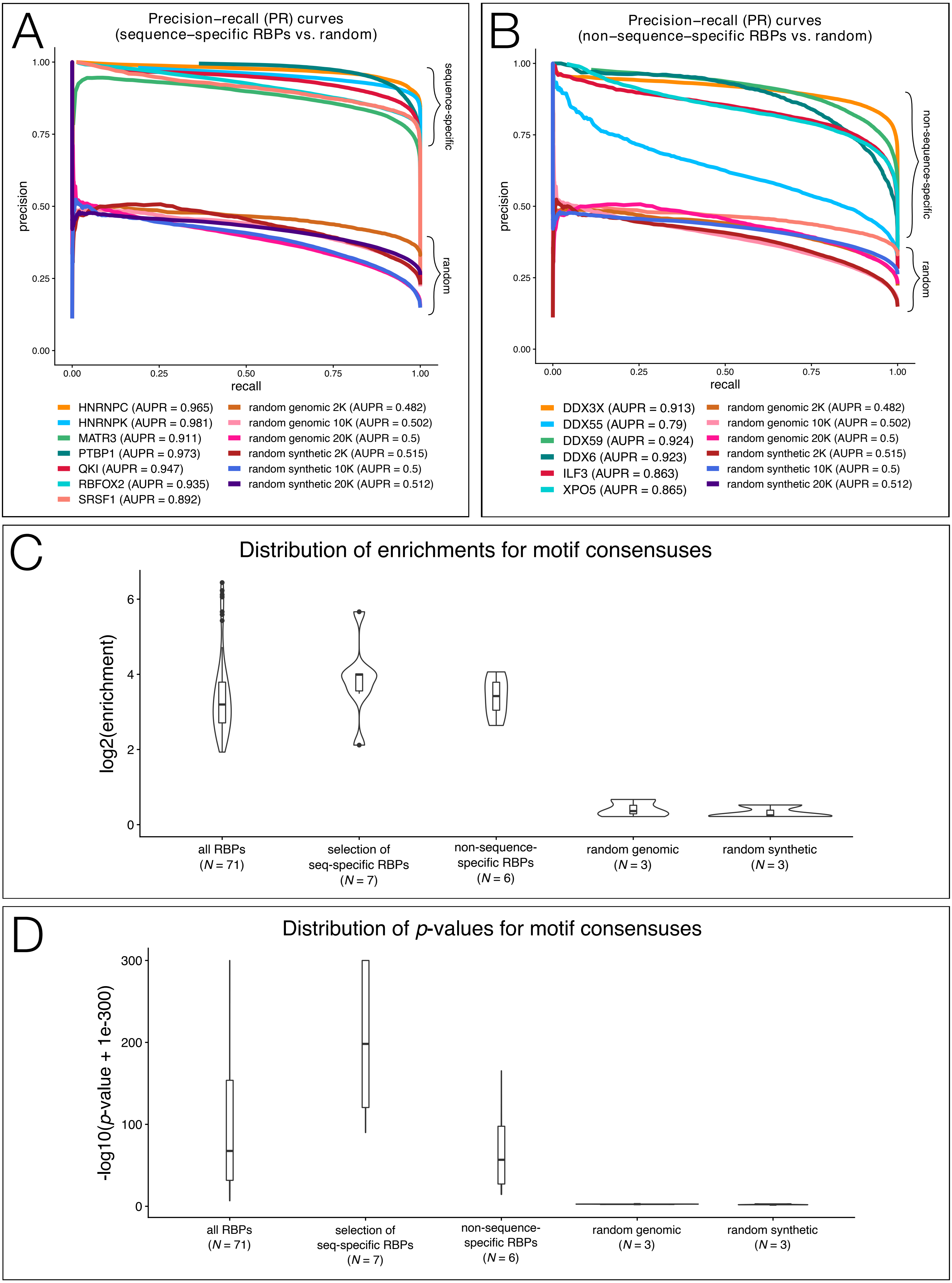


Supplemental Figure S8

### Supplemental Figure S9

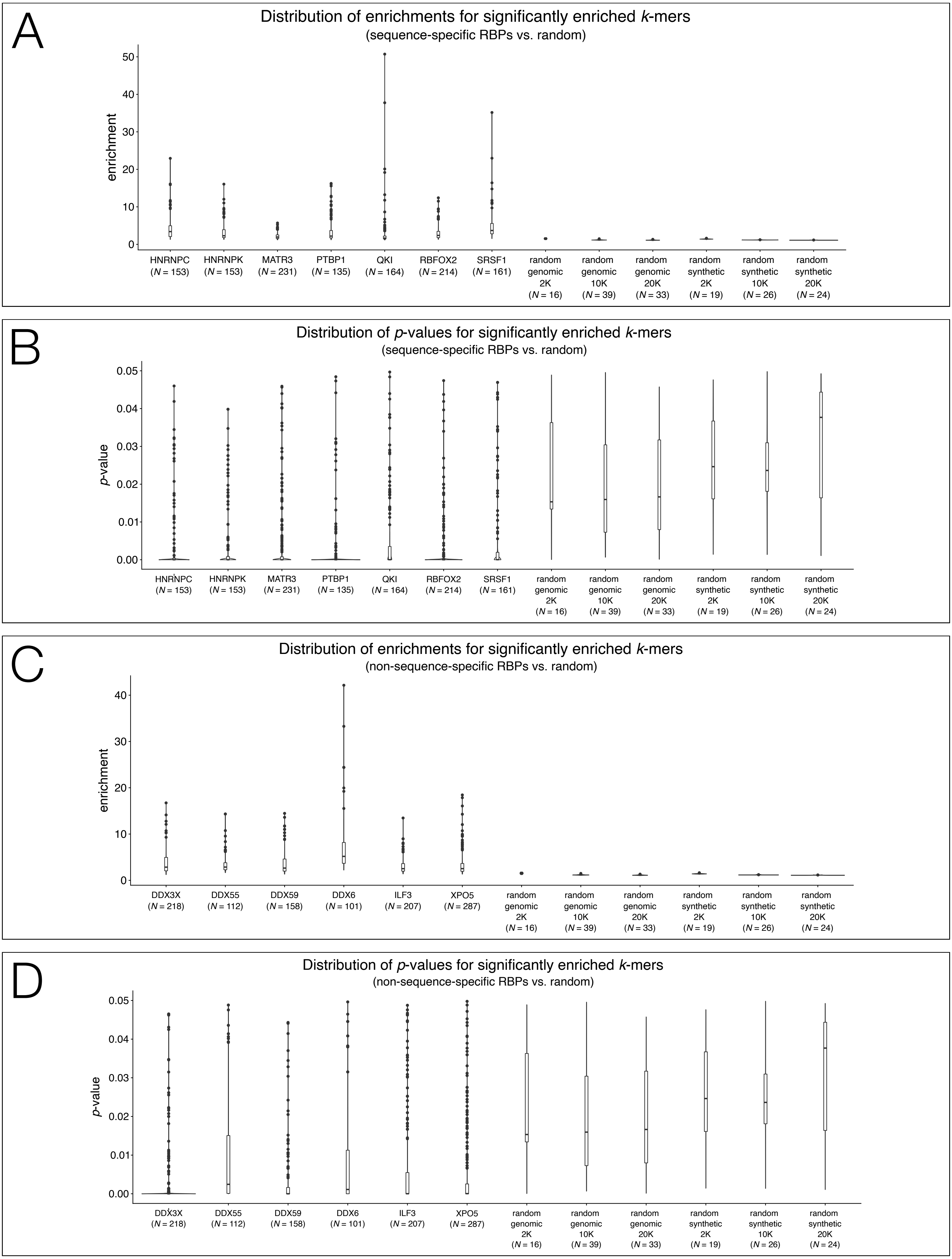


Supplemental Figure S9

### Supplemental Figure S10

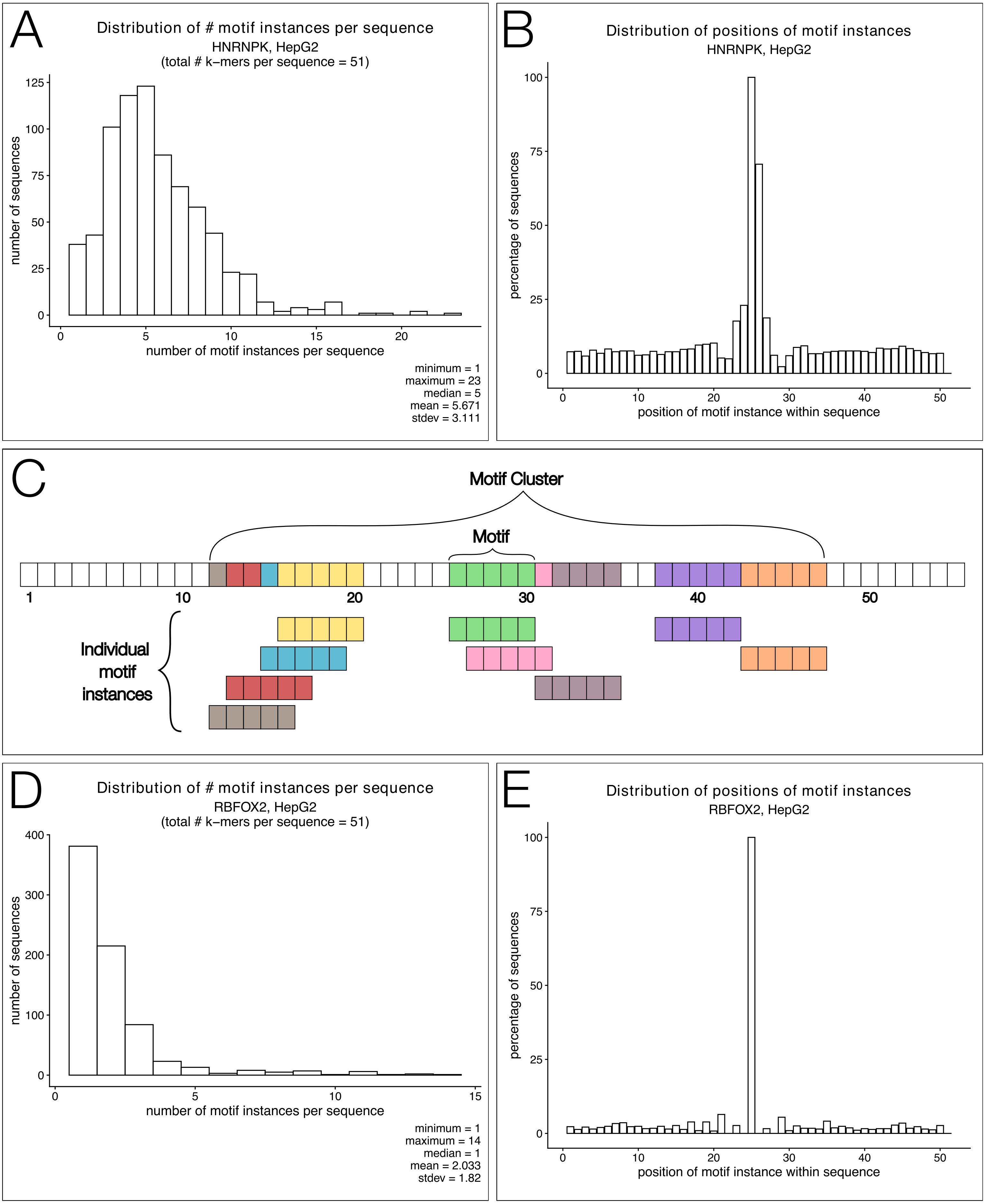


Supplemental Figure S10

### Supplemental Figure S11

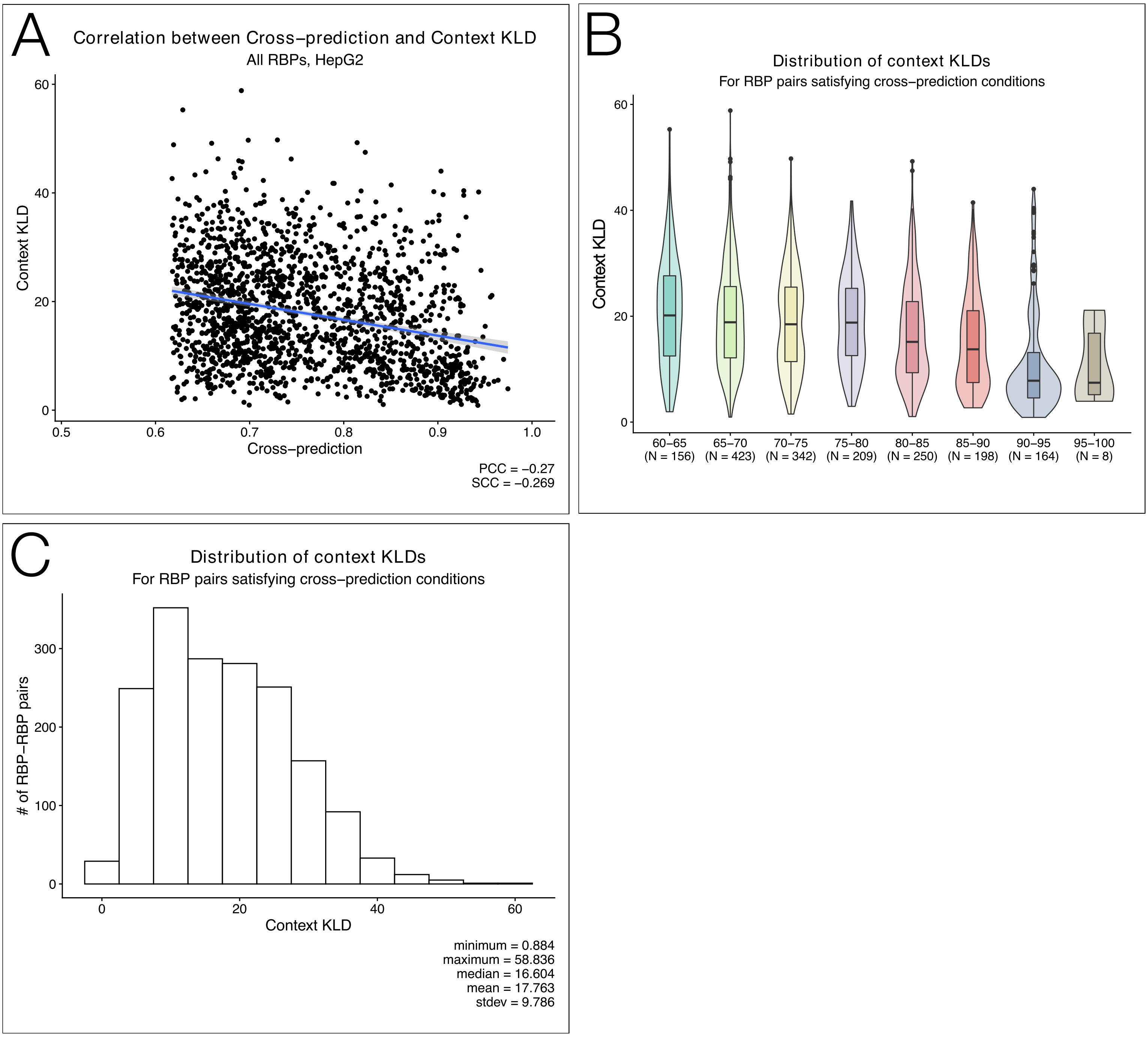


Supplemental Figure S11

### Supplemental Figure S12

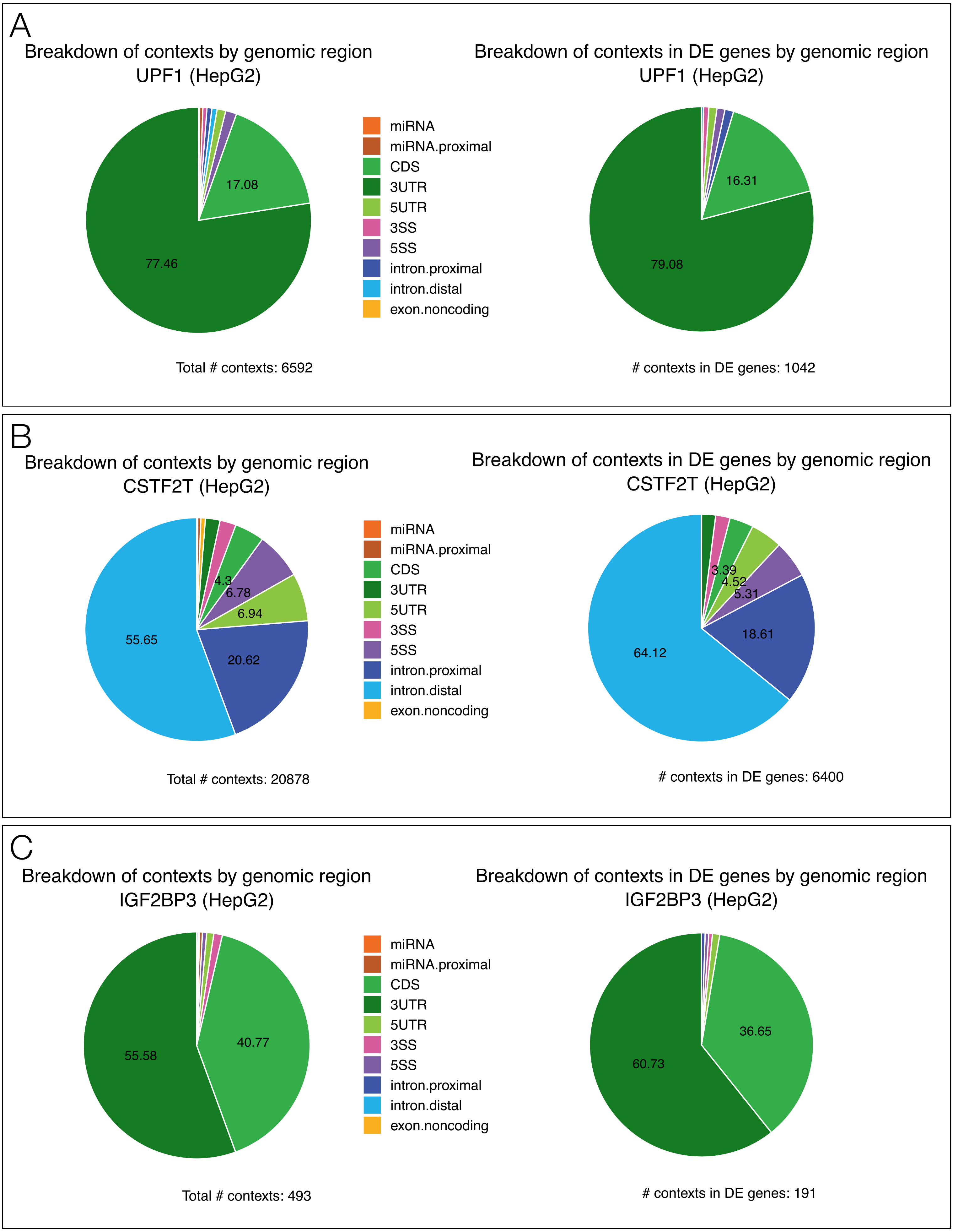


Supplemental Figure S12

### Supplemental Figure S13

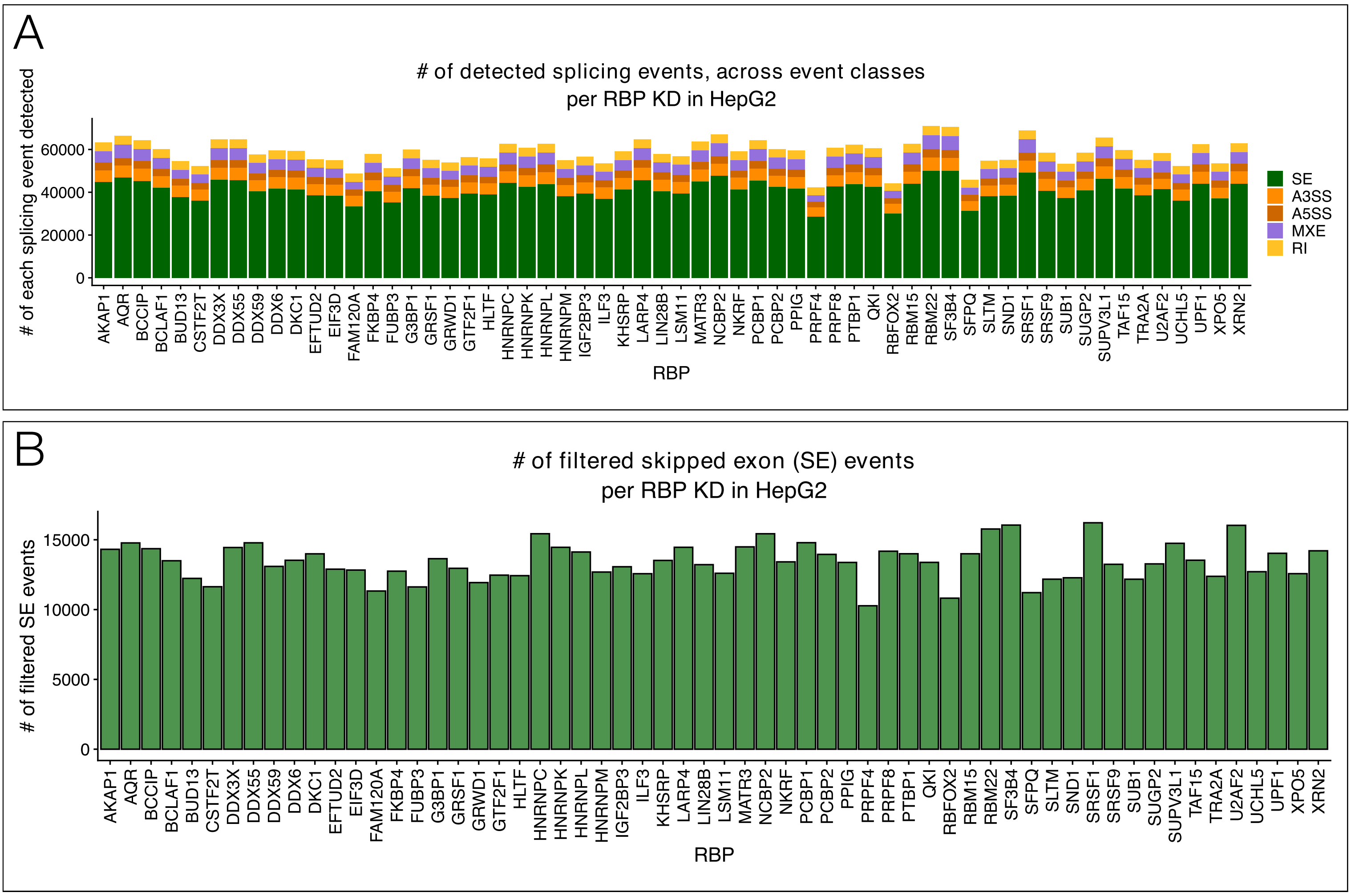


Supplemental Figure S13

### Supplemental Figure S14

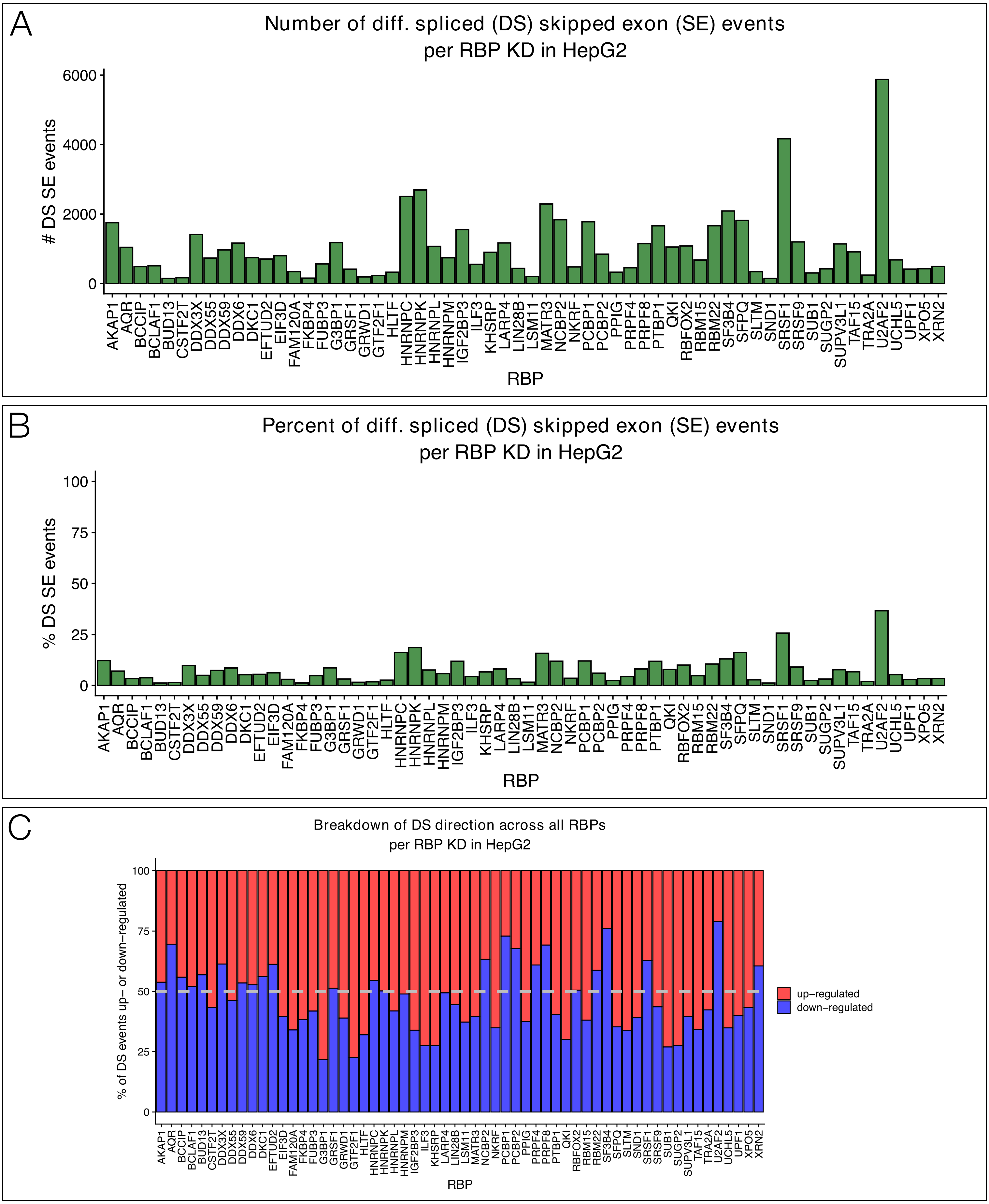

### Supplemental Figure S15

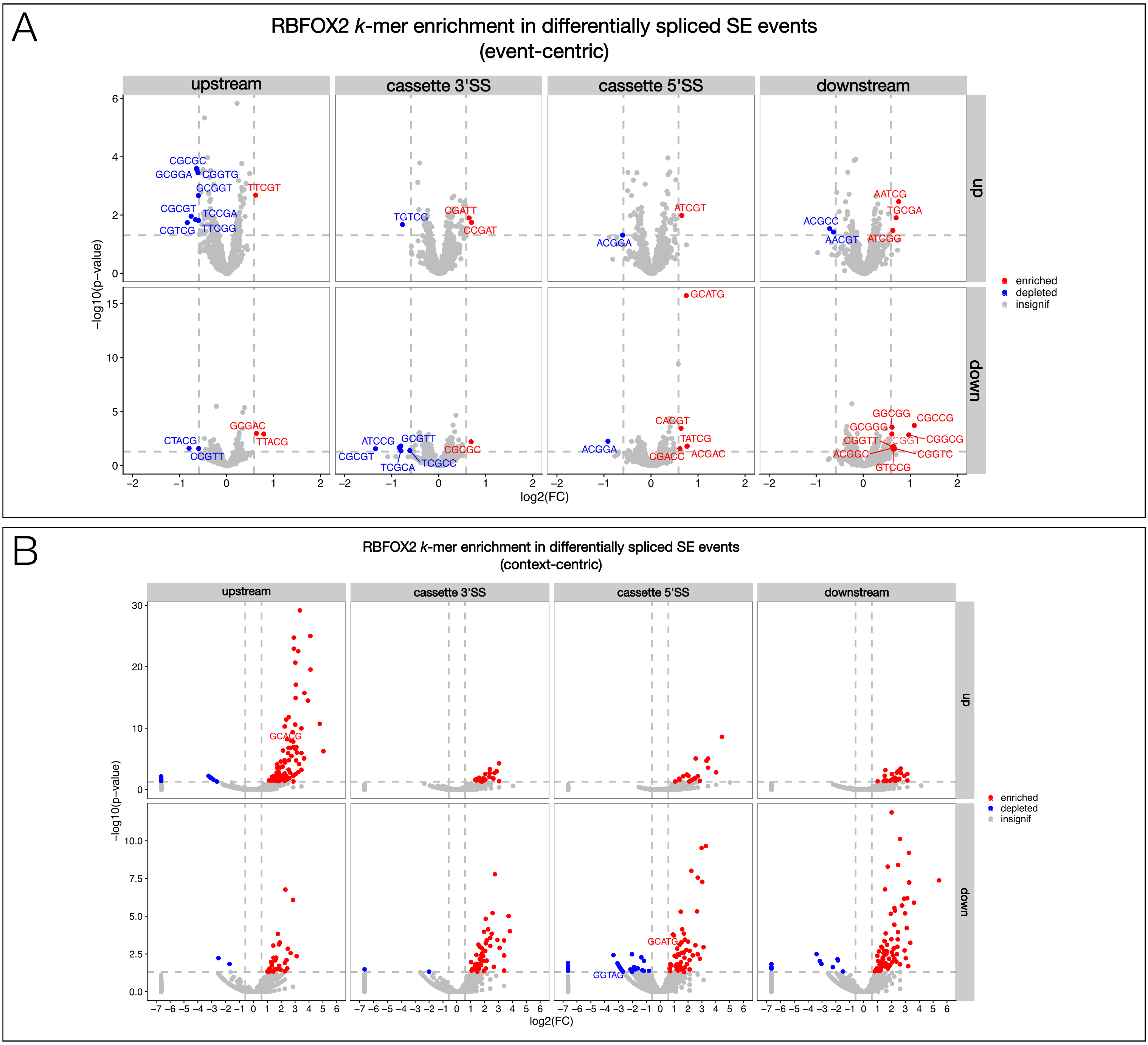

### Supplemental Figure S16

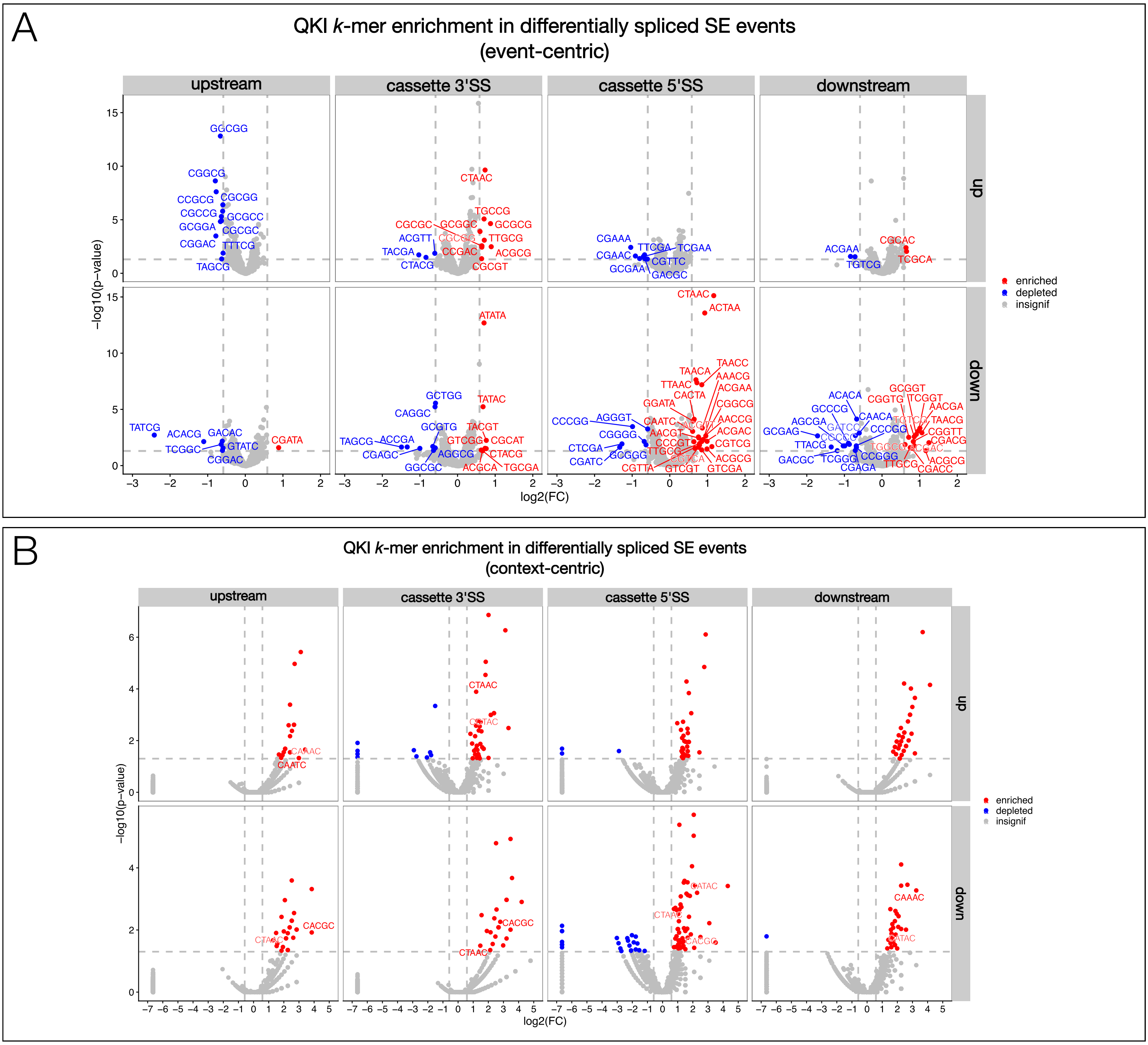


Supplemental Figure S16

### Supplemental Figure S17

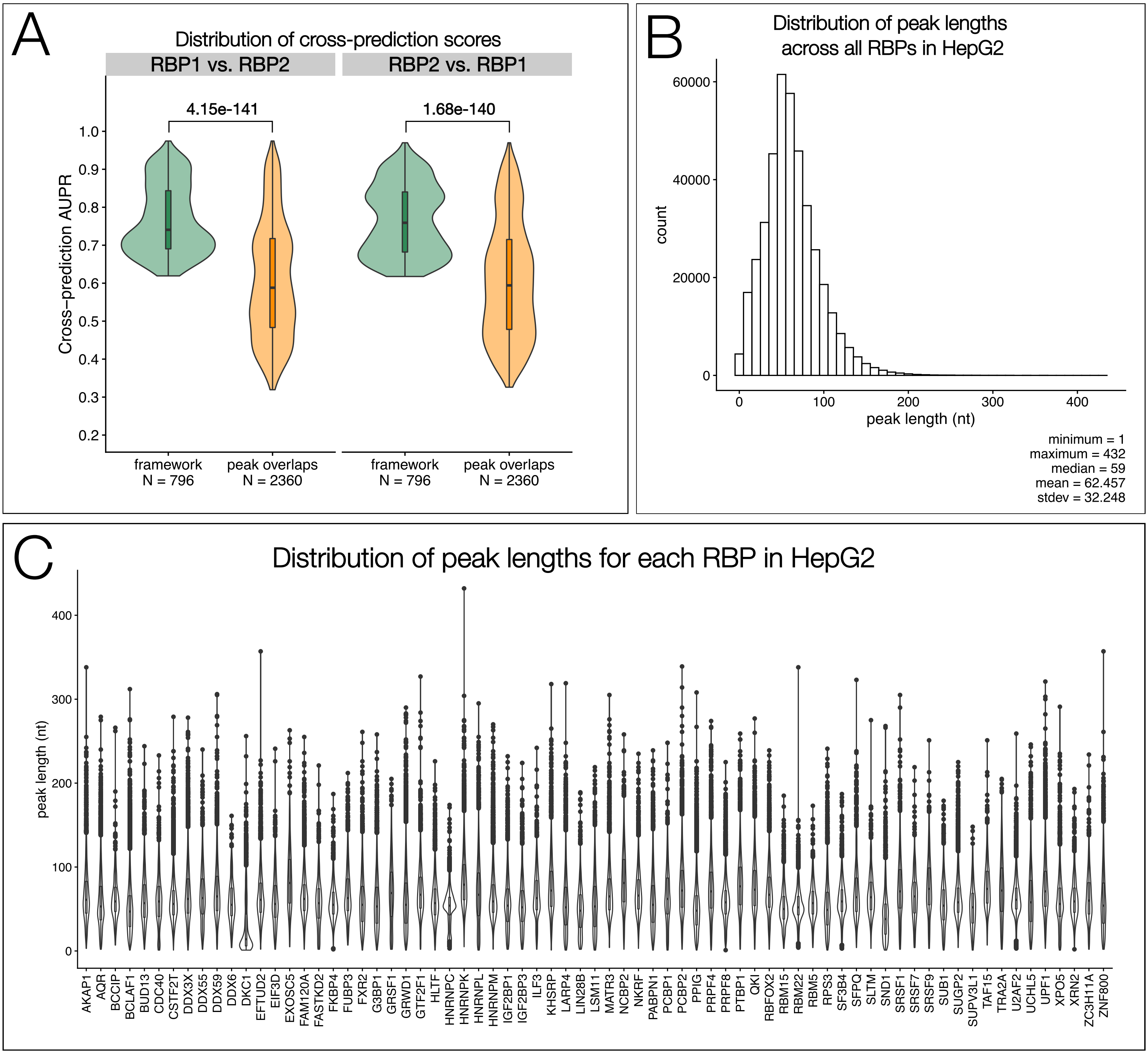

### Supplemental Figure S18

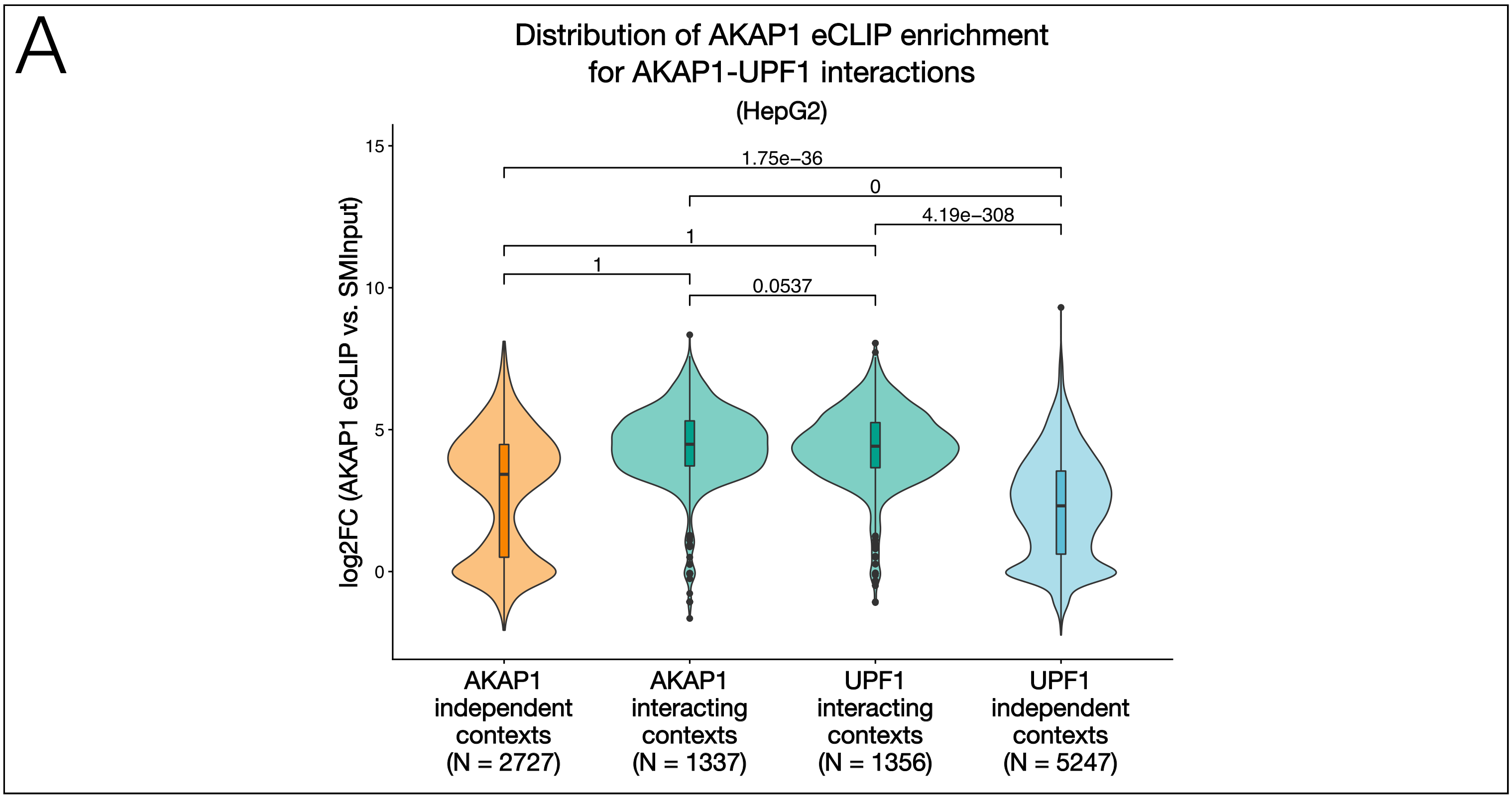

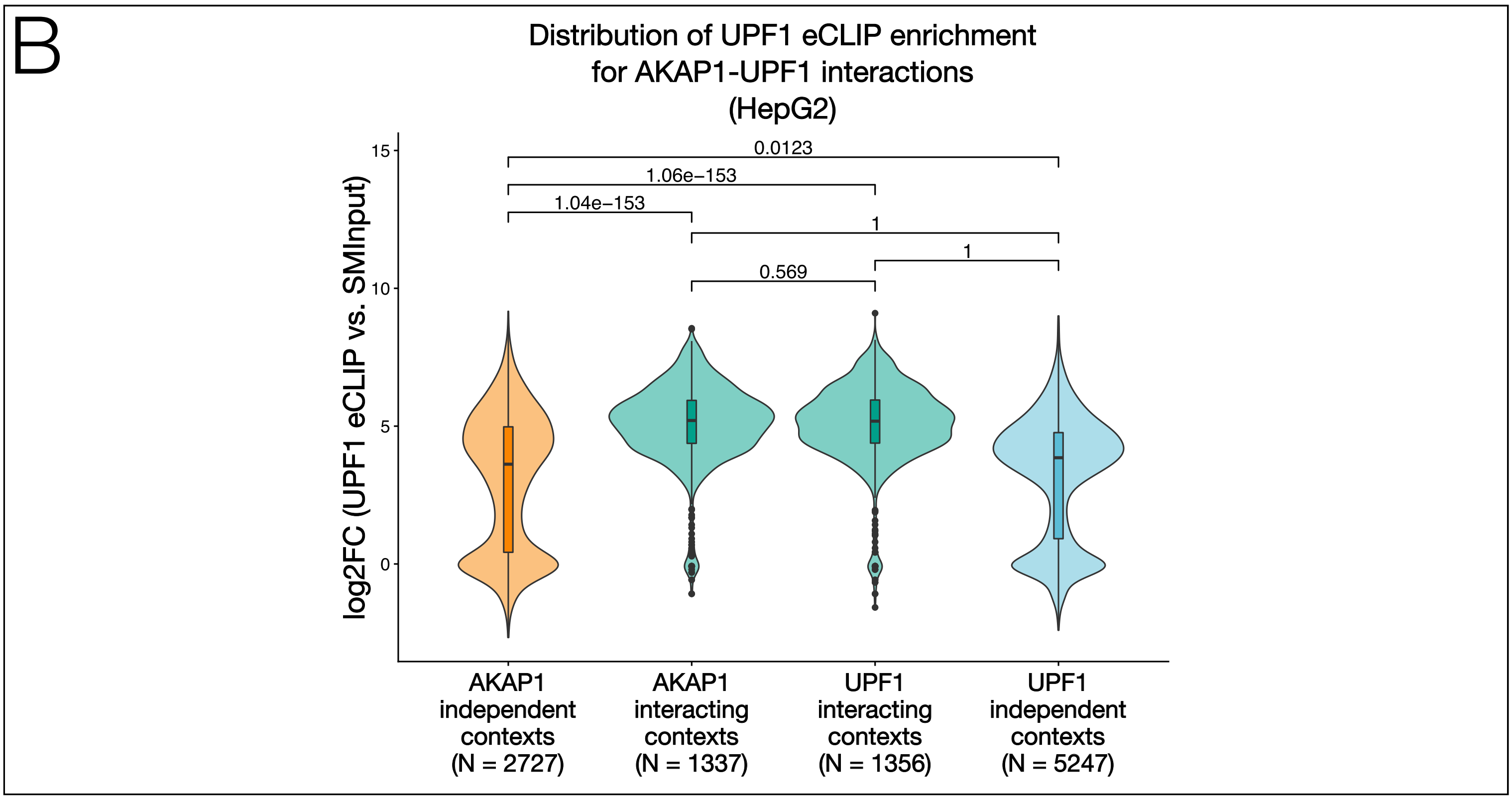
