## Supplemental Figure S3 for "A novel NLP-based method and algorithm to discover RNA-binding protein (RBP) motifs, contexts, binding preferences, and interactions"

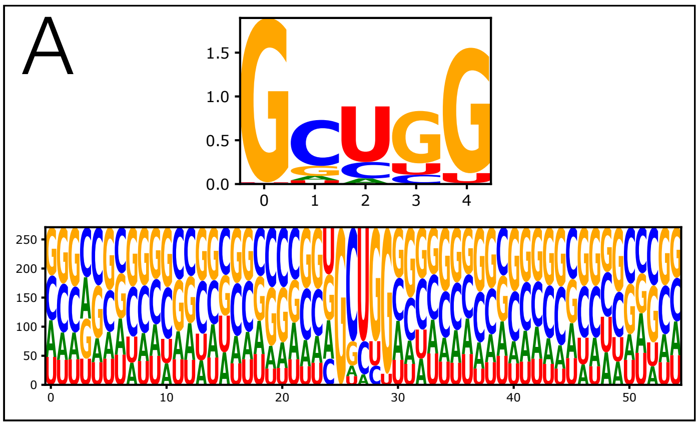


B

| Motif | Logo | P-value | E-value |
|---|---|---|---|
| 1-GGGSCCGGGS | | 4.2e-114 | 4.2e-113 |
| 2-GGCCCC | | 4.0e-058 | 4.0e-057 |
| 3-SSAGGCS | | 7.8e-045 | 7.8e-044 |
| 4-BCCCUGCC | | 1.8e-023 | 1.8e-022 |
| 5-GGGCC | | 2.2e-017 | 2.2e-016 |
| 6-GGGAGG | | 1.2e-015 | 1.2e-014 |
| 7-GGGCUGG | | 4.0e-009 | 4.0e-008 |
| 8-GGWGGG | | 1.4e-008 | 1.4e-007 |
| 9-CCUCCCC | | 3.2e-008 | 3.2e-007 |
| 10-GCCCC | | 1.5e-005 | 1.5e-004 |

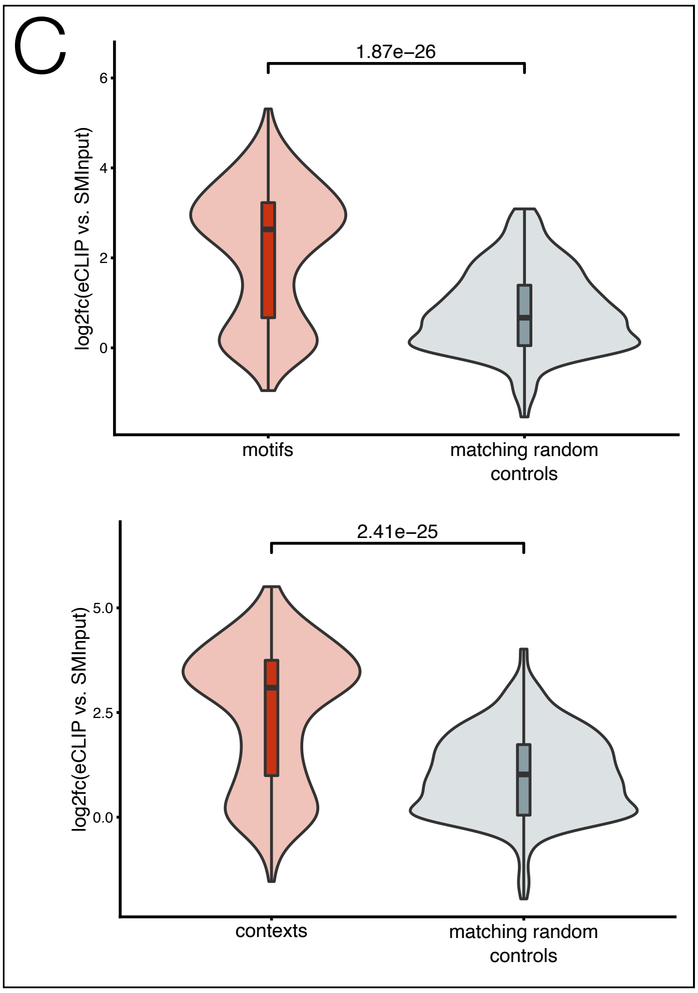


Supplemental Figure S3
