## Supplemental Figure S7 for "A novel NLP-based method and algorithm to discover RNA-binding protein (RBP) motifs, contexts, binding preferences, and interactions"

A

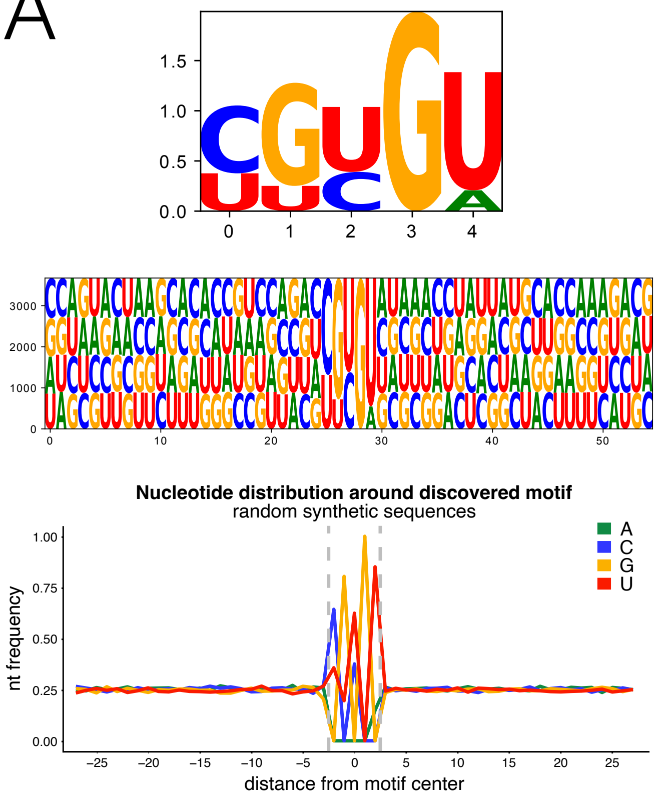


B

| Motif | Logo | P-value | E-value |
|---|---|---|---|
| 1-AUGAUAUUUG | 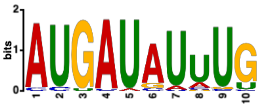 | 5.2e-048 | 5.2e-047 |
| 2-CACCAUGAGCA | 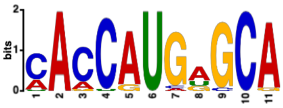 | 1.8e-037 | 1.8e-036 |
| 3-CCAGCACCCA | 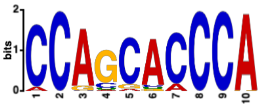 | 5.8e-036 | 5.8e-035 |
| 4-GAGUUCUUCGGG | 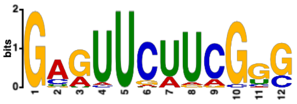 | 1.6e-027 | 1.6e-026 |
| 5-CGUAUCGCACAYUUG | 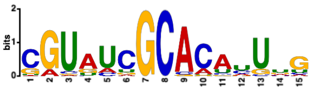 | 2.0e-025 | 2.0e-024 |
| 6-AGAUCCUGUAG | 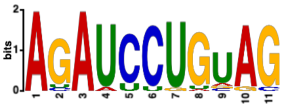 | 1.6e-024 | 1.6e-023 |
| 7-ACGAGWUGGCACAUG | 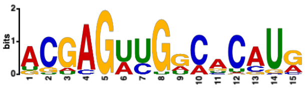 | 3.5e-024 | 3.5e-023 |
| 8-UUAUGCUCACCG | 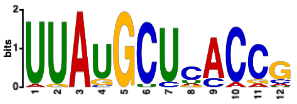 | 2.8e-022 | 2.8e-021 |
| 9-ACCGACCGUUG | 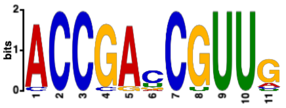 | 6.7e-021 | 6.7e-020 |
| 10-UGUCGGUU | 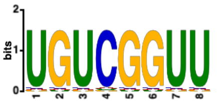 | 1.2e-017 | 1.2e-016 |

Supplemental Figure S7
